# The P-body component Edc3 is required for yeast proliferation in the phyllosphere

**DOI:** 10.1101/2025.09.14.675019

**Authors:** Fuka Sekioka, Kosuke Shiraishi, Miho Akagi, Akari Habata, Yumi Arima, Yasuyoshi Sakai, Hiroya Yurimoto

## Abstract

**Impact statement:** Edc3-dependent functions of P-bodies participate in the regulation of methanol-induced mRNAs and contribute to yeast adaptation to the phyllosphere

Processing bodies (P-bodies) are major cytosolic ribonucleoprotein granules involved in post-transcriptional regulation. Yeast has been an invaluable model for elucidating the functions of P-bodies under laboratory conditions. However, the physiological significance of P-bodies in natural environments remains unclear. Here, we demonstrate that the P-body component Edc3 is required for yeast proliferation in the phyllosphere, the aerial parts of plants. Deletion of *EDC3*, a gene critical for P-body formation, impaired proliferation of the methanol-utilizing yeast *Candida boidinii* on *Arabidopsis thaliana* leaves where the yeast assimilates methanol as the carbon source while adapting to changes in environmental conditions. *In vitro* experiments showed that Edc3 contributes to the spatiotemporal regulation of methanol-induced mRNAs (mimRNAs). These mimRNAs form cytosolic dot structures (termed mimRNA granules) that harbor multiple kinds of mimRNAs. In the *edc3Δ* strain, the formation of mimRNA granules was reduced along with a decrease in mimRNA abundance. Under oxidative stress, colocalization of P-bodies with mimRNA granules markedly increased and growth of the *edc3Δ* strain on methanol was suppressed, suggesting that Edc3 contributes to stress tolerance, possibly through enhanced sequestration of mimRNAs into P-bodies. Time-lapse microscopy revealed dynamic interactions between P-bodies and mimRNA granules with transient colocalization. Together, our findings indicate that Edc3-dependent functions of P-bodies participate in the regulation of mimRNAs and contribute to yeast adaptation to the phyllosphere.

## Introduction

Ribonucleoprotein (RNP) granules are membrane-less organelles formed through liquid–liquid phase separation (LLPS), concentrating RNAs and RNA-binding proteins into condensates within the cytosol or nucleus (Ripin & Parker, 2023). Among these, processing bodies (P-bodies) are recognized as major cytosolic RNP granules that function as hubs for post-transcriptional regulation, particularly under stress conditions (Decker & Parker, 2012; Kedersha & Anderson, 2009). P-bodies contain RNA-binding proteins and translationally repressed mRNAs, mediating mRNA storage and translational repression (Nissan et al., 2010). These granules are highly dynamic, undergo rapid and reversible assembly and disassembly, and frequently exchange components with other RNP granules (Buchan, 2024; Decker & Parker, 2012), including stress granules (SGs), which harbor translation initiation complexes (Swisher & Parker, 2010). P-bodies and SGs allow cells to protect key mRNAs from degradation and modulate their availability for translation in response to environmental stimuli through transient mRNA sequestration, offering an energy-efficient strategy compared to continuous cycles of mRNA degradation and *de novo* synthesis. Yeast has been an invaluable model for elucidating the functions of RNP granules under controlled laboratory conditions. Yet, the physiological relevance of RNP granules in complex and changing natural environments remains largely unexplored.

The phyllosphere, the aerial parts of plants, is one of the largest microbial habitats in nature (Ruinen, 1956). The total surface area of both sides of leaves on Earth is estimated at 6.4 × 10^8^ km^2^ and harbors a wide array of microbes, including yeasts (Morris & Kinkel, 2002). Despite its ecological richness, the phyllosphere represents a harsh environment characterized by oxidative stress, elevated temperatures and nutrient limitations (Beattie & Lindow, 1995; Lindow & Brandl, 2003; Vorholt, 2012). To survive under these conditions, phyllosphere microbes have evolved diverse adaptive mechanisms, including robust stress responses and metabolic flexibility.

Among these microbial residents, methylotrophic microbes have developed unique strategies to adapt to the phyllosphere environment. A key adaptation is their ability to utilize methanol, which is released as a product of pectin demethylesterification in plant cell walls during growth and is available for microbial use as a carbon source (Fall & Benson, 1996; Nemecek-Marshall et al., 1995; Yurimoto et al., 2021). Our previous study demonstrated that methanol concentrations on leaf surfaces oscillate diurnally, rising during the dark period (0.048–0.21%) and falling during the light period (0.014–0.015%) on growing plant leaves, while remaining elevated (≥0.8%) on wilting or dead plant leaves (Kawaguchi et al., 2011). Using the methylotrophic yeast *Candida boidinii*, which is frequently isolated from plant samples (Cadete et al., 2012; Mota et al., 2024), we showed that the yeast proliferates by using methanol as the carbon source on plant leaves and adapts to oscillating methanol availability by adjusting the transcription of methanol-induced genes in response to local methanol concentrations (Kawaguchi et al., 2011). Building on these findings, and drawing on recent findings that mRNAs encoding metabolic enzymes can be sequestered in P-bodies for temporal storage prior to translation (Lavut & Raveh, 2012; Lui et al., 2014; Standart & Weil, 2018), we hypothesized that *C. boidinii* adapts to environmental methanol availability not only transcriptionally, but also post-transcriptionally via P-body–mediated mRNA regulation. Yeast methylotrophy involves various enzymes localized in either peroxisomes or the cytosol (Yurimoto et al., 2011) (cf. Figure 2–figure supplement 1). In peroxisomes, alcohol oxidase (AOD) and dihydroxyacetone synthase (DAS) catalyze methanol oxidation and formaldehyde fixation, while catalase (CTA) and peroxisomal peroxidase (PMP20) detoxify reactive oxygen species (ROS). In the cytosol, formaldehyde dehydrogenase (FLD), *S*-formylglutathione hydrolase (FGH) and formate dehydrogenase (FDH) further metabolize formaldehyde and formate to CO_2_, yielding energy. Transcription of these enzymes is strongly induced by methanol, resulting in the accumulation of large quantities of methanol-induced mRNAs (mimRNAs) prior to protein synthesis. However, the intracellular fates of these mRNAs—whether they are stored, degraded or relocalized—remain unknown.

In this study, we investigated whether P-bodies contribute to yeast survival in the phyllosphere using a nature-mimicking experimental system with the methylotrophic yeast *C. boidinii* and plant *Arabidopsis thaliana*. We also examined the involvement of P-bodies in the intracellular dynamics of mimRNAs under methanol-utilizing conditions. Our findings provide new insight into the role of Edc3-dependent functions of P-bodies in the spatiotemporal regulation of mimRNAs and their contribution to yeast adaptation to the phyllosphere.

## Results

### The P-body component Edc3 is required for yeast proliferation in the phyllosphere

We first examined whether *C. boidinii* forms P-bodies in the phyllosphere using Edc3-Venus as a fluorescent marker because Edc3 localizes to P-bodies and is essential for their formation (Decker et al., 2007; Tishinov & Spang, 2021). The wild-type (WT) strain expressing Edc3-Venus (strain Edc3-V) was inoculated onto growing leaves of *A. thaliana* (2–3 weeks post-germination). Yeast cells were collected over time from plants grown under a daily light/dark cycle. Fluorescence microscopy detected cytosolic dots in yeast cells at 1, 3 and 7 days post-inoculation (dpi) (Figure 1A). Quantitative analysis showed that the dot formation of Edc3-Venus was similar under light and dark conditions and that the number of dots slightly increased over time (Figure 1B), demonstrating the constitutive presence of P-bodies in *C. boidinii* on leaves.

**Figure 1.**
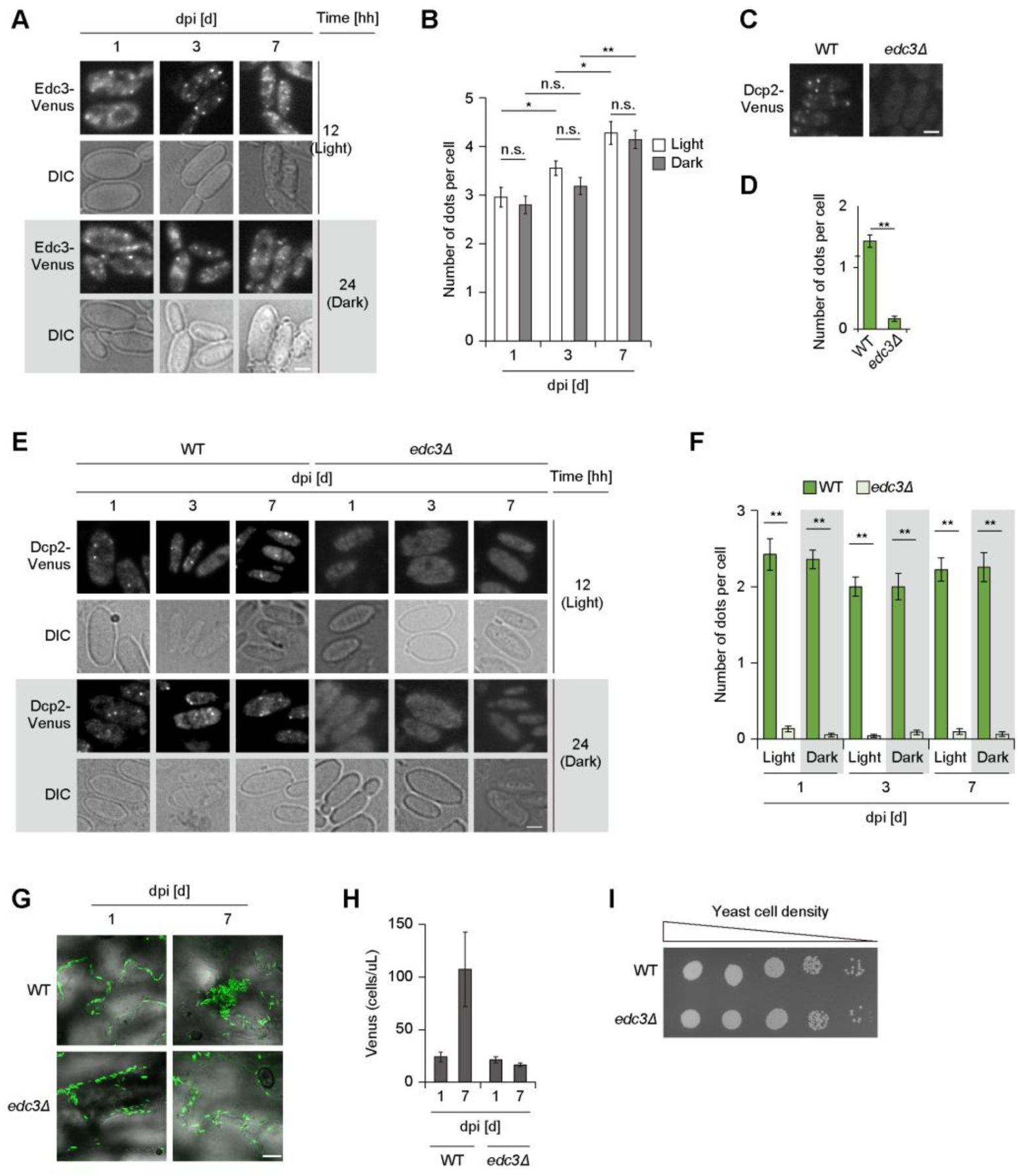
Formation and significance of yeast P-bodies on *A. thaliana* leaves. (A) P-body formation in the WT strain on growing *Arabidopsis* leaves. Edc3-Venus fusion protein was expressed under the control of the native *EDC3* promoter as a marker. The strain was spotted on leaves, collected from the leaves at the indicated day/time post-inoculation and observed by a fluorescence microscope. DIC: differential interference contrast. Bar, 2 µm. dpi, days post-inoculation. (B) Quantification of Edc3-Venus dot formation detected in (**A**). Cell count analysis was performed on fluorescence microscopy images (n = 30, n: number of cells analyzed; b = 3, b: technical replicates; total = 90 cells). The number of dots is presented as mean ± standard error (SE) of 90 cells. Asterisks indicate statistical significance: * *p* < 0.05; ** *p* < 0.01; n.s. means not significant. (C) P-body formation in the WT and *edc3Δ* strains on SM media. Dcp2-Venus fusion protein was expressed under the control of the native *DCP2* promoter as a marker. The strains grown on SM medium were collected for microscopic observation. Bar, 2 µm. (D) Quantification of Dcp2-Venus dot formation detected in (**C**). Cell count analysis was performed on fluorescence microscopy images (n = 30, n: number of cells analyzed; b = 3, b: technical replicates; total = 90 cells). The number of dots is presented as mean ± standard error (SE) of 90 cells. Asterisks indicate statistical significance: ** *p* < 0.01. (E) P-body formation in the WT and *edc3Δ* strains on growing *Arabidopsis* leaves. The strains expressing Dcp2-Venus were spotted on leaves, collected at the indicated day/time post-inoculation during the light and dark periods, and observed by fluorescence microscopy. Bar, 2 µm. (F) Quantification of Dcp2-Venus dot formation detected in (**E**). Cell count analysis was performed on fluorescence microscopy images (n = 30, n: number of cells analyzed; b = 3, b: technical replicates; total = 90 cells). The number of dots is presented as mean ± SE of 90 cells. Asterisks indicate statistical significance: ** *p* < 0.01. (G) Confocal microscope images of the Venus-labeled WT and *edc3Δ* strains on growing *Arabidopsis* leaves. Venus fluorescent protein was expressed under the control of the constitutive *ACT1* promoter. The strains were spotted on leaves and observed at the indicated days post-inoculation (dpi). Merged images of differential interference contrast (DIC) and Venus are depicted. Bar, 20 µm. (H) Quantification of *C. boidinii* cell populations on *A. thaliana* leaves by flowcytometry (FCM). The leaves onto which the strains were inoculated were retrieved at the indicated days post-inoculation (dpi) and were subjected to FCM for counting the cell population expressing Venus. Values are indicated as mean ± SE of four biological replicates. (I) Growth assay of cells cultured on methanol. Cells were grown to early log phase, adjusted to OD_610_=1, and 3 μL of 10-fold serial dilutions were dropped onto SM plates. Subsequently, the SM plates were incubated at 28°C. Cell growth was analyzed after 2 days. Growth of the wild-type (WT) and *edc3Δ* was compared.

To assess the physiological significance of the P-body component Edc3 in the phyllosphere, we disrupted the corresponding gene. The WT and *edc3Δ* strains expressing Venus fused to Dcp2, another P-body marker (Buchan et al., 2008), were examined under both synthetic medium and phyllosphere conditions. The suitability of Dcp2 as a P-body marker in *C. boidinii* was supported by the colocalization of 73% of Edc3-Venus dots with Dcp2-mCherry dots in cells grown in synthetic methanol (SM) medium (Figure 1–figure supplement 1). Deletion of *EDC3* led to a marked reduction in Dcp2-Venus dots on SM medium (Figure 1C, D). On *A. thaliana* leaves, Dcp2-Venus dots were also significantly reduced in the *edc3Δ* strain at 1, 3 and 7 dpi under both light and dark conditions (Figure 1E, F). These results indicate that Edc3 plays important roles in P-body formation in *C. boidinii* under both synthetic medium and phyllosphere conditions. Next, we investigated whether Edc3 is required for yeast proliferation in the phyllosphere. The WT and *edc3Δ* strains expressing Venus under the constitutive *ACT1* promoter were inoculated onto growing leaves and monitored for seven days. In the WT strain, confocal microscopy showed a progressive increase in fluorescent cells (Figure 1G), which was consistent with an approximately 4-fold increase in cell number as measured by flow cytometry (FCM) (Figure 1H). In contrast, no significant increase in cell number was observed in the *edc3Δ* strain (Figure 1G, H), although the mutant showed growth comparable to that of the WT strain in SM medium (Figure 1I). These results indicate that Edc3 is required for yeast proliferation on plant leaves.

Because the phyllosphere environment changes over the course of plant development (Kawaguchi et al., 2011; Shiraishi et al., 2015), we examined P-body formation in *C. boidinii* at different stages of plant growth: growing (2–3 weeks), matured (5–6 weeks), wilting (2 months) and dead (3 months post-germination). Edc3-Venus dots were consistently observed in yeast cells collected from leaves at all plant developmental stages (Figure 1–figure supplement 2A). We further asked whether SGs form under these conditions using the *C. boidinii* WT strain expressing Pbp1-Venus, a marker for SGs (Shiraishi et al., 2018) (strain Pbp1-V). No cytosolic dots were detected in strain Pbp1-V on leaves at any developmental stage under the tested conditions (Figure 1–figure supplement 2B). These observations demonstrate that only P-bodies and not SGs are formed in *C. boidinii* on leaf surfaces.

### P-bodies partially but continuously colocalize with “methanol-induced mRNA granules (mimRNA granules)”

We hypothesized that P-bodies contribute to the post-transcriptional regulation of genes encoding enzymes for methanol metabolism (shown in Figure 2–figure supplement 1), supporting yeast proliferation on leaves. To test this physiological role of P-bodies in the phyllosphere, we investigated the intracellular dynamics of P-bodies and mimRNAs. We selected *DAS1* mRNA as a representative mimRNA because *DAS1* is one of the most strongly induced genes upon methanol induction (Yurimoto et al., 2000), making its transcripts suitable for visualization. *DAS1* mRNA was visualized by a Venus-tagged U1A-based RNA system (Takizawa & Vale, 2000) (see Figure 2–figure supplement 2A for methodological overview) and P-bodies by Cerulean-tagged Edc3 in the WT strain (strain Edc3-C/DAS1m-V). The validity of the U1A system in *C. boidinii* was confirmed by colocalization between U1A-based signals and fluorescence *in situ* hybridization (FISH) (Long et al., 1995) signals for *DAS1* mRNA (Figure 2–figure supplement 2B, C), as well as a complementary growth assay (Figure 2–figure supplement 2D). Strain Edc3-C/DAS1m-V was then inoculated onto growing *Arabidopsis* leaves and confocal microscopy was used to detect cytosolic dots of both Edc3-Cerulean and Venus-tagged *DAS1* mRNA, some of which colocalized with each other (Figure 2A). Fluorescence microscopy of yeast cells collected from leaves provided clearer images and confirmed dot formation under both light and dark conditions, with 13% of Edc3-Cerulean and Venus-tagged *DAS1* mRNA dots showing colocalization (Figure 2B, C). Partial colocalization was also detected in yeast cells on wilting leaves (Figure 2–figure supplement 3A, B).

**Figure 2.**
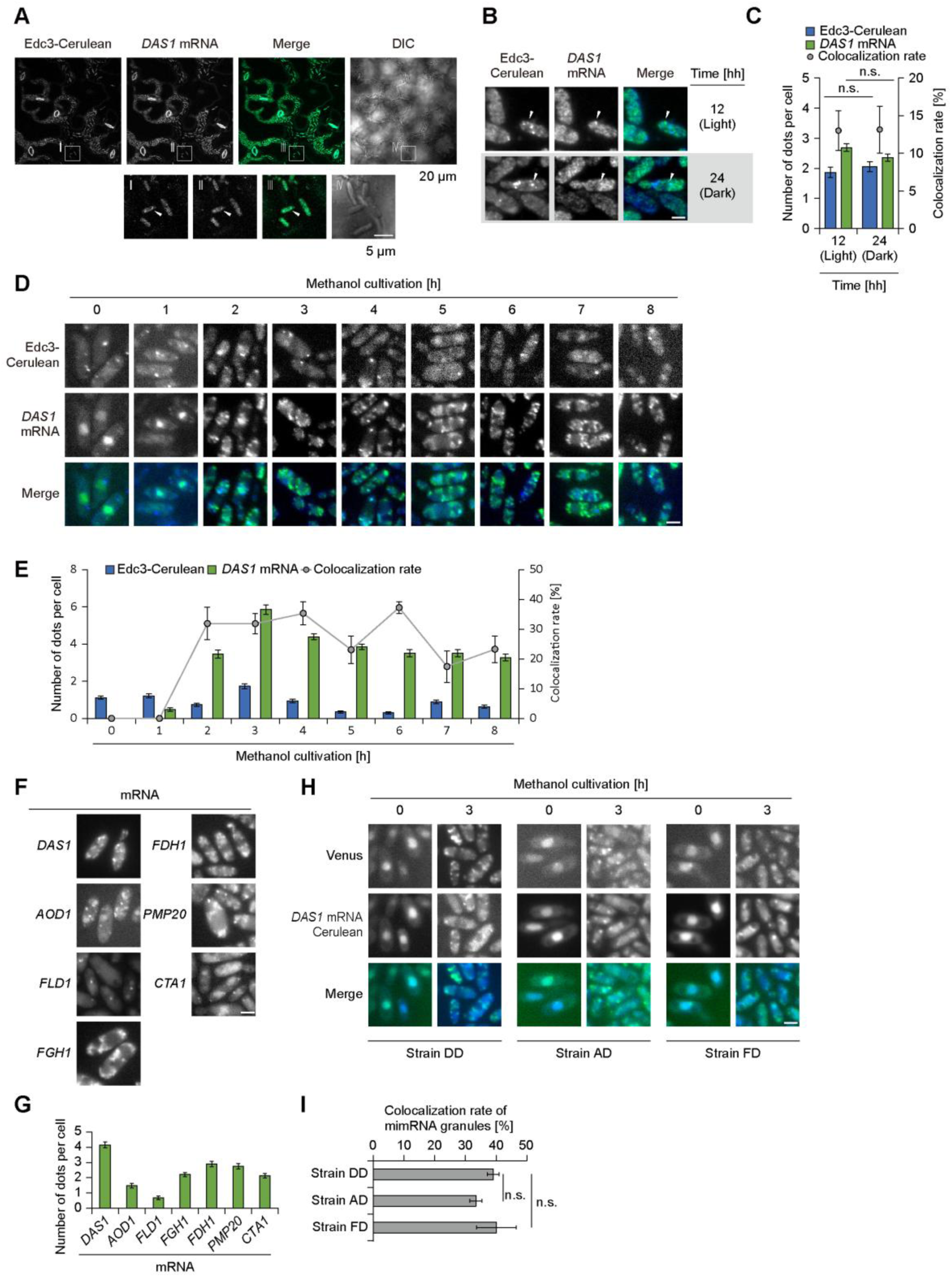
Intracellular dynamics of P-bodies and mimRNAs on methanol growth conditions. (A) Confocal microscopy images of the WT strain expressing Edc3-Cerulean and Venus-tagged *DAS1* mRNA (strain Edc3-C/DAS1m-V) on growing *Arabidopsis* leaves. The strain was spotted on the leaves and observed 4 hours after inoculation. Squares I, II, III and VI were enlarged. Merged images were generated by combining Venus and Cerulean fluorescence images. White arrows indicate colocalization of Edc3-Cerulean and Venus-tagged *DAS1* mRNA dots. The lengths of the bars are indicated in the images. DIC: differential interference contrast. (B) Fluorescence microscopy images of strain Edc3-C/DAS1m-V on growing *Arabidopsis* leaves. The strain, spotted on leaves, was collected for observation from the leaf surface 4 hours after inoculation during both the light and dark periods. Merged images were generated by combining Venus and Cerulean fluorescence images. White arrows indicate colocalization of Edc3-Cerulean and Venus-tagged *DAS1* mRNA dots. Bar, 2 µm. (C) Quantification of dot formation of Edc3-Cerulean and Venus-tagged *DAS1* mRNA and their colocalization detected in (**B**). Cell count analysis was performed on fluorescence microscopy images (n = 30, n: number of cells analyzed; b = 3, b: technical replicates; total = 90 cells). The number of dots is presented as the mean ± SE of 90 cells, and colocalization rates are shown as mean ± SE from three technical replicates (n = 30 cells per replicate). n.s. means not statistically significant. (D) Time-course analysis of the intracellular localization of P-bodies and mimRNAs in strain Edc3-C/DAS1m-V during methanol induction. The strain grown on an SD medium was transferred to SM medium and collected at the indicated time points for fluorescence microscopy. Merged images were generated by combining Venus and Celurean fluorescence images. Bar, 2 µm. (E) Quantification of dot formation of Edc3-Cerulean and Venus-tagged *DAS1* mRNA and their colocalization detected in (**D**). Cell count analysis was performed on fluorescence microscopy images (n = 30, n: number of cells analyzed; b = 3, b: technical replicates; total = 90 cells). The number of dots is presented as mean ± SE of 90 cells and colocalization rates are shown as mean ± SE from three technical replicates (n = 30 cells per replicate). (F) Fluorescence microscopy images of the strains visualizing *DAS1*, *AOD1*, *FLD1*, *FGH1*, *FDH1*, *PMP20* and *CTA1* mRNAs. The strains grown on an SD medium were transferred to SM medium and incubated for 3 hours before fluorescence microscopy. Bar, 2 µm. (G) Quantification of mimRNA granule formation detected in (**F**). Cell count analysis was performed on fluorescence microscopy images (n = 30, n: number of cells analyzed; b = 3, b: technical replicates; total = 90 cells). The number of dots is presented as mean ± SE of 90 cells. (H) Intracellular localization of multiple mimRNAs during methanol induction. In strain DD, *DAS1* mRNA was visualized by the Venus-tagged U1A-based RNA system and Cerulean-tagged MS2-based RNA system. In strain AD and strain FD, *AOD1* or *FLD1* mRNA was visualized by Venus-tagged U1A-based RNA system and *DAS1* mRNA by Cerulean-tagged MS2-based RNA system. Merged images were generated by combining Venus and Cerulean fluorescence images. Bar, 2 µm. (I) Quantification of the colocalization of the multiple mimRNA granules detected in (**H**). Cell count analysis was performed on fluorescence microscopy images (n = 30, n: number of cells analyzed; b = 3, b: technical replicates; total = 90 cells). The number of dots is presented as mean ± SE of 90 cells. n.s. means not statistically significant.

Since *DAS1* mRNA dots were also observed independently of P-bodies, we next conducted *in vitro* experiments to test whether Edc3-Cerulean and Venus-tagged *DAS1* mRNA dots are distinct. Strain Edc3-C/DAS1m-V grown on synthetic glucose (SD) medium was transferred to SM medium to induce the expression of genes involved in methanol metabolism. Time-course microscopy showed that Edc3-Cerulean dots were present throughout, with moderate changes in number (0.3 ∼ 1.7 dots per cell) (Figure 2D, E). By contrast, Venus-tagged *DAS1* mRNA dots appeared only under methanol growth conditions. The dot number increased sharply at 2 h and remained at a comparable level with modest variations until 8 h (3.5 ∼ 5.9 dots per cell) (Figure 2D, E). During this period, approximately 30% of Edc3-Cerulean dots colocalized with Venus-tagged *DAS1* mRNA dots (Figure 2E). Confocal imaging also confirmed colocalization of Edc3-Cerulean and Venus-tagged *DAS1* mRNA dots (Figure 2–figure supplement 4). A comparable colocalization rate (approximately 30%) was observed at 3 h after the shift to SM medium when Dcp2 was used as a P-body marker (Figure 2–figure supplement 5). These results indicate that Venus-tagged *DAS1* mRNA dots and P-bodies are distinct structures, although a subset of these structures colocalizes with each other during growth on methanol.

To characterize the mimRNA dots further, we visualized six additional mimRNAs: *AOD1*, *FLD1*, *FGH1*, *FDH1*, *PMP20* and *CTA1* (cf. Figure 2–figure supplement 1). We confirmed the functionality of the U1A system for mimRNAs encoding cytosolic enzymes by a complementary growth assay, using *FLD1* mRNA (Figure 2–figure supplement 2D). Fluorescence microscopy showed dot formation for all tested mimRNAs under methanol growth conditions, although dot numbers varied (Figure 2F, G). To assess the intracellular dynamics of multiple mimRNAs, we employed the MS2-based RNA system, another method for visualizing mRNAs (Bertrand et al., 1998b).

Approximately 40% of *DAS1* mRNA dots visualized by the MS2 system overlapped with those detected by the U1A system in strain DD during growth on methanol (Figure 2H, I). We then visualized *AOD1* mRNA or *FLD1* mRNA with the Venus-tagged U1A-based system in strains expressing Cerulean-tagged *DAS1* mRNA by the MS2 system (strain AD and strain FD, respectively). In both strains, partial colocalization was observed (Figure 2H), with a comparable rate to strain DD (Figure 2I), suggesting that multiple mimRNAs share granules at least in part. Together, we defined methanol-induced cytosolic dots enriched with multiple mimRNAs as “mimRNA granules”.

### The P-body component Edc3 contributes to the post-transcriptional regulation of mimRNAs

To further investigate whether Edc3 contributes to the post-transcriptional regulation of mimRNAs, we compared *DAS1* mRNA dot formation in the WT and *edc3Δ* strains (strain DAS1m-V and strain edc3Δ-DAS1m-V, respectively). Fluorescence microscopy detected Venus-tagged *DAS1* mRNA dots in both strains during growth on methanol (Figure 3A). However, quantitative analysis revealed significantly fewer dots in strain edc3Δ-DAS1m-V compared to strain DAS1m-V (Figure 3B), suggesting that Edc3 contributes to the formation and/or maintenance of mimRNA granules. To assess the impact of Edc3 deletion on transcript abundance, we measured the *DAS1* transcript level by qRT-PCR. *DAS1* transcript levels were lower in the *edc3Δ* strain than in the WT (Figure 3C). Similar results were obtained for *AOD1* mRNA (Figure 3–figure supplement 1A). These results, in conjunction with previous suggestions (Wang et al., 2018), led us to infer that Edc3 contributes to maintaining mimRNA levels, possibly through sequestration of mimRNAs in P-bodies. Protein levels of DAS and AOD (Figure 3–figure supplement 1B, C) and cell growth on SM medium (Figure 1I) in the *edc3Δ* strain were comparable to those of the WT strain under the tested conditions.

**Figure 3.**
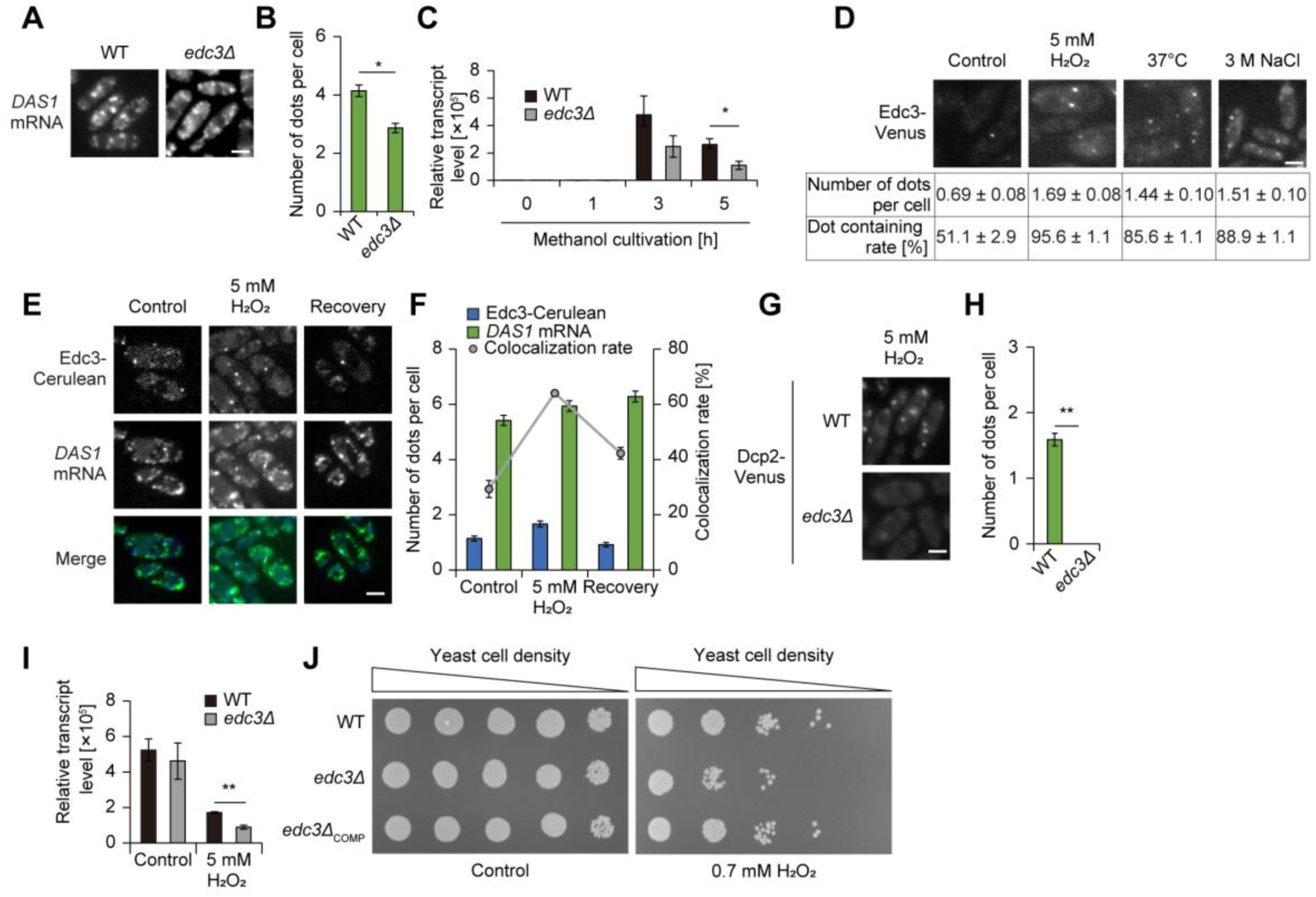
Role of P-bodies on post-transcriptional regulation of mimRNAs. (A) Dot formation of Venus-tagged *DAS1* mRNA in the WT and *edc3Δ* strains on SM medium. The strains grown on an SD medium were transferred to SM medium and incubated for 3 hours before fluorescence microscopy. Bar, 2 µm. (B) Quantification of Venus-tagged *DAS1* mRNA dot formation detected in (**A**). Cell count analysis was performed on fluorescence microscopy images (n = 30, n: number of cells analyzed; b = 3, b: technical replicates; total = 90 cells). The number of dots is presented as mean ± SE of 90 cells. Asterisks indicate statistical significance: * *p* < 0.05. (C) Relative transcript levels of *DAS1* mRNA in the WT and *edc3Δ* strains on SM medium. Values are indicated as mean ± SE of three biological replicates. Statistical significance between the WT and *edc3Δ* strains were assessed. Asterisks indicate statistical significance: * *p* < 0.05. (D) P-body formation under stress conditions. Strain Edc3-V was exposed to oxidative stress with 5 mM H_2_O_2_, heat shock at 37 degrees or salt stress with 3 M NaCl for 1 hour before microscopic observation. Cell count analysis was performed on fluorescence microscopy images (n = 30, n: number of cells analyzed; b = 3, b: technical replicates; total = 90 cells). The number of dots is presented as mean ± SE of 90 cells, and the proportion of cells containing at least one dot (dot containing rate) are shown as mean ± SE from three technical replicates (n = 30 cells per replicate). Bar, 2µm. (E) Fluorescence microscopy images of strain Edc3-C/DAS1m-V during oxidative stress and recovery from the stress. The strain incubated on an SM medium was treated with 5 mM H_2_O_2_ for 1 hour and then transferred to an SM medium. Merged images were generated by combining Venus and Cerulean fluorescence images. Bar, 2 µm. (F) Quantification of dot formation of Edc3-Cerulean and Venus-tagged *DAS1* mRNA and their colocalization detected in (**E**). Cell count analysis was performed on fluorescence microscopy images (n = 30, n: number of cells analyzed; b = 3, b: technical replicates; total = 90 cells). The number of dots is presented as mean ± SE of 90 cells and colocalization rates are shown as mean ± SE from three technical replicates (n = 30 cells per replicate). (G) P-body formation in the WT and *edc3Δ* strains expressing Dcp2-Venus under oxidative stress. The strains grown on SM medium were treated with 5 mM H_2_O_2_ for 1 hour and collected for microscopic observation. (H) Quantification of Dcp2-Venus dot formation detected in (**G**). Cell count analysis was performed on fluorescence microscopy images (n = 30, n: number of cells analyzed; b = 3, b: technical replicates; total = 90 cells). The number of dots is presented as mean ± SE of 90 cells. Asterisks indicate statistical significance: ** *p* < 0.01. (I) Relative transcript level of *DAS1* mRNA in the WT and *edc3Δ* strains on SM medium under oxidative stress. The strain was incubated on SM medium for 3 hours and then treated with 5 mM H₂O₂ for 1 hour. Transcript levels of all genes were first normalized to *ACT1* and then expression levels were calculated relative to samples collected at 0 h of methanol cultivation. In the graph, the sample collected at 3 h is labeled as “Control” for comparison. Values are indicated as mean ± SE of three biological replicates. Statistical significance between the WT and *edc3Δ* strains was assessed. Asterisks indicate statistical significance: ** *p* < 0.01. (J) Growth assay of cells cultured under oxidative stress conditions. Cells were grown to early log phase, adjusted to OD_610_=1, and 3 μL of tenfold serial dilutions were dropped onto SM plates with or without 0.7 mM H_2_O_2_. Subsequently, the SM plates were incubated at 28°C. Cell growth was analyzed after 2 days. Growth of the wild-type (WT), *edc3Δ,* and *edc3Δ* expressing Edc3-Venus (*edc3Δ*_COMP_) under the native promoter was compared.

The phyllosphere is a harsh environment where yeast cells experience various stresses, including oxidative stress and elevated temperatures, many of which are known to enhance P-body formation (Teixeira & Parker, 2007). We therefore examined whether these stress conditions affect P-body formation and the recruitment of mimRNAs to P-bodies. To test this, we first examined P-body formation in *C. boidinii* exposed to various stresses. Treatment with 5 mM H_2_O_2_ (oxidative stress), 37°C (heat shock) or 3 M NaCl (salt stress) increased both the number of Edc3-Venus dots and the proportion of cells containing at least one dot, indicating enhanced P-body formation (Figure 3D). The strongest effect was observed under oxidative stress, with an average of 1.69 dots per cell and 95.6% cells containing at least one dot.

Subsequently, we examined the intracellular dynamics of P-bodies and mimRNAs under oxidative stress during growth on methanol. Strain Edc3-C/DAS1m-V was exposed to 5 mM H_2_O_2_ for 1 hour and analyzed by fluorescence microscopy. Following oxidative stress, the colocalization rate of Edc3-Cerulean dots with Venus-tagged *DAS1* mRNA dots increased from approximately 30% to 60% (Figure 3E, F), indicating that P-bodies sequester mimRNAs more efficiently. Heat shock evoked similar increases, while salt stress did not result in a significant change (Figure 3–figure supplement 2A, B). We also monitored the intracellular dynamics of P-bodies and mimRNAs during recovery from oxidative stress. Strain Edc3-C/DAS1m-V, after being exposed to 5 mM H_2_O_2_ for 1 hour, was transferred to SM medium and observed after 1 hour. The colocalization rate declined from approximately 60% to 40%, suggesting that mimRNA sequestration into P-bodies is reversible (Figure 3E, F).

We next evaluated the role of Edc3 in yeast survival under oxidative stress. Deletion of *EDC3* markedly reduced P-body formation, as indicated by a reduction in the number of Dcp2-Venus dots (Figure 3G, H), and decreased *DAS1* transcript levels (Figure 3I). These results were consistent with our observations under normal conditions (cf. Figure 1C, D, 3C). We then performed a spot assay using SM agar plates with or without H_2_O_2_. Growth of the *edc3Δ* strain was impaired under oxidative stress, whereas no clear difference was observed under normal conditions (Figure 3J). The expression of Edc3-Venus restored stress tolerance. These results suggest that Edc3 contributes to yeast tolerance to oxidative stress.

### mimRNA granule formation is conserved among methylotrophic yeasts and is regulated in accordance with the transcript levels

We next investigated how mimRNA granule formation is regulated. Fluorescence microscopy showed that Venus fluorescence in strain DAS1m-V remained localized to the DAPI-stained nucleus until 2–2.5 h after the medium shift, whereas cytosolic dot structures began to appear around 2 h after the shift (Figure 4A, B). Quantitative analysis showed that the number of Venus-tagged *DAS1* mRNA dots started to increase at 2 h, peaked for the next 1 to 2 hours, and then declined moderately (Figure 4C). A similar mountain-shaped curve pattern was observed at the *DAS1* transcript level (Figure 4D). DAS protein levels began to increase at 2.5 h (Figure 4E), 0.5 h after the rise in both Venus-tagged *DAS1* mRNA dot number and *DAS1* mRNA level. Similar patterns were obtained for the other tested mimRNAs (Figure 4–figure supplement 1A-M), suggesting that mimRNA granule formation is induced by methanol and is regulated in accordance with the transcript levels of methanol-induced genes. To further test the correlation between granule formation and transcript levels of mimRNAs, strain DAS1m-V grown on SD medium was transferred to SM medium supplemented with either 0.005% methanol (low-methanol) or 0.5% methanol (high-methanol), and harvested 3 h after the medium shift for microscopy and qRT-PCR analyses. Under low-methanol conditions, both the number of Venus-tagged *DAS1* mRNA dots and *DAS1* transcript levels were lower than those observed under high-methanol conditions (Figure 4–figure supplement 2A-C), supporting our idea that mimRNA granule formation correlates with its transcript level. In addition to *C. boidinii*, we observed dot structures of methanol-induced *AOX1* mRNA in *Komagataella phaffii*, another methylotrophic yeast, following a medium shift from SD to SM (Figure 4–figure supplement 3). This finding suggests that the regulatory mechanism of mimRNA granule formation is conserved among methylotrophic yeasts.

**Figure 4.**
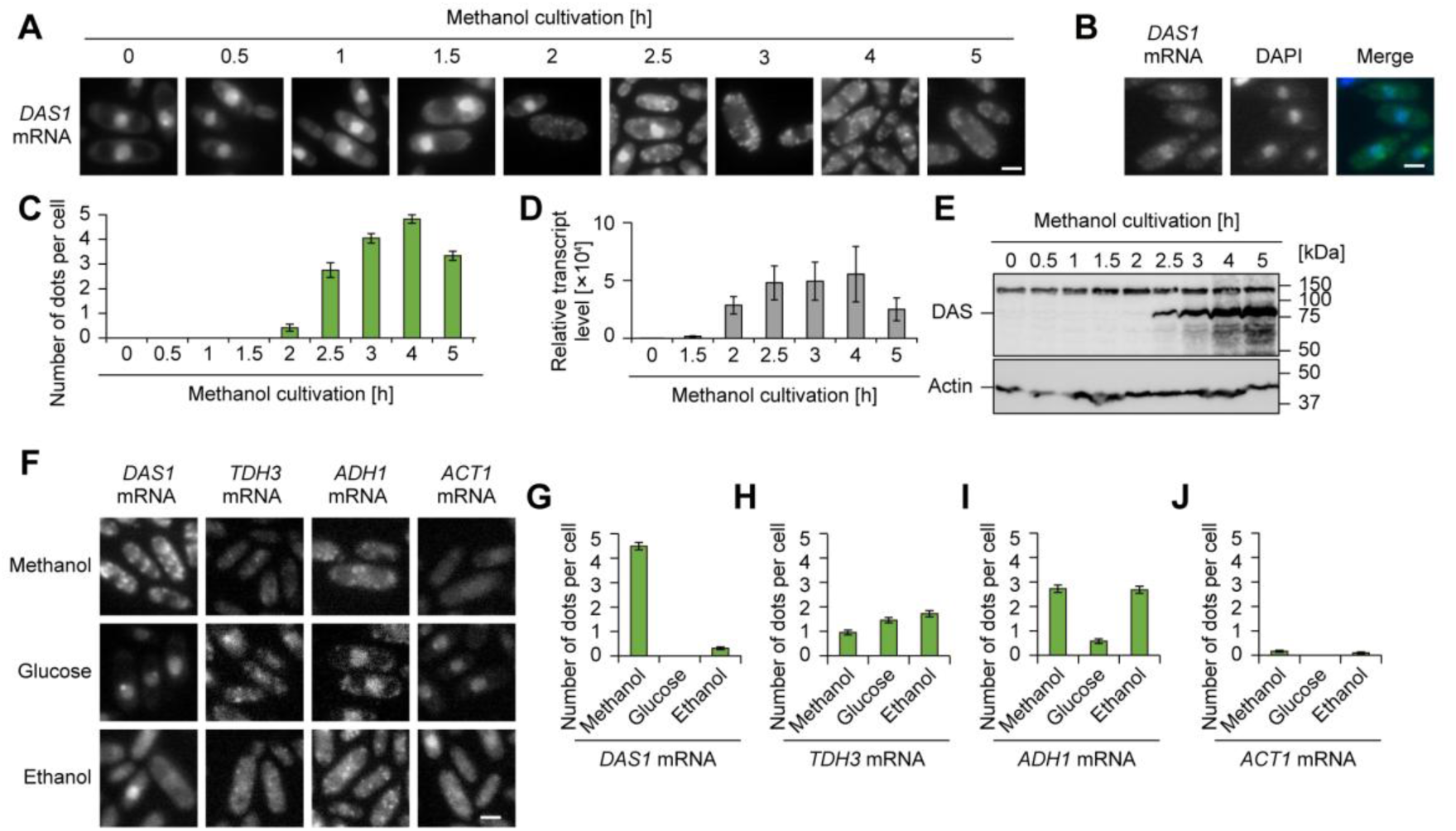
Regulation of mimRNA granule formation. (A) Time-course analysis of the intracellular localization of Venus-tagged *DAS1* mRNA during methanol induction. The strain grown on an SD medium was transferred to SM medium and incubated for up to 5 hours and collected at indicated time points. Bar, 2 µm. (B) Nuclear localization of Venus-tagged *DAS1* mRNA. The strain grown on an SD medium was collected for fluorescence microscopy. A merged image was generated by combining Venus and DAPI fluorescence images. Bar, 2 µm. (C) Quantification of dot formation of Venus-tagged *DAS1* mRNA detected in (**A**). Cell count analysis was performed on fluorescence microscopy images (n = 30, n: number of cells analyzed; b = 3, b: technical replicates; total = 90 cells). The number of dots is presented as mean ± SE of 90 cells. (D) Relative transcript level of *DAS1* mRNA in the WT strain during methanol induction. Values are indicated as the mean ± SE of three biological replicates. (E) Immunoblot analysis of DAS in the WT strain during methanol induction. The expected molecular weight of DAS is 78.2 kDa. (F) Fluorescence microscopy images of the strains visualizing *DAS1*, *TDH3*, *ADH1* and *ACT1* mRNAs during cultivation on various carbon sources. The strains grown on an SD medium were transferred to either SM, SD or SE medium and incubated for 3 hours. Bar, 2 µm. (**G-J**) Quantification of mRNA dot formation detected in (**F**). Cell count analysis was performed on fluorescence microscopy images (n = 30, n: number of cells analyzed; b = 3, b: technical replicates; total = 90 cells). The number of dots is presented as mean ± SE of 90 cells. (**G**) *DAS1* mRNA, (**H**) *TDH3* mRNA, (**I**) *ADH1* mRNA and (**J**) *ACT1* mRNA.

To compare the regulation of dot formation between mimRNAs and non-methanol-induced mRNAs, we visualized *TDH3*, *ADH1* and *ACT1* mRNAs in *C. boidinii* using the U1A system. These mRNAs encode the glycolytic enzyme glyceraldehyde-3-phosphate dehydrogenase (GAPDH), alcohol dehydrogenase involved in ethanol assimilation, and actin, respectively (Sasano et al., 2008; Wylin et al., 1998; Yurimoto et al., 2004). In contrast to *DAS1* mRNA (which showed a dramatic increase in dot formation under methanol growth conditions), Venus-tagged *TDH3* and *ADH1* mRNAs formed dots under all tested conditions. *ACT1* mRNA exhibited only a few dots during growth on methanol and ethanol (Figure 4F-J). Because *TDH3* mRNA formed dots under all tested conditions, we examined its dot formation during methanol induction.

Unlike methanol-induced *DAS1* mRNA, Venus-tagged non-methanol-induced *TDH3* mRNA formed cytosolic dots even before methanol induction (Figure 4–figure supplement 4A), and the number of dots decreased after the medium shift to SM medium (Figure 4–figure supplement 4B). The *TDH3* transcript levels decreased after the medium shift (Figure 4–figure supplement 4C) and GAPDH protein was detected under all conditions (Figure 4–figure supplement 4D), suggesting that *TDH3* mRNA dots are not induced by methanol. We also examined colocalization of Edc3-Cerulean with Venus-tagged *TDH3* mRNA dots during growth on methanol using strain Edc3-C/TDH3m-V. Under normal conditions, the colocalization rate was 7%, which increased to 15% after 1 h of oxidative stress treatment (Figure 4–figure supplement 5A, B).

### Spatiotemporal properties and dynamics of mimRNA granules

Finally, we sought to characterize the properties of mimRNA granules. To this end, we examined their dynamics alongside P-bodies in live cells during cultivation on methanol. Time-lapse microscopy revealed that *DAS1* mRNA dots underwent rapid assembly and disassembly, similar to P-bodies (Figure 5A, Figure 5—video 1). For example, a new Venus-tagged *DAS1* mRNA dot appeared at 56 seconds in the video, which colocalized with an Edc3-Cerulean dot from 1 min 24 s to 2 min 48 s, and disappeared at 3 min 16 s, while the Edc3-Cerulean dots remained. These observations indicate that P-bodies and mimRNA granules transiently colocalize with each other. Additionally, at 7 min 56 s, a Venus-tagged *DAS1* mRNA dot appeared without colocalizing with a P-body; this granule remained at the same site at 8 min 24 s but disappeared at 8 min 52 s, suggesting possible relocalization of the mimRNA.

**Figure 5.**
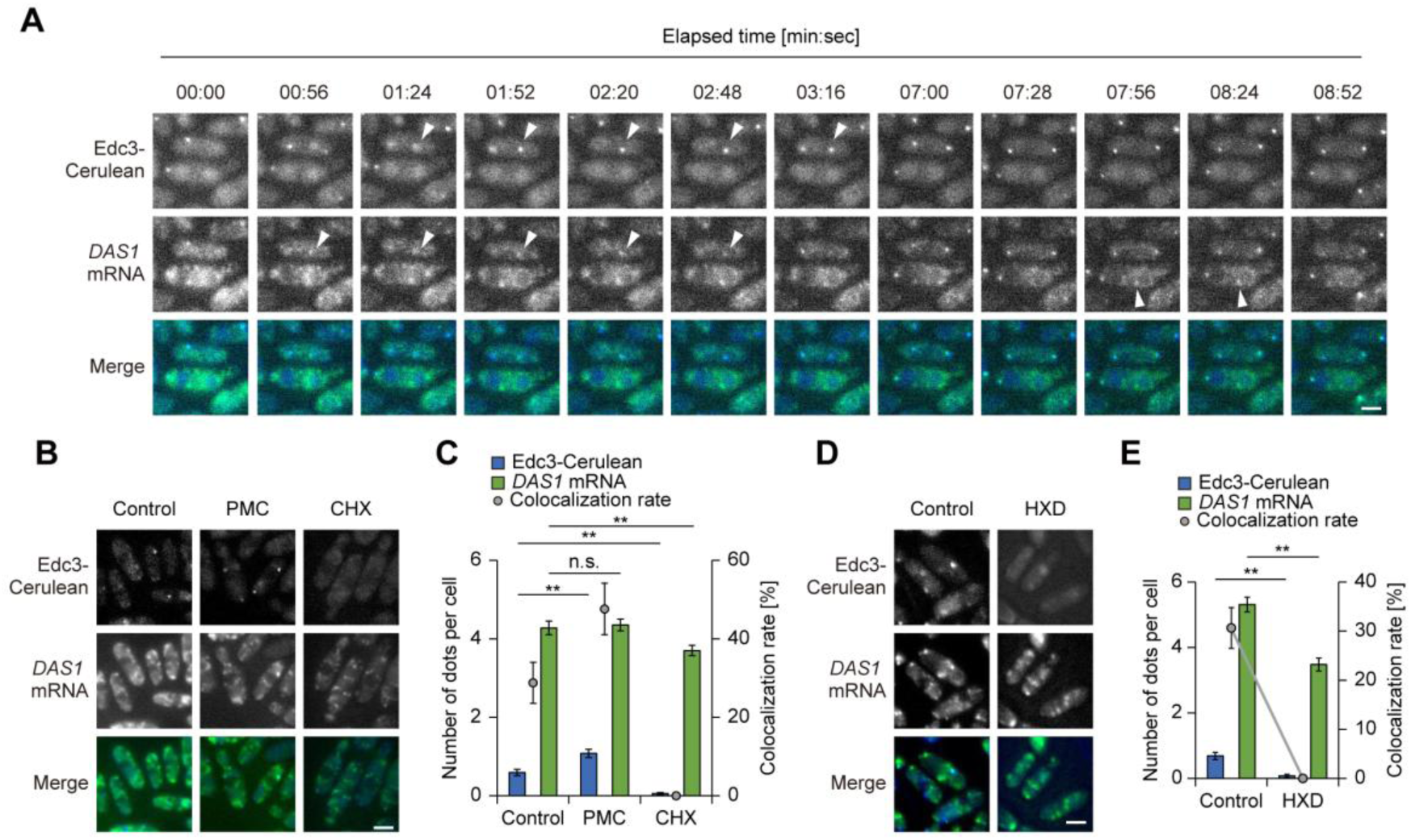
Properties of mimRNA granules. (A) Time-lapse images of strain Edc3-C/DAS1m-V under methanol-grown conditions. Twelve images extracted at the indicated time points are shown as a video (see also Figure 5–video 1). Merged images were generated by combining Venus and Cerulean fluorescence images. White arrows indicate the dynamic properties of Edc3-Cerulean and Venus-tagged *DAS1* mRNA dots. A new Venus-tagged *DAS1* mRNA dot appeared at 56 seconds in the video, which colocalized with an Edc3-Cerulean dot from 1 min 24 s to 2 min 48 s, and disappeared at 3 min 16 s, while the Edc3-Cerulean dots remained. At 7 min 56 s, a Venus-tagged *DAS1* mRNA dot appeared without colocalizing with a P-body; this granule remained at the same site at 8 min 24 s but disappeared at 8 min 52 s. Bar, 2 μm. (B) Fluorescence microscopy images of strain Edc3-C/DAS1m-V after treatment with protein synthesis inhibitors. The strain incubated on an SM medium was treated with either 0.1 mg/mL cycloheximide (CHX) or 0.1 mg/mL puromycin (PMC) before microscopic observation. Merged images were generated by combining Venus and Cerulean fluorescence images. Bar, 2 µm. (C) Quantification of dot formation of Edc3-Cerulean and Venus-tagged *DAS1* mRNA and their colocalization detected in (**B**). Cell count analysis was performed on fluorescence microscopy images (n = 30, n: number of cells analyzed; b = 3, b: technical replicates; total = 90 cells). The number of dots is presented as mean ± SE of 90 cells and colocalization rates are shown as mean ± SE from three technical replicates (n = 30 cells per replicate). Asterisks indicate statistical significance. ** *p* < 0.01. n.s. means not significant. (D) Fluorescence microscopy images of strain Edc3-C/DAS1m-V after treatment with 1,6-hexandiol (HXD). The strain incubated on an SM medium was treated with 10% HXD before microscopic observation. Merged images were generated by combining Venus and Cerulean fluorescence images. Bar, 2 µm. (E) Quantification of dot formation of Edc3-Cerulean and Venus-tagged *DAS1* mRNA and their colocalization detected in (**D**). Cell count analysis was performed on fluorescence microscopy images (n = 30, n: number of cells analyzed; b = 3, b: technical replicates; total = 90 cells). The number of dots is presented as mean ± SE of 90 cells and colocalization rates are shown as mean ± SE from three technical replicates (n = 30 cells per replicate). Asterisks indicate statistical significance. ** *p* < 0.01.

To investigate how mimRNAs colocalized with P-bodies respond to translational regulation, we treated strain Edc3-C/DAS1m-V with puromycin (PMC) and cycloheximide (CHX) (Riggs et al., 2020). PMC promotes the accumulation of ribosome-free mRNAs (Cougot et al., 2004; Zheng et al., 2008), thereby promoting RNP granule formation, including P-bodies (Eulalio et al., 2007; Kedersha et al., 2005). Consistent with previous studies, PMC treatment increased the number of Edc3-Cerulean dots. We also observed that their colocalization with Venus-tagged *DAS1* mRNA dots increased without significantly altering the total number of *DAS1* mRNA dots (Figure 5B, C). Similar treatment was performed with the strain expressing Cerulean-tagged Edc3 and Venus-tagged *TDH3* mRNA (strain Edc3-C/TDH3m-V). Upon PMC treatment, the colocalization rate of Edc3-Cerulean with *TDH3* mRNA increased from 10% to 23%, which was lower than that of *DAS1* mRNA (29% to 48%) (Figure 5–figure supplement 1A-B). CHX is known to stall ribosomes on mRNAs, which prevents the accumulation of ribosome-free mRNAs (Schneider-Poetsch et al., 2010) and inhibits the formation of RNP granules, including P-bodies (Andrei et al., 2005; Grousl et al., 2009; Kedersha et al., 2000). As expected, CHX suppressed Edc3-Cerulean dot formation and reduced colocalization with Venus-tagged *DAS1* mRNA dots, while partially reducing the number of *DAS1* mRNA dots (Figure 5B, C). A similar reduction in colocalization was observed in strain Edc3-C/TDH3m-V upon CHX treatment (Figure 5–figure supplement 1A-B).

In addition, we treated strain Edc3-C/DAS1m-V with 1,6-hexanediol (HXD), which is commonly used to disrupt LLPS-mediated granules (Kroschwald et al., 2015). Similar to CHX, HXD treatment inhibited the formation of Edc3-Cerulean dots and partially reduced the number of Venus-tagged *DAS1* mRNA dots (Figure 5D, E). These results suggest that some mimRNA granules are resistant to HXD, while P-bodies are significantly suppressed, further indicating that mimRNAs can also accumulate at P-body-independent sites. We also examined the effect of HXD on strain Edc3-C/TDH3m-V. Although HXD treatment significantly reduced the number of Edc3-Cerulean dots, it did not affect the number of Venus-tagged *TDH3* mRNA dots (Figure 5–figure supplement 1C-D), suggesting that the colocalization with P-bodies differs between methanol-induced and non-methanol-induced mRNA granules during methanol-utilizing conditions.

As some mimRNA granules were resistant to HXD, we explored potential locations of mimRNAs other than P-bodies. The endoplasmic reticulum (ER) and peroxisomes were visualized using mCherry-tagged localization sequences (Gould et al., 1989; Lajoie et al., 2012) and actin patches (Amberg, 1998) were stained with rhodamine-phalloidin. Venus-tagged *DAS1* mRNA dots were also detected in association with these compartments (Figure 5–figure supplement 2A-D), raising the possibility of additional sites of intracellular localization for mimRNAs.

## Discussion

We demonstrate that the P-body component Edc3 is required for the proliferation of methylotrophic yeast *C. boidinii* in the phyllosphere where the yeast uses methanol as the carbon source while adapting to varying environmental conditions. In the leaf environment, Edc3-dependent functions of P-bodies may contribute to post-transcriptional regulation of methanol-induced genes. Methanol triggers the transcription of genes involved in methanol assimilation, leading to the accumulation of mimRNAs in the cytosol. Based on the responses to translation inhibitors, these mimRNAs form cytosolic “mimRNA granules”, which may contain both ribosome-free and ribosome-associated transcripts. These granules are partially colocalized with P-bodies. Under stresses such as oxidative stress, mimRNAs colocalize more frequently with P-bodies, indicating more efficient sequestration of mimRNAs by P-bodies. Upon stress removal, the colocalization between these granules decreases, potentially allowing mimRNAs to be translated into the cytosol. These findings suggest that Edc3-dependent functions of P-bodies contribute to the regulation of mimRNAs in response to environmental changes, supporting yeast adaptation to changing leaf environments (Figure 6).

**Figure 6.**
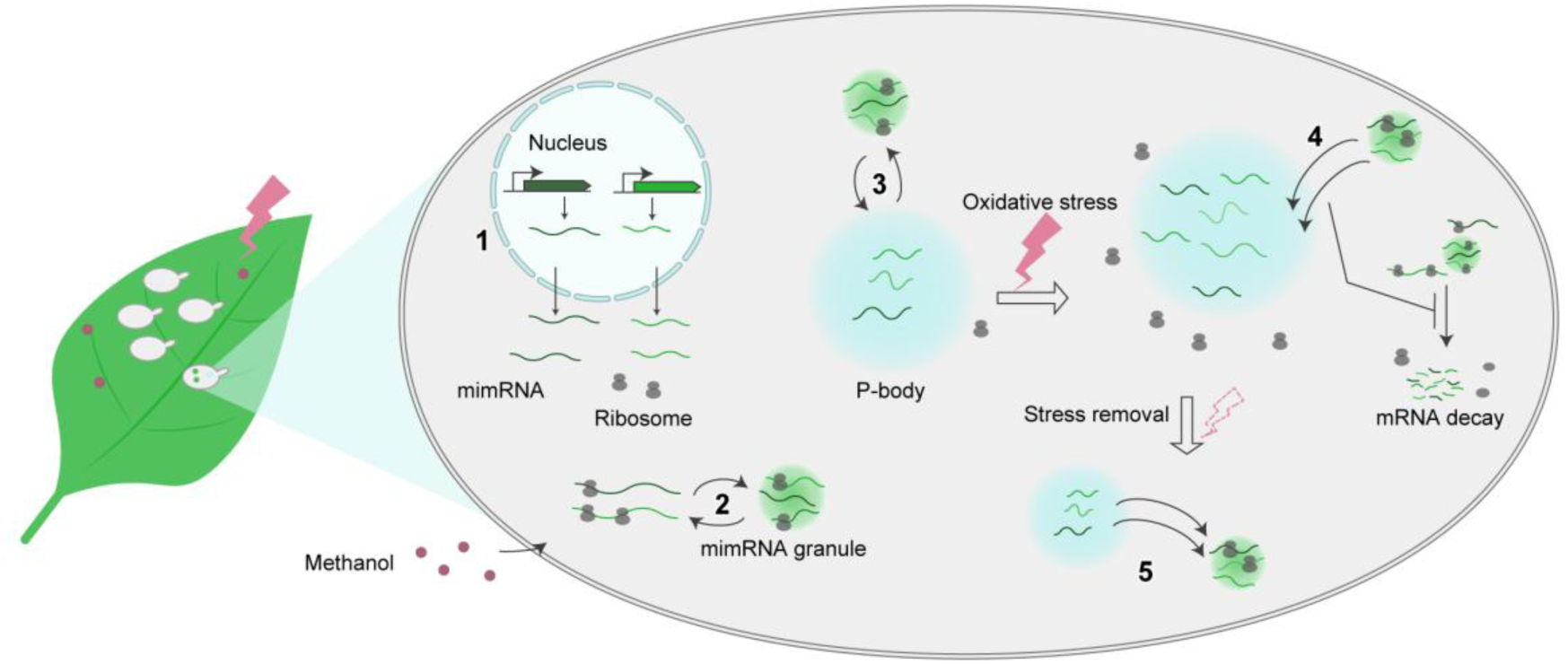
Schematic image illustrating the role of yeast Edc3-dependent functions of P-bodies in the post-transcriptional regulation of methanol-induced genes in the phyllosphere. On *A. thaliana* leaves, *C. boidinii* assimilates methanol as the carbon source while adapting to environmental stresses. In this leaf environment, Edc3-dependent functions of P-bodies contribute to regulating methanol-induced genes post-transcriptionally through the following steps: 1) Methanol induces transcription of genes involved in methanol metabolism leading to the accumulation of methanol-induced mRNAs (mimRNAs) in the cytosol; 2) Multiple mimRNAs accumulate into cytosolic mimRNA granules that may contain both ribosome-free and ribosome-associated mimRNAs; 3) mimRNA granules dynamically interact with P-bodies where ribosome-free mimRNAs may partially accumulate; 4) In response to environmental stimuli such as oxidative stress, mimRNAs colocalize more frequently with P-bodies, possibly sequestration for protecting them from degradation; 5) Upon stress removal, P-bodies release mimRNAs to the cytosol potentially for translation.

Our observation that Venus-tagged *DAS1* mRNA colocalized with Edc3-Cerulean on *Arabidopsis* leaves (Figure 2A–C and Figure 2–figure supplement 3), together with *in vitro* results, including their dynamic properties during transient colocalization (Figure 5A, Figure 5–video 1), supports our speculation: P-bodies sequester mimRNAs for protection from degradation and release them into the cytosol for translation in response to environmental cues. This mechanism could promote energy conservation by preserving critical transcripts and minimizing the need for constant mRNA synthesis.

This interpretation aligns with previous reports showing that highly expressed mRNAs are often sequestered in P-bodies (Lavut & Raveh, 2012) and that stored mRNAs can re-enter the translational pool when conditions change (Bhattacharyya et al., 2006; Brengues et al., 2005; Parker & Sheth, 2007). However, direct evidence that P-bodies protect mimRNAs from degradation and release them into the cytosol for translation in *C. boidinii* remains to be obtained.

We examined mimRNA granule formation and their colocalization with P-bodies on *Arabidopsis* leaves under light and dark conditions. Although methanol concentrations are higher during the dark period (Kawaguchi et al., 2011), both the number of mimRNA granules (2.4–2.7 dots per cell) and the colocalization rate with P-bodies (∼13%) remained low and largely unchanged between light and dark conditions (Figure 2C), and the number of granules was lower than that in SM medium (3.5–5.9 dots per cell; Figure 2E). Partial colocalization of mimRNA granules and P-bodies was also observed on wilting leaves (Figure 2–figure supplement 3), despite their elevated methanol levels (Kawaguchi et al., 2011). This contrasts with *in vitro* assays (Figure 4–figure supplement 2) where mimRNA granule formation clearly increased with higher methanol concentrations, i.e., 0.5% versus 0.005%. These findings suggest that granule formation does not depend solely on methanol in the phyllosphere, but is influenced by additional environmental factors, such as stress-specific signaling. Additionally, mimRNAs may be rapidly directed to translation, limiting their accumulation into granules or P-bodies to survive under nutrient-scarce conditions.

The spatiotemporal regulation of mimRNAs appeared to be especially critical under stress conditions, notably oxidative stress (Figure 3). In the phyllosphere, microbial cells are exposed to ROS generated by photosynthesis and to sunlight, which can damage nucleic acids and other cellular components. Microbes deploy protective mechanisms such as pigment production (Jacobs et al., 2005) and DNA repair (Gunasekera & Sundin, 2006) for survival. Sequestration of mimRNAs in P-bodies may represent an additional protective strategy for these transcripts under exposure to ROS.

It is possible that P-bodies preferentially sequester mimRNAs over non-methanol-induced mRNAs under stress during growth on methanol: ∼60% of P-bodies were colocalized with mimRNAs under oxidative stress, compared to only 15% with *TDH3* non-methanol-induced mRNAs (Figure 3E, F, Figure 4–figure supplement 5). While treatment with HXD significantly reduced the number of *DAS1* mRNA dots, it did not affect the number of *TDH3* mRNA dots (Figure 5D-E, Figure 5–figure supplement 1C-D). This raises the possibility that P-bodies preferentially store mimRNAs during stress, while non-essential mRNAs are more likely targeted for degradation and recycling. Our interpretation is supported by previous studies showing that P-bodies regulate translation selectively to optimize cell survival (Jang et al., 2019; Sorenson & Bailey-Serres, 2014). However, the molecular mechanisms governing the selective sequestration of mimRNAs in methylotrophic yeasts under methanol growth conditions remain to be elucidated. The principle of selectivity in cellular regulation extends beyond RNA. Our previous study showed that peroxisomes undergo selective autophagy, i.e., pexophagy (Oku & Sakai, 2016) in *C. boidinii* on the leaf surface in response to light/dark cycles, while non-selective bulk autophagy, i.e., macroautophagy occurs continuously (Kawaguchi et al., 2011). Similarly, the yeast nitrate reductase Ynr1 escapes degradation via a selective autophagy termed the cytoplasm-to-vacuole targeting (CVT) pathway (Hutchins & Klionsky, 2001; Scott et al., 1997) on growing leaves but becomes targeted on aged leaves (Shiraishi et al., 2015). Together, these studies illustrate a microbial strategy: the selective preservation and deployment of cellular components in response to environmental changes and nutrient availability.

While our results highlight the importance of Edc3-dependent functions P-bodies in mimRNA regulation, we do not exclude their broader roles in supporting yeast proliferation in the phyllosphere. Since P-bodies influence the quality and fate of various transcripts (Wang et al., 2018), they may also sequester other mRNAs essential for yeast proliferation, including those involved in autophagy (Kawaguchi et al., 2011). Beyond RNA regulation, P-bodies can safeguard a key kinase protein under low nutrient conditions, protecting it from degradation and enabling proper stress responses (Zhang et al., 2018). Similar protective roles may apply to other essential proteins in the phyllosphere. Furthermore, yeast P-bodies interact with other RNP granules, such as hybrid granules induced by hypoosmotic stress and SGs (Escalante & Gasch, 2021), suggesting that loss of P-body formation could disrupt important inter-granule communication. In particular, Edc3 functions as an enhancer of mRNA decapping, in addition to P-body formation. As such, it is possible that decapping-related functions of Edc3 contribute to the phenotypes of the *edc3Δ* strain observed in the present study. Taken together, while mimRNA sequestration is a key function, Edc3-dependent functions of P-bodies are likely to support yeast adaptation to the phyllosphere through multiple mechanisms: regulating mRNA turnover beyond mimRNAs, stabilizing key proteins and engaging in dynamic interactions with other RNP granules to meet the challenges of complex plant-surface environments.

In addition to P-bodies, mimRNA accumulation was observed at membrane-bound organelles such as the ER and peroxisomes (Figure 5–figure supplement 2). The localization of mRNAs to these compartments is increasingly recognized as a key layer of spatial control for translation (Dahan et al., 2022; Morales-Polanco et al., 2021; Pilaz et al., 2023; Singer-Krüger & Jansen, 2014). Various mRNAs have been reported to be targeted to the ER for co-translational processing (Aronov et al., 2007; Fundakowski et al., 2012; Reid & Nicchitta, 2015). A recent study showed that up to 80% of *PEX* mRNAs that encode peroxins essential for the formation and function of peroxisomes are translated locally at peroxisomes (Dahan et al., 2022; Zipor et al., 2009). Notably, only about 30% of *DAS1* mRNA localized to peroxisomes (Figure 5–figure supplement 2), suggesting that mimRNAs such as *DAS1* and *AOD1* are primarily translated at the ER, with their protein products subsequently transported to peroxisomes via peroxisomal targeting sequences (PTS) (Gould et al., 1989). This difference may reflect the temporal distinction in protein demand: peroxins are needed for peroxisome biogenesis whereas methanol-metabolizing enzymes like DAS function after peroxisomes are formed.

Our inhibitor experiments suggest that mimRNAs colocalized with P-bodies are in a ribosome-free state, consistent with previous studies (Eulalio et al., 2007). Treatment with PMC, which generates ribosome-free mRNAs, enhanced P-body formation and mimRNA colocalization, whereas CHX, which stalls ribosomes on mRNAs, suppressed both (Figure 5B, C). These results further support previous reports (Cougot et al., 2004; Zheng et al., 2008) showing that translationally inactive, ribosome-free mRNAs accumulate in P-bodies.

A limitation of this study is that many of our analyses of mimRNA regulation, including time-course and time-lapse imaging, stress treatments, inhibitor treatments and transcript quantification, were performed under synthetic medium conditions. These analyses were undertaken to characterize the processes observed in the phyllosphere in greater detail, because such experiments are technically difficult to perform on leaf surfaces. Therefore, whether the regulatory mechanisms observed *in vitro* actually operate in the phyllosphere remains to be determined.

In conclusion, our study demonstrates that the P-body component Edc3 is required for yeast proliferation in the phyllosphere. P-bodies may function as hubs for the temporary storage of mimRNAs, enabling yeast cells to respond rapidly to the complex and changing phyllosphere environment. These findings advance our understanding of post-transcriptional regulation and RNP granule biology in natural microbial ecosystems, while also opening new avenues for investigating eukaryotic microbial dynamics.

## Materials and methods

## Appendix 1: Key resources table

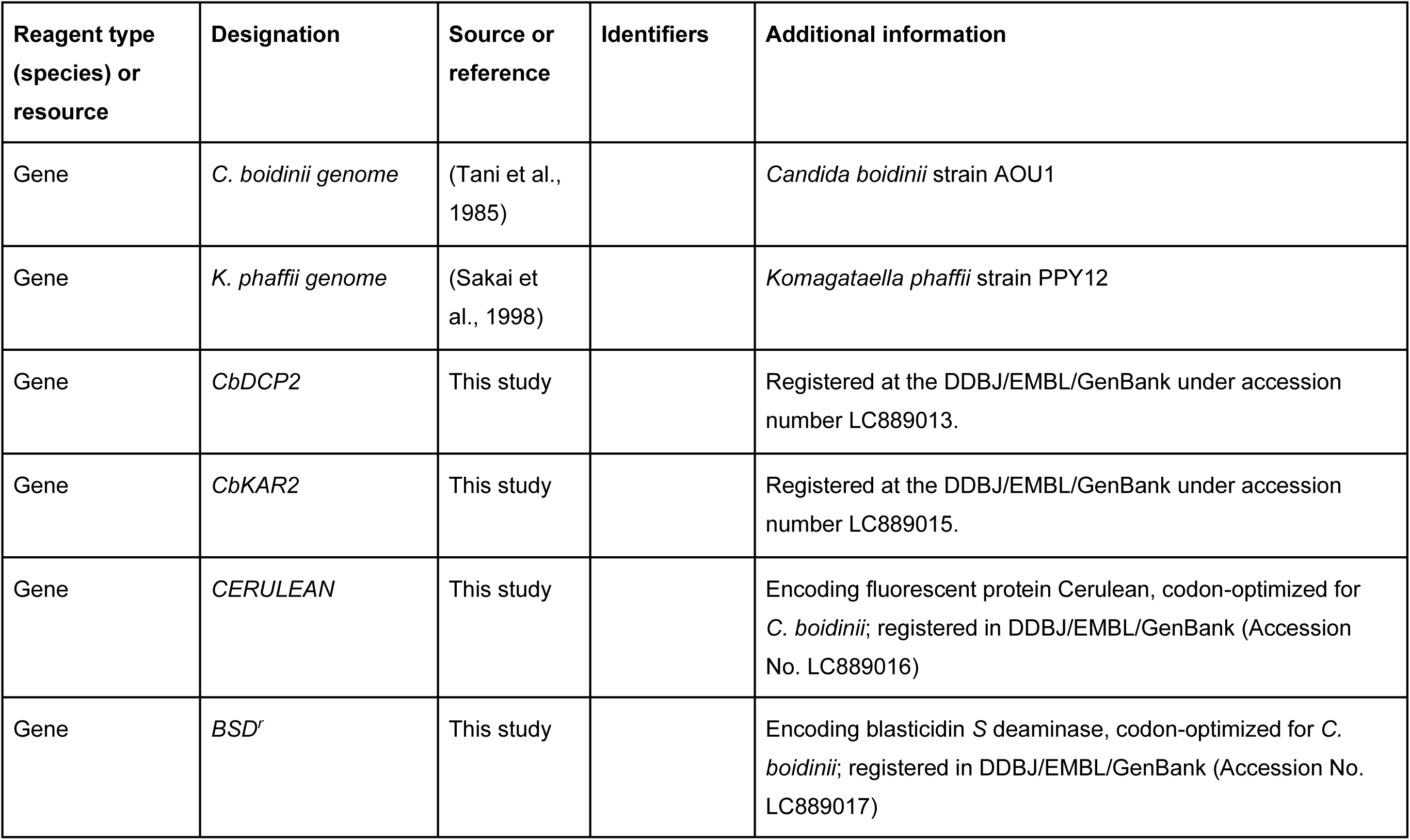
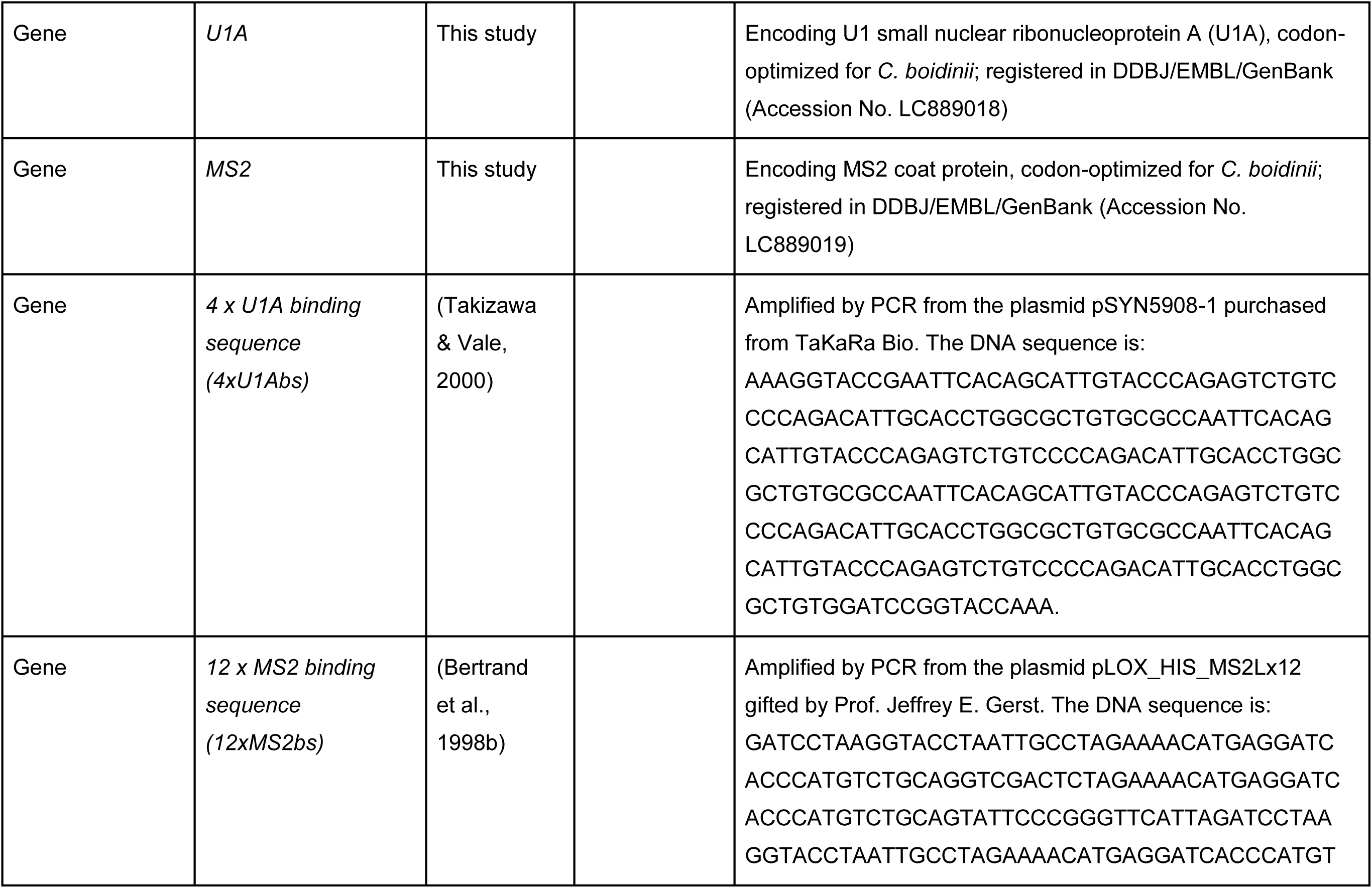
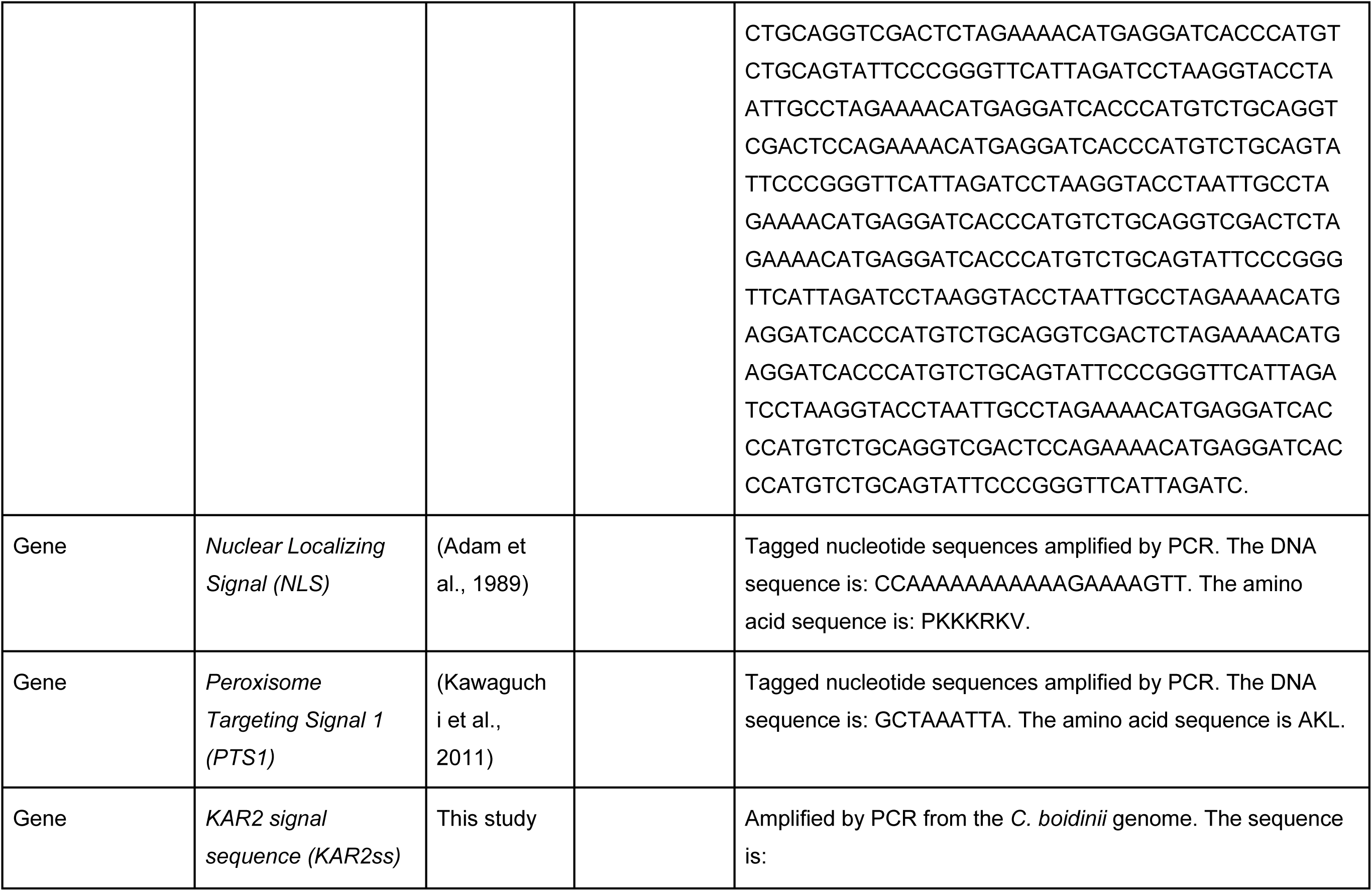
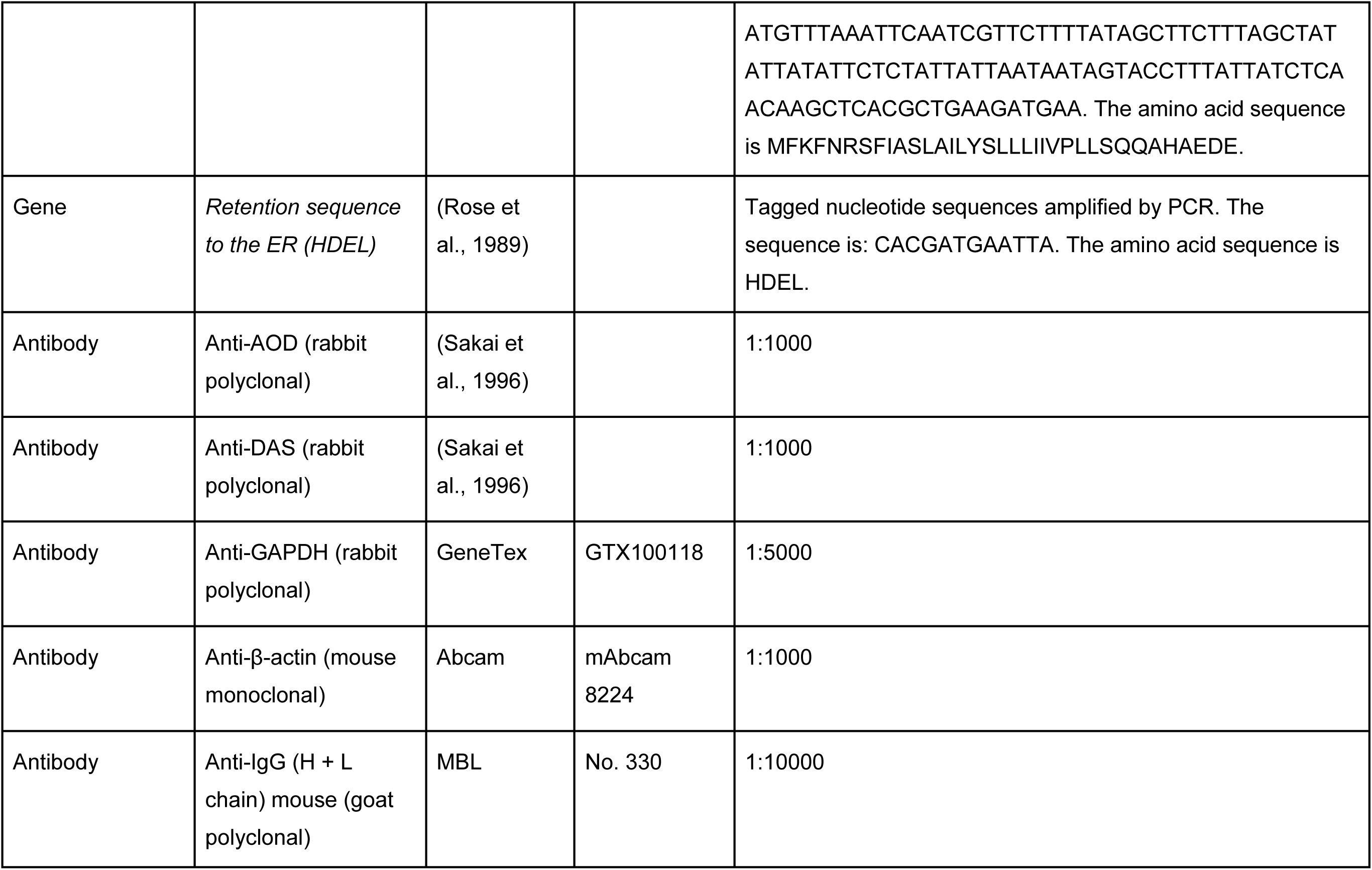
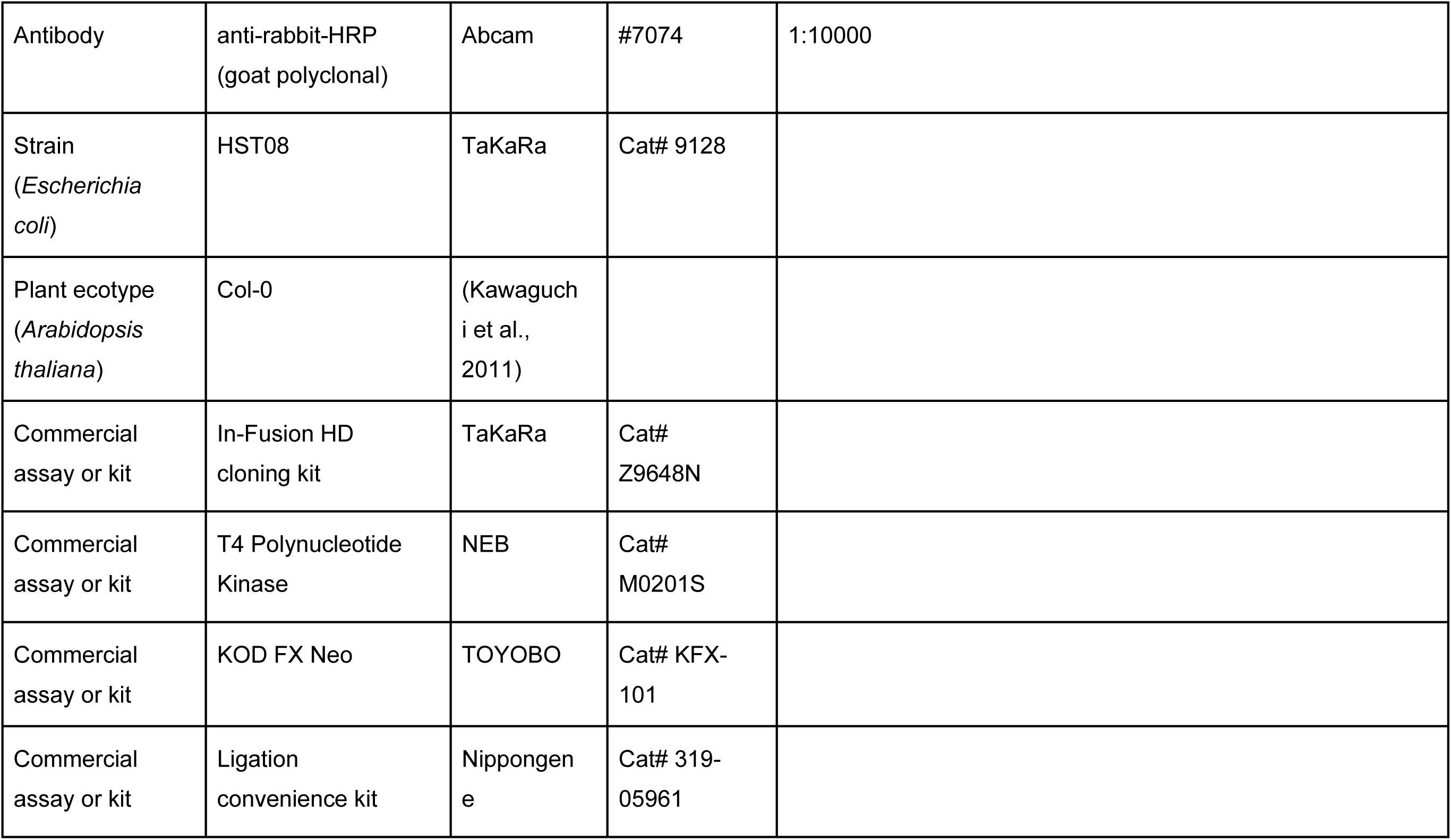
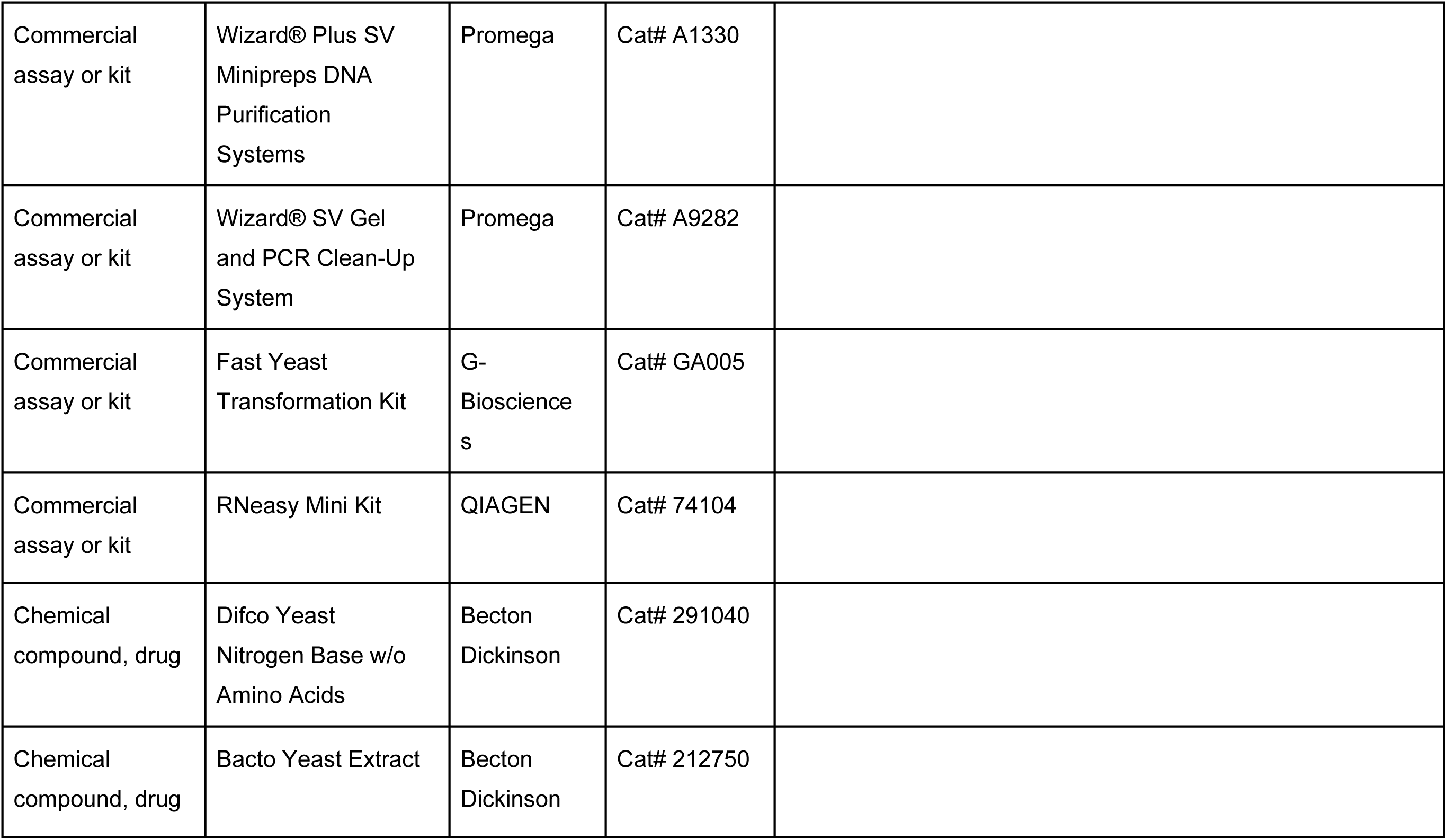
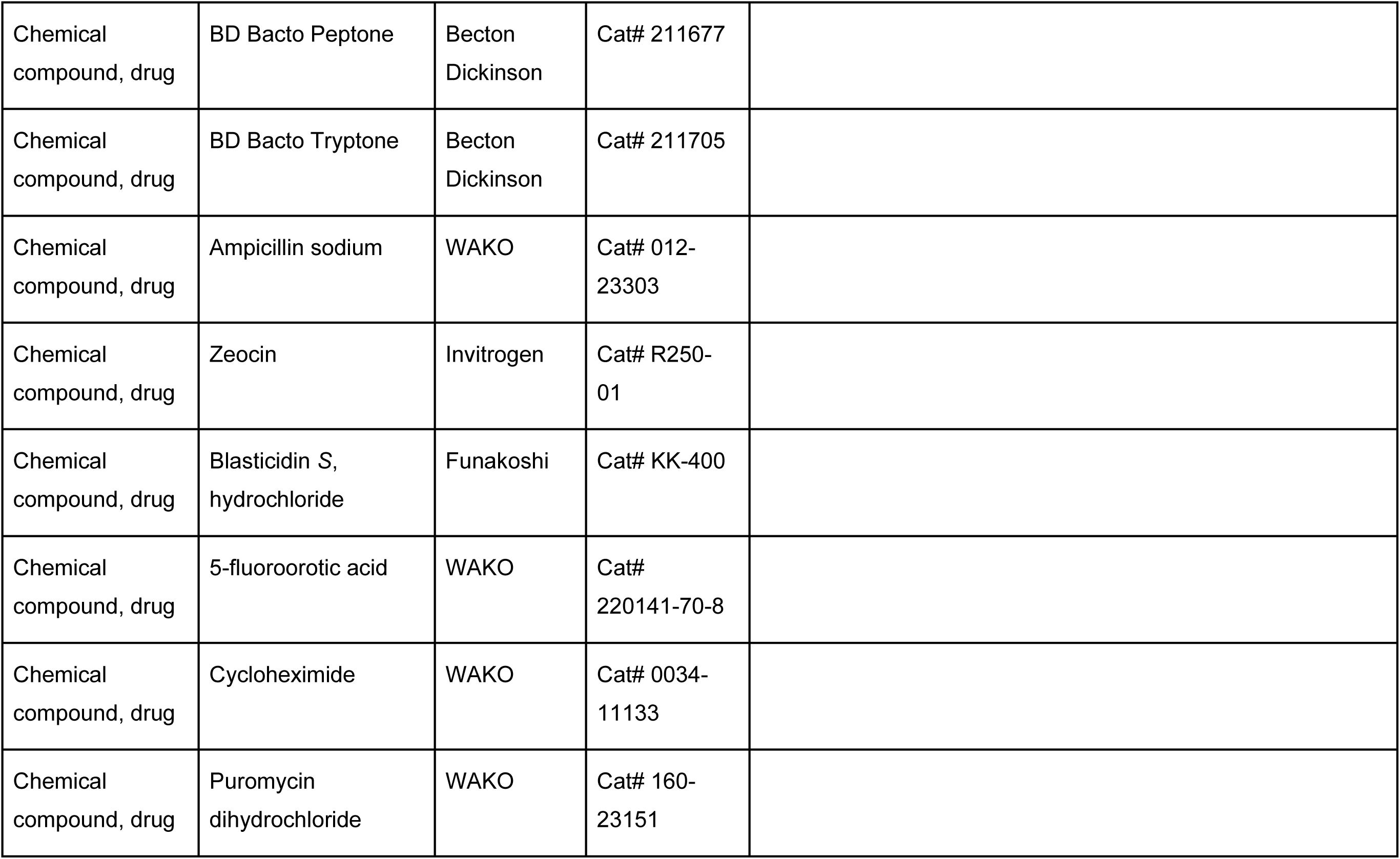
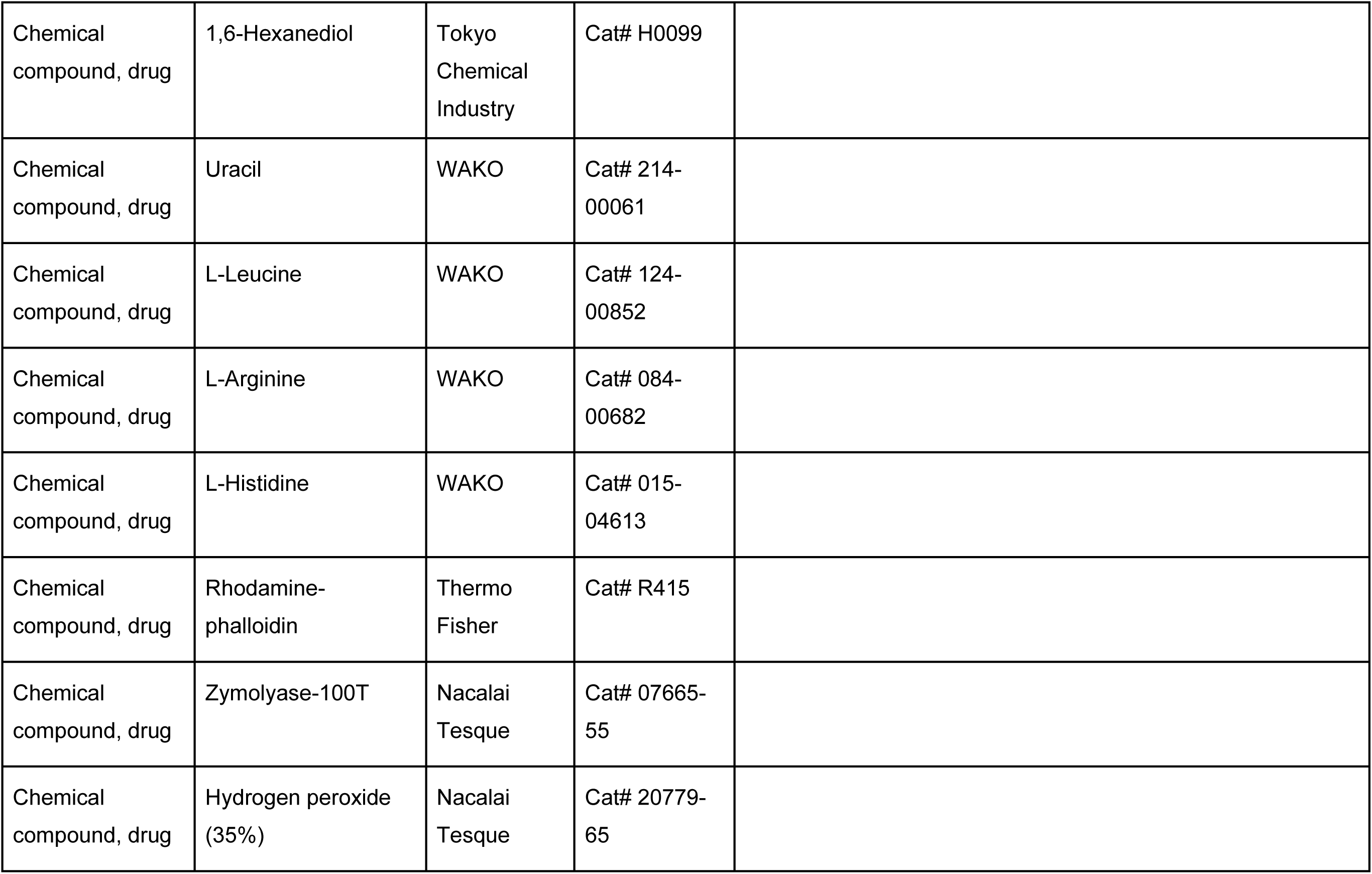
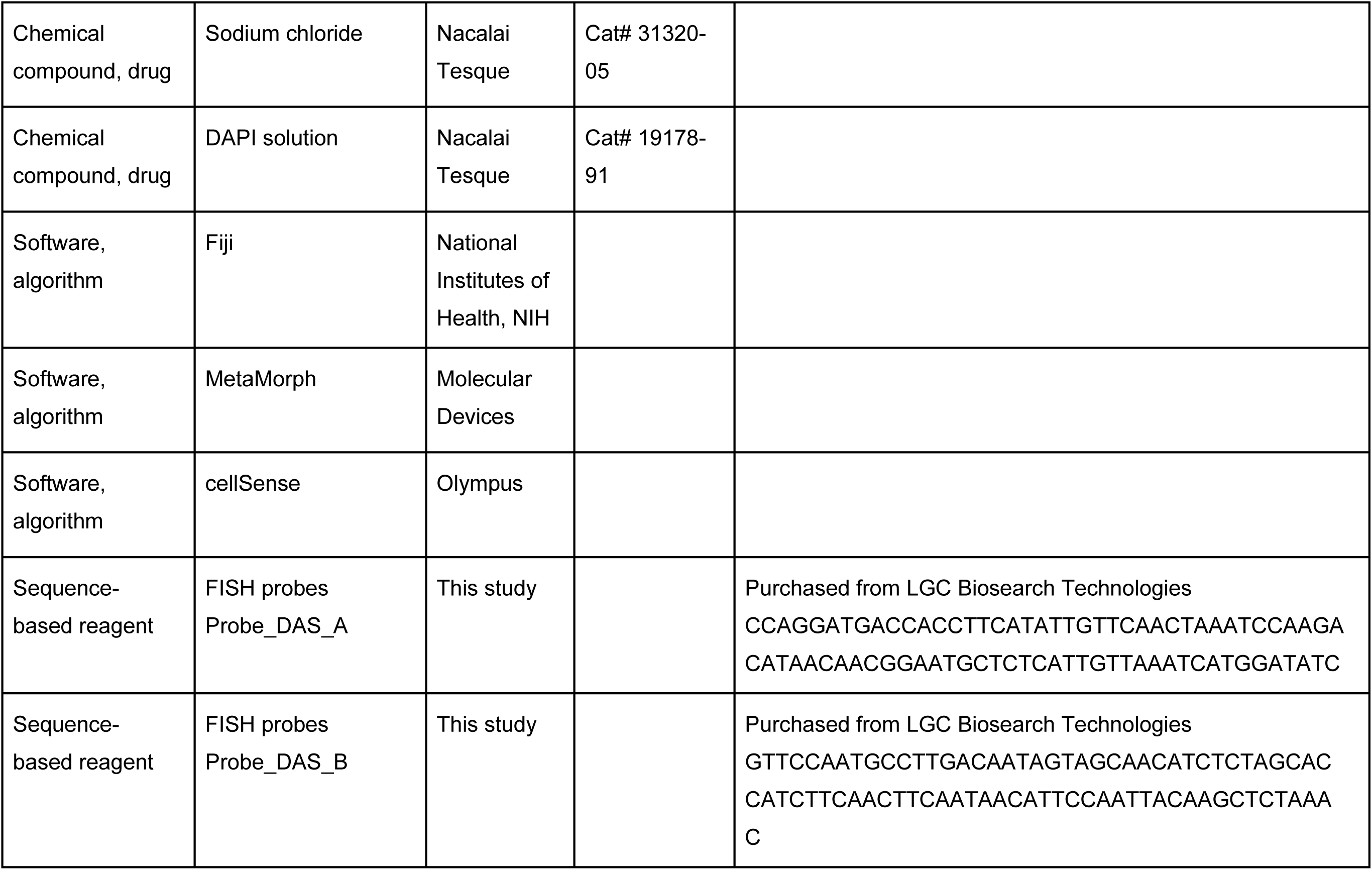
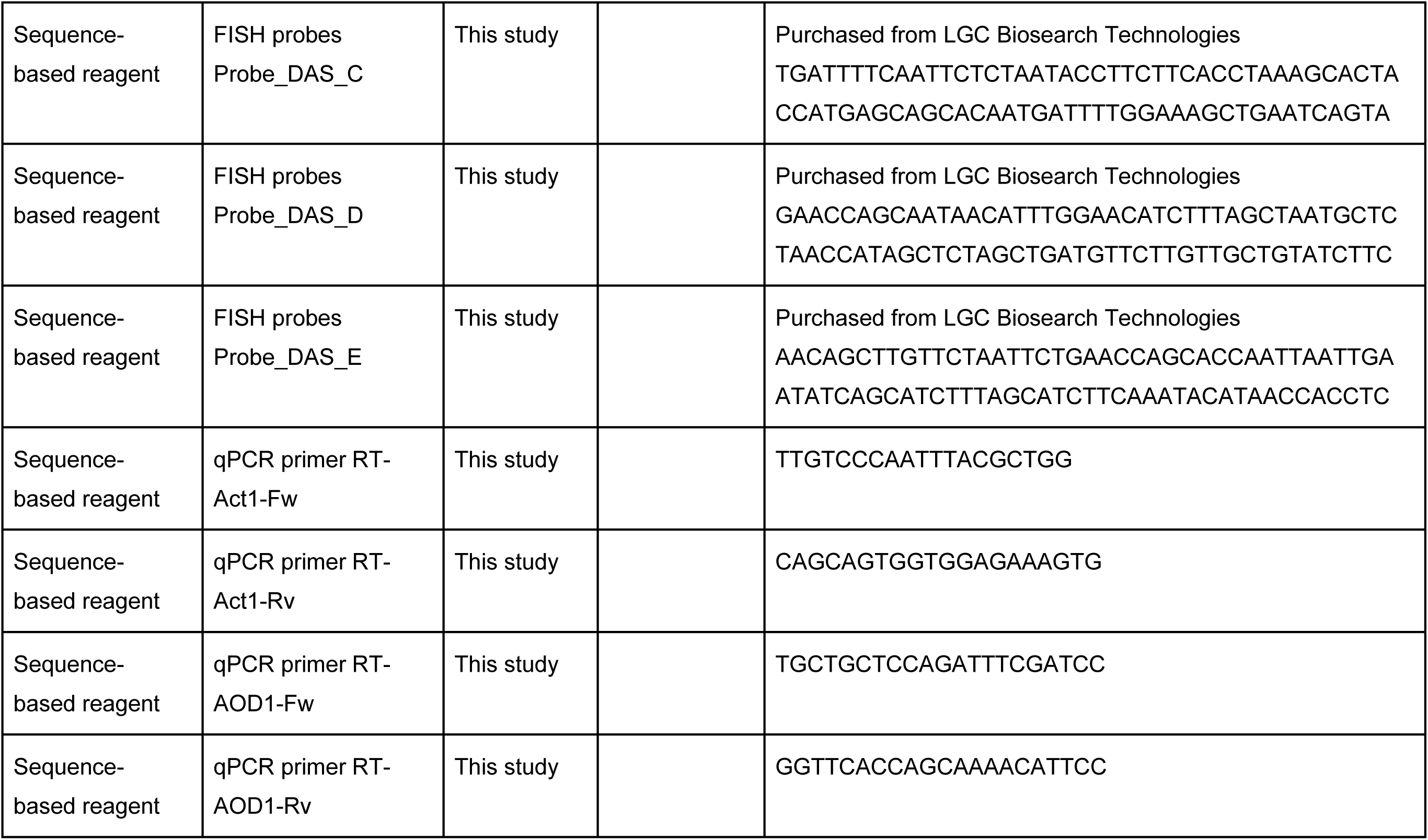
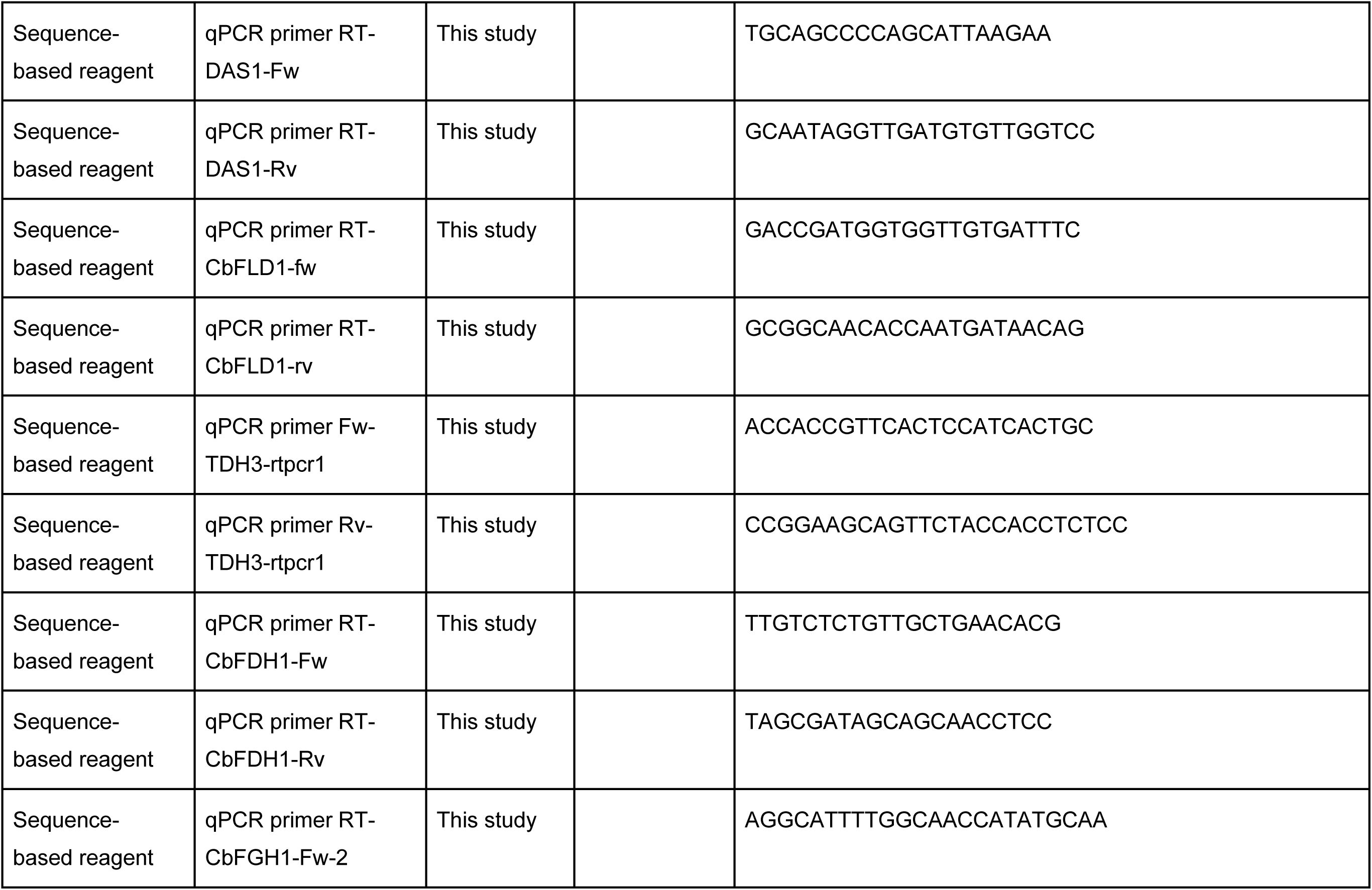
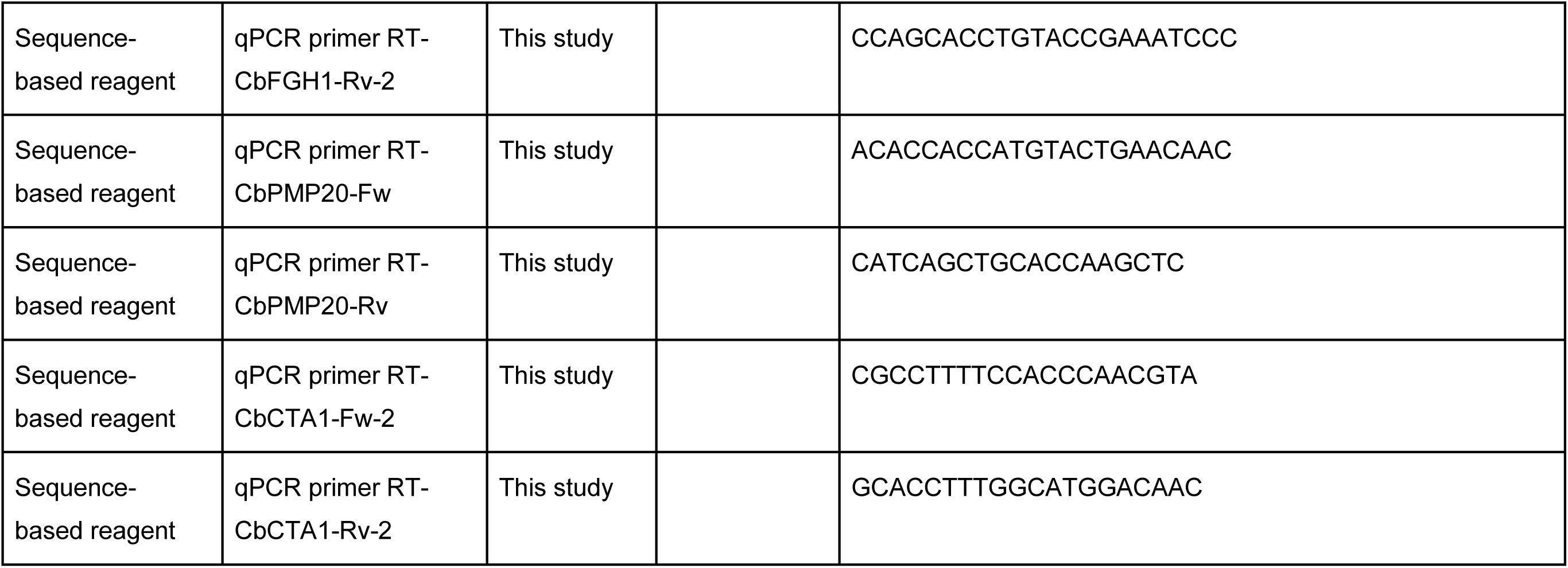

### Yeast strains and culture conditions

Yeast strains used in this study are listed in Supplementary file Table S1. Cells were grown at 28°C in the media described below. YPD (yeast extract peptone dextrose) medium consisted of 1% yeast extract, 2% bactopeptone and 2% glucose. Synthetic media were prepared with 0.67% yeast nitrogen base without amino acids and supplemented with one of the following carbon sources: 2% glucose for SD (synthetic dextrose), 2% ethanol for SE (synthetic ethanol) or 0.7% methanol for SM (synthetic methanol) medium. These media were supplemented with the appropriate amino acids (20 μg/mL uracil, 100 μg/mL leucine, 100 µg/mL arginine and/or 100 µg/mL histidine) and/or antibiotics (50 μg/mL zeocin or 100 μg/mL blasticidin) and/or 0.08% w/v of 5-fluoroorotic acid (5-FOA), as necessary. The pH of SD, SE and SM media was adjusted to 6.0 with NaOH. Cell growth was monitored by measuring the optical density at 610 nm (OD_610_). Solid media were prepared by the addition of 2% agar to the appropriate liquid media.

*Escherichia coli* HST08 premium competent cells (TaKaRa Bio Inc., Shiga, Japan) were used for gene cloning. *E. coli* was grown at 37°C on LB medium (1% Bacto tryptone, 0.5% Bacto yeast extract and 0.5% NaCl) in the presence of ampicillin (50 μg/mL).

### DNA isolation and transformation

Plasmid DNA from *E. coli* was isolated using the Wizard® Plus SV Minipreps DNA Purification System (Promega, Madison, WI). Transformation of *E. coli* was carried out following the method of Dower et al (Dower et al., 1988). Genomic DNA from yeast was extracted using the method of Cryer et al (Cryer et al., 1975). Transformation of *C. boidinii* was performed using the Fast Yeast Transformation Kit (G-Biosciences, Maryland Heights, MO) or a modified lithium acetate method (Sakai & Tani, 1992), and *K. phaffii* transformation was conducted as previously described (Higgins & Cregg, 1998).

### Plasmid construction

Primers used in this study are listed in Supplementary file Table S2, and plasmids used in this study are listed in Supplementary file Table S3. Details of the plasmid construction are described in Supplementary file.

### Visualization of mRNAs in fixed cells

Fluorescence *in situ* hybridization (FISH) was performed using the method previously described (McIsaac et al., 2013) with Stellaris^®^ RNA FISH Probe (LGC Biosearch Technologies, Petaluma, CA). Custom oligonucleotide probes complementary to the target mRNA were designed with Cy5 conjugated at the 5′ end. Probe sequences are listed in Key Resources Table. Dried probes were dissolved in TE buffer (10 mM Tris-HCl, 1 mM EDTA, pH 7.0) and stored at −20°C before use.

Briefly, yeast cells were cultured in SM medium and harvested at 3 hours after the start of cultivation. Cells were then fixed with 35% formaldehyde at a final concentration of 3.5% (v/v), washed with fixation buffer (1.0 M Sorbitol, 0.1 M K_2_HPO_4_) and resuspended in fixation buffer supplemented with Zymolyase for spheroplastization. After washing with fixation buffer, cells were resuspended in 70% ethanol and stored at 4°C overnight. Subsequently, fixed cells were washed with SSC-formamide wash buffer, and then hybridized overnight at 37°C in hybridization buffer containing the Cy5-labelled probe at a final concentration of 12.5 µM. Finally, cells were resuspended in wash buffer, mounted on glass slides, and observed by fluorescence microscopy to determine the subcellular localization of target mRNAs based on Cy5 signal distribution.

### Visualization of mRNAs in live cells

To visualize mRNAs in living cells, we employed both the U1A-based and MS2-based RNA systems. In the U1A system, the human U1A RNA-binding protein was used to tether the fluorescent protein Venus to target mRNAs (Chung & Takizawa, 2011). Four tandem repeats of the U1A binding sequence (U1Abs) were directly inserted into the 3ˈ untranslated region (3ˈUTR) of the target genes at their endogenous genomic loci. The localization of these tagged mRNAs was monitored by co-expression of a fusion protein consisting of U1A and Venus with a nuclear localization signal (NLS) at its N terminus (NLS-U1A-Venus) (see Figure 2–figure supplement 2 for schematic diagram).

In parallel, the MS2 system was also used to visualize mRNAs where the bacteriophage-derived MS2 coat protein (MCP) was used to tether the fluorescent protein Cerulean to target mRNAs (Bertrand et al., 1998a). Twelve tandem repeats of the MS2 binding sequence (12xMS2bs) were directly inserted into the 3’UTR sequences of the target genes at their endogenous genomic loci. The localization of these tagged mRNAs was monitored by co-expression of the NLS-MCP-Cerulean.

### Fluorescence microscopy

Two microscopy systems were used for fluorescence imaging: a fluorescence microscope and a confocal laser scanning microscope.

Fluorescence microscopy was performed using the IX81 inverted fluorescence microscope (Olympus, Tokyo, Japan) equipped with an XF52 filter set (Omega Optical Inc., Brattleboro, VT) suitable for detecting Cy5, DAPI, Rhodamine-Phalloidin and fluorescent proteins. Images were captured with a SenSys^TM^ charged-coupled device (CCD) camera (PhotoMetrics, Tucson, AZ) and analyzed on MetaMorph software (Universal Imaging Corporation, Downingtown, PA), Fiji (Schindelin et al., 2012) and Adobe Photoshop (Adobe Inc., San Jose, CA). Scale bars in figures represent the indicated size.

Time-lapse imaging was performed using the IX83 inverted fluorescence microscope (Evident, Tokyo, Japan) equipped with an IX3-SSU motorized stage (Evident), a drift compensation module (IX3-ZDC, Evident) and a filter wheel unit (96A357 and MAC6000, Ludl Electronic Products Ltd., Hawthorne, NY, USA) suitable for detecting fluorescent proteins. Images were captured with ORCA-Fusion Digital CMOS camera (USB3.0 set, Hamamatsu Photonics, Hamamatsu, Japan) and analyzed on cellSens software (Evident) and Fiji. Cells were cultured under continuous medium flow using the CellASIC® ONIX2 Microfluidic System (Merck, Darmstadt, Germany) with the CellASIC® ONIX B04A-03 Microfluidic Bacteria Plate (Merck), according to the manufacturer’s instructions. Images were acquired every 28 seconds for 20 cycles.

Confocal microscopy was conducted using the FV3000 confocal microscope (Olympus) equipped with six solid-state diode lasers (405, 445, 488, 514, 561 and 640 nm). The system utilizes high-numerical-aperture objectives, including UPLSAPO 100× (NA 1.35, silicone oil immersion; Olympus). Fluorescence signals were detected with high-sensitivity GaAsP detectors, allowing for the capture of faint signals with high signal-to-noise ratios. Venus fluorescence was excited at 488 nm and detected with a 500–600 nm emission filter, whereas Cerulean fluorescence was excited at 445 nm and detected with a 460–500 nm emission filter. Differential interference contrast (DIC) images were acquired using transmitted light with Nomarski optics. The images were captured and processed using cellSens software (Olympus), Fiji and Photoshop.

### Morphometric and statistical analyses

Cell counts were analyzed from fluorescence microscopy images (n = 30 cells per technical replicate, with 3 technical replicates). Cytosolic dots were counted manually, as described previously (Shiraishi et al., 2018). Fluorescent foci that were clearly distinguishable from the diffuse cytosolic signal were counted as dots. Colocalization was defined as the overlap of dots in merged images. In total, 90 cells were analyzed to quantify dot numbers and colocalization rates. Statistical significance was assessed using Welch’s unpaired two-tailed t-test.

### 1,6-hexanediol treatment

1,6-hexanediol (HXD) treatment was performed using the method previously described (Kroschwald et al., 2017). HXD disrupts weak hydrophobic interactions such as those involved in phase-separated biomolecular condensates. Cells were pre-cultured in SD medium for 12 hours and then transferred to SM medium. After 3 hours of methanol induction, 800 μL of cell culture was collected. Then, digitonin was added to a final concentration of 10 μg/mL, mixed gently and incubated for 5 minutes to mildly permeabilize the plasma membrane. Subsequently, 200 μL of 50% (v/v) HXD was added to achieve a final concentration of 10%. After a 1-minute incubation, cells were immediately subjected to fluorescence microscopy to assess the dissolution of condensates or subcellular structures.

### Cycloheximide/puromycin treatment

Cells were pre-cultured in SD medium for 12 hours and then transferred to SM medium containing either cycloheximide (CHX) or puromycin (PMC) at a final concentration of 0.1 mg/mL. After the addition, cells were subjected to fluorescence microscopy at the indicated time points. CHX was used to stall ribosomes on mRNAs, preserving translation complexes, while PMC was used to release nascent peptides by causing premature termination. These treatments were used to examine the effects of translational inhibition.

### Nuclear staining

After cultivation for the indicated time period, formaldehyde was added to the culture at a final concentration of 5% (v/v) to fix the cells. Following a 30-minute incubation, cells were harvested and washed twice with 1 mL of phosphate-buffered saline (PBS: 137 mM NaCl, 2.7 mM KCl, 10 mM Na_2_HPO_4_, 1.8 mM KH_2_PO_4_, pH 7.4). The fixed cells were then resuspended in 200µl of DAPI (4’,6-diamidino-2-phenylindole) solution at a final concentration of 0.1 μg/mL. After 20 minutes of incubation, cells were then washed twice with 1 mL of PBS and were used for fluorescence microscopy.

### Actinpatch staining

After cultivation for the indicated time, formaldehyde was added to the culture at a final concentration of 5% (v/v) to fix the cells. Following a 30-minute incubation, cells were harvested and washed three times with 1 mL of PBS. The fixed cells were then resuspended in 150 μL of PBS containing 3 μL of Rhodamine-Phalloidin and incubated for 2 hours at room temperature. After staining, cells were washed three times with 150 μL of PBS and used for fluorescence microscopy.

### Preparation of cell-free extract and qRT-PCR analysis

Extraction of total RNA and cDNA synthesis were performed as previously described (Ohsawa et al., 2017). qRT-PCR was performed with SYBR Premix Ex Taq (Takara Bio Inc.) using QuantStudio 1 Real-Time PCR System (Thermo Fisher Scientific, Waltham, MA). Transcript levels of all genes were normalized to *ACT1*. The primers for *ACT1*, *AOD1*, *DAS1*, *FLD1*, *FGH1*, *FDH1*, *CTA1* and *PMP20* are listed in the Key Resources Table.

### Preparation of protein extracts from yeast cells and immunoblot analysis

Protein extracts from yeast cells and immunoblot analysis were prepared as previously described (Shiraishi et al., 2024). The primary antibody and secondary antibody used in this study were anti-AOD (rabbit polyclonal) (Sakai et al., 1996), anti-DAS (rabbit polyclonal) (Sakai et al., 1996), anti-GAPDH (GTX100118, rabbit polyclonal, GeneTex, Irvine, CA), anti-βactin (mAbcam 8224, mouse monoclonal, Abcam, Cambridge, UK), Anti-IgG (H + L chain) mouse (No. 330, goat polyclonal, MBL Co., Ltd., Tokyo, Japan) and anti-rabbit-HRP (#7074, goat polyclonal, Abcam).

### Host plant growth conditions and yeast inoculation on plant leaves

*A. thaliana* plants were cultivated on mini-pots of cultivable lock-fiber (Nittobo, Tokyo, Japan) with Hoagland solution in growth chambers, as previously described (Kawaguchi et al., 2011). *C. boidinii* cells were cultured to OD_610_ = 1.0. To follow yeast proliferation on the leaves of *A. thaliana*, 1 µl of yeast cell suspension (OD_610_ = 0.5) was spotted onto the upper sides of the leaves. To observe Venus and/or Cerulean expression in *C. boidinii*, 5 µl of yeast suspension (OD_610_ = 0.5) was spotted onto the upper sides of the leaves. Then, the yeast cells were incubated on *A. thaliana* leaves until they were harvested for analysis. For Figure 2B and C, cells were precultured in SM medium before inoculation.

### Quantitation of yeast cell numbers on A. thaliana

After *C. boidinii* cells were inoculated on the leaves of *A. thaliana* for the indicated days, 5 leaves were collected. These leaves were placed in a 1.5 mL tube, and then, 100 µL of PBS was added per 10 mg of leaf and vortexed for 15 minutes. 20 µL of this collection sample was mixed with 20 µL of BD^TM^ Liquid counting Beads (BD Biosciences) and 200 µL of PBS and the total volume was used for flow cytometry (FCM) analysis to count the cell population. Quantitation of yeast cell numbers was performed as previously described, using the FACSAria™ III Cell Sorter (Becton Dickinson, Franklin Lakes, NJ) (Katayama et al., 2025). Briefly, fluorescent channel and light scatter were set at log gain. The forward scatter (FSC) was set at a photomultiplier tube (PMT) voltage of 2220 with a threshold of 200. The PMT voltages of side scatter (SSC) were set at 250 with a threshold of 200. Venus fluorescence was excited with a 488-nm laser and the emission at 530/30 nm was detected. Cells were counted referring to 2000 quantitative beads as standards. To ensure that FCM sorts Venus-labelled cells properly, we set gates for these labelled cells by using the Venus-labelled cells cultivated in SM medium. FACSDiva 8 software (Becton Dickinson) was used for data acquisition.

## Data availability

All sequence data obtained in this study have been deposited in the DDBJ Sequence Read Archive. The accession numbers for the sequence data are LC889013 (*CbDCP2*), LC889015 (*CbKAR2*), LC889016 (*CERULEAN*), LC889017 (*BSD^r^*), LC889018 (*U1A*) and LC889019 (*MS2*). All data generated or analyzed during this study are included in the manuscript and supporting files; Source Data files have been provided for Figures 1–5, and Figure 1–figure supplement 1, Figure 2–figure supplement 2, Figure 2–figure supplement 3, Figure 2–figure supplement 5, Figure 3–figure supplement 1, Figure 3–figure supplement 2, Figure 4–figure supplement 1, Figure 4–figure supplement 2, Figure 4–figure supplement 4, Figure 4–figure supplement 5, Figure 5–figure supplement 1 and Figure 5–figure supplement 2. These files contain the numerical data used to generate the figures. Figure 4–Source Data 2, Figure 3–figure supplement 1–Source Data 2, and Figure 4–figure supplement 4–Source Data 2 contain the original data of the images in each figure. The lists of yeast strains, plasmids and primers, along with details for plasmid construction, are provided in the Supplementary file.

## Acknowledgments

We thank Shiori Katayama for technical assistance in the investigation of yeast proliferation using FCM; and Jeffrey E. Gerst for providing a plasmid harboring 12 repeats of the MS2 coat protein (MCP) binding sequence. This work was supported in part by KAKENHI Grants-in-Aid for Early-Career Scientists 22K14862 and 25K08894 (to KS), Grant in Aid for Scientific Research (B) 23K25056 and 25K03321 and Grant in Aid for Challenging Research (Exploratory) 16K15089 (to HY) from the Japan Society for the Promotion of Science, ACT-X Grant Number JPMJAX2118 (to KS), GteX Grant Number JPMJGX23B4 (to KS) and SPRING Grant Number JPMJSP2110 (to FS) from the Japan Science and Technology Agency, Noda Institute for Scientific Research Grant (to KS), Joint Research of the Exploratory Research Center on Life and Living Systems (ExCELLS) program No. 26EXC202 (to KS) and by the Program for the Development of Next-generation Leading Scientists with Global Insight (L-INSIGHT), sponsored by the Ministry of Education, Culture, Sports, Science and Technology (MEXT), Japan (to KS).

## Author details

**Fuka Sekioka**

Division of Applied Life Sciences, Graduate School of Agriculture, Kyoto University, Kitashirakawa-Oiwake, Sakyo-ku, Kyoto 606-8502, Japan

**Contribution**: Conceptualization; Methodology; Formal analysis; Investigation; Resources; Data curation; Writing – original draft preparation; Visualization; Funding acquisition

**Competing interests:** No competing interests declared

**Kosuke Shiraishi**

Division of Applied Life Sciences, Graduate School of Agriculture, Kyoto University, Kitashirakawa-Oiwake, Sakyo-ku, Kyoto 606-8502, Japan

**Contribution**: Conceptualization; Methodology; Validation; Formal analysis; Investigation; Resources; Data curation; Writing – original draft preparation; Writing – review & editing; Visualization; Supervision; Project administration; Funding acquisition

**Competing interests:** No competing interests declared

**iD:** 0000-0001-8780-9053

**Miho Akagi**

Division of Applied Life Sciences, Graduate School of Agriculture, Kyoto University, Kitashirakawa-Oiwake, Sakyo-ku, Kyoto 606-8502, Japan

**Contribution**: Methodology; Formal analysis; Investigation; Resources; Data curation; Visualization

**Competing interests:** No competing interests declared

**Akari Habata**

Division of Applied Life Sciences, Graduate School of Agriculture, Kyoto University, Kitashirakawa-Oiwake, Sakyo-ku, Kyoto 606-8502, Japan

**Contribution**: Methodology; Formal analysis; Investigation; Resources; Data curation;

Visualization

**Competing interests:** No competing interests declared

**Yumi Arima**

Division of Applied Life Sciences, Graduate School of Agriculture, Kyoto University, Kitashirakawa-Oiwake, Sakyo-ku, Kyoto 606-8502, Japan

**Contribution**: Formal analysis; Investigation; Resources; Data curation; Visualization

**Competing interests:** No competing interests declared

**Yasuyoshi Sakai**

Department of Bioscience and Biotechnology, Faculty of Bioenvironmental Sciences, Kyoto University of Advanced Science, Kameoka, Kyoto, Japan

**Contribution**: Conceptualization; Methodology; Validation; Writing – review & editing; Supervision

**Competing interests:** No competing interests declared

**Hiroya Yurimoto**

Division of Applied Life Sciences, Graduate School of Agriculture, Kyoto University, Kitashirakawa-Oiwake, Sakyo-ku, Kyoto 606-8502, Japan

**Contribution**: Conceptualization; Methodology; Validation; Writing – review & editing; Supervision; Funding acquisition

**Competing interests:** No competing interests declared

**iD:** 0000-0001-7506-6184

## Funding

Japan Society for the Promotion of Science (22K14862)

- Kosuke Shiraishi

Japan Society for the Promotion of Science (25K08894)

- Kosuke Shiraishi

Japan Society for the Promotion of Science (23K25056)

- Hiroya Yurimoto

Japan Society for the Promotion of Science (25K03321)

- Hiroya Yurimoto

Japan Society for the Promotion of Science (16K15089)

- Hiroya Yurimoto

Japan Science and Technology Agency (JPMJAX2118)

- Kosuke Shiraishi

Japan Science and Technology Agency (JPMJGX23B4)

- Kosuke Shiraishi

Japan Science and Technology Agency (JPMJSP2110)

- Fuka Sekioka

Noda Institute for Scientific Research Grant

- Kosuke Shiraishi

Joint Research of the Exploratory Research Center on Life and Living Systems (ExCELLS) program No. 26EXC202

- Kosuke Shiraishi

Program for the Development of Next-generation Leading Scientists with Global Insight (L-INSIGHT), sponsored by the Ministry of Education, Culture, Sports, Science and Technology (MEXT), Japan

- Kosuke Shiraishi

---

**Figure 1–figure supplement 1.**
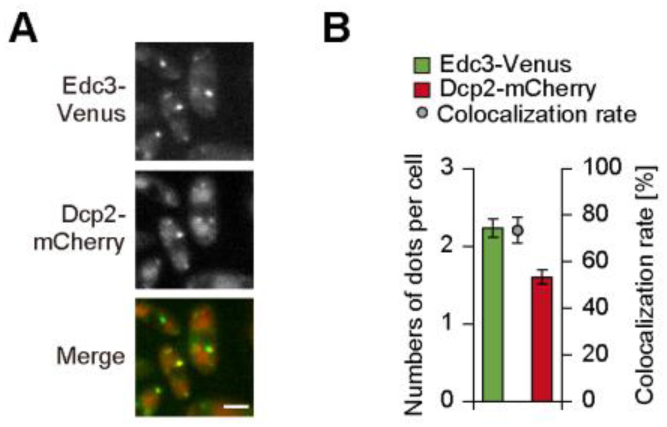
Intracellular dynamics of Edc3-Venus and Dcp2-mCherry. (A) Fluorescence microscopy images of the WT strain expressing Edc3-Venus and Dcp2-mCherry under the control of their native promoters. The strain grown on SD medium was transferred to SM medium and incubated for 3 hours before fluorescence microscopy. Merged images were generated by combining Venus and mCherry fluorescence images. Bar, 2 µm. (B) Quantification of colocalization of Edc3-Venus and Dcp2-mCherry dots detected in (**A**). Cell count analysis was performed on fluorescence microscopy images (n = 30, n: number of cells analyzed; b = 3, b: technical replicates; total = 90 cells). The colocalization rate indicates the percentage of Edc3-Venus dots that colocalized with Dcp2-mCherry dots and is shown as mean ± SE from three technical replicates (n = 30 cells per replicate).

**Figure 1–figure supplement 2.**
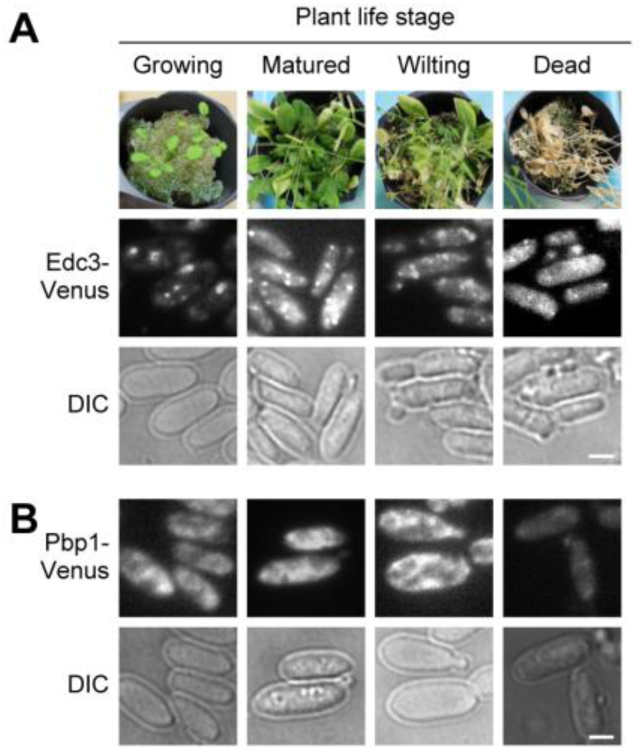
Intracellular localization of Edc3-Venus and Pbp1-Venus on *A. thaliana* leaves. (**A-B**) Investigation of the P-body and SG formation in the WT strains on *Arabidopsis* leaves. Edc3-Venus fusion protein was expressed under the control of the native *EDC3* promoter as a P-body marker (**A**), whereas Pbp1-Venus fusion protein expressed under the control of the native *PBP1* promoter was used as an SG marker (**B**). The strains were spotted on growing, matured, wilting or dead leaves, collected from the leaves 4 hours after inoculation and observed by a fluorescent microscope. Bar, 2 µm. DIC: differential interference contrast.

**Figure 2–figure supplement 1.**
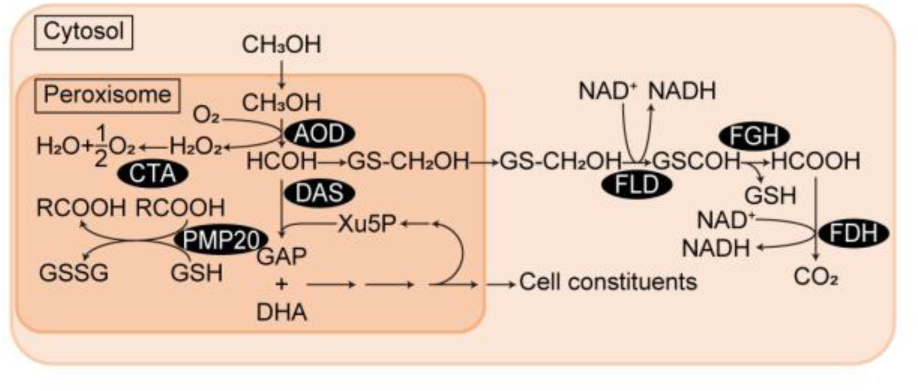
Diagram of methanol metabolism in yeast. Enzymes: AOD, alcohol oxidase; DAS, dihydroxyacetone synthase; FDH, formate dehydrogenase; FGH, *S*-formylglutathione hydrolase; FLD, formaldehyde dehydrogenase; CTA, catalase; PMP20, peroxisome membrane protein with glutathione peroxidase activity. Abbreviations: DHA, dihydroxyacetone; GAP, glyceraldehyde 3-phosphate; GS-CH2OH, S-hydroxymethyl glutathione; GS-CHO, S-formylglutathione; GSH, reduced form of glutathione; GSSG, oxidized form of glutathione; RCOOOH, alkyl hydroperoxide.

**Figure 2–figure supplement 2.**
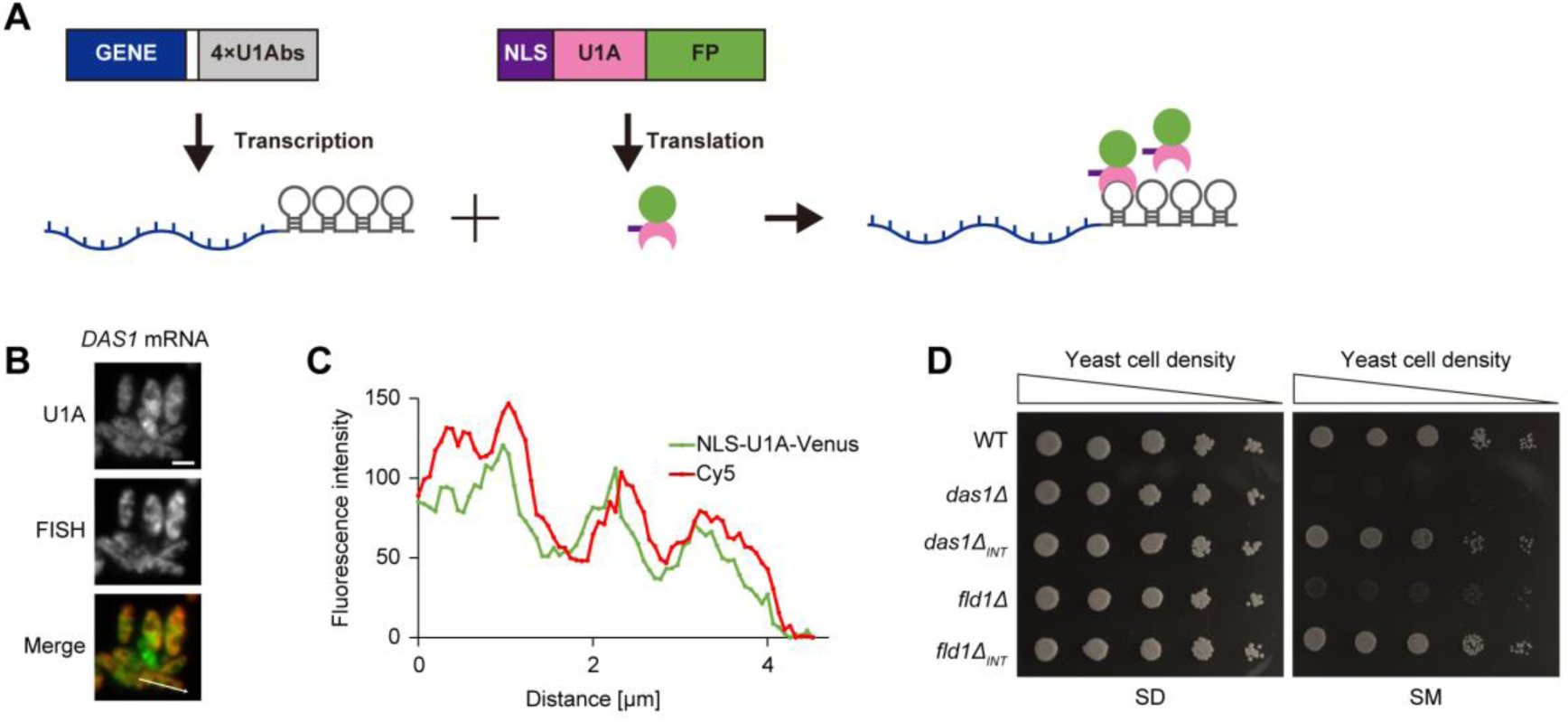
Integration of the U1Abs did not alter protein function. (**A**) Schematic overview for visualizing methanol-induced mRNAs (mimRNAs) *in vivo*. GENE, DNA sequence encoding a target mRNA; White box, UTR; 4xU1Abs, 4 x U1A binding sequence; NLS, nuclear localization sequence; U1A, U1A coding sequence; FP, fluorescent protein coding sequence. Yeast cells were transformed with vectors expressing a fusion of U1Abs with a gene sequence encoding the target mRNA sequence (GENE-4xU1Abs) and a fluorescent protein combined with NLS-U1A (NLS-U1A-FP). GENE-4xU1Abs was expressed under its own promoter, whereas NLS-U1A-FP was expressed under the *TDH3* promoter. (**B**) Fluorescent microscopic images of *DAS1* mRNA visualized by the U1A-based RNA system and fluorescence *in situ* hybridization (FISH) method. FISH was performed for visualizing the canonical *DAS1* gene (methodological details are described in Materials and Methods section). Strain DAS1m-V grown on an SD medium was transferred to an SM medium and incubated for 3 hours. Merged images were generated by combining Venus and cy5 fluorescence images. Bar: 2 µm. (**C**) Colocalization of the *DAS1* mRNA detected by the U1A-based RNA system and FISH method in (**B**). mRNA localization was visualized by FISH using Cy5-labeled probes and by the U1A-based RNA system using Venus fluorescence. Fluorescence intensity was measured along a defined line across the cells. Fluorescence intensity (Y-axis) was plotted against distance (X-axis) to show the spatial distribution of mRNA signals. (**D**) Growth assay of *C. boidinii* strains on SD or SM agar plates. The yeast strains were grown to early log phase, adjusted to OD_610_=1, and 3μL of tenfold serial dilutions were streaked onto SD or SM plates. The *das1Δ_INT_* strain expresses 4xU1Abs-tagged native *DAS1* gene under its native promoter and the *fld1Δ_INT_* strain expresses 4xU1Abs-tagged native *FLD1* gene under its native promoter.

**Figure 2–figure supplement 3.**
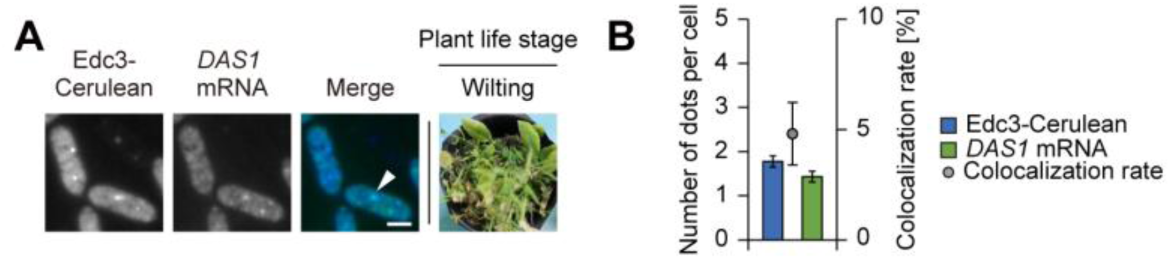
Intracellular dynamics of P-bodies and mimRNAs on wilting *A. thaliana* leaves. (**A**) Fluorescence microscopy images of strain Edc3-C/DAS1m-V on wilting *Arabidopsis* leaves. The strain, spotted on leaves, was collected from the leaf surface 4 hours after inoculation for observation. Merged images were generated by combining Venus and Cerulean fluorescence images. White arrows indicate colocalization of Edc3-Cerulean and Venus-tagged *DAS1* mRNA dots. Bar, 2 µm. (**B**) Quantification of dot formation of Edc3-Cerulean and Venus-tagged *DAS1* mRNA and their colocalization detected in (**A**). Cell count analysis was performed on fluorescence microscopy images (n = 30, n: number of cells analyzed; b = 3, b: technical replicates; total = 90 cells). The number of dots is presented as mean ± SE of 90 cells and colocalization rates are shown as mean ± SE from three technical replicates (n = 30 cells per replicate).

**Figure 2–figure supplement 4.**
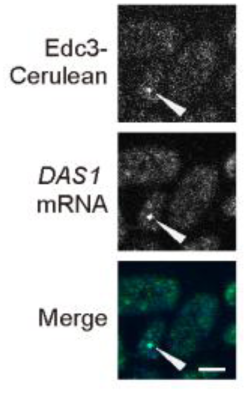
Confocal microscopy analysis of Edc3-Cerulean and Venus-tagged *DAS1* mRNA. Confocal microscopy images of the WT strain expressing Edc3-Cerulean and Venus-tagged *DAS1* mRNA (strain Edc3-C/DAS1m-V). The strain grown on SD medium was transferred to SM medium and incubated for 3 hours before observation. Images of a single optical section are shown. Merged images were generated by combining Venus and Cerulean fluorescence images. White arrows indicate overlapping Edc3-Cerulean and Venus-tagged *DAS1* mRNA dots. Bar, 2 µm.

**Figure 2–figure supplement 5.**
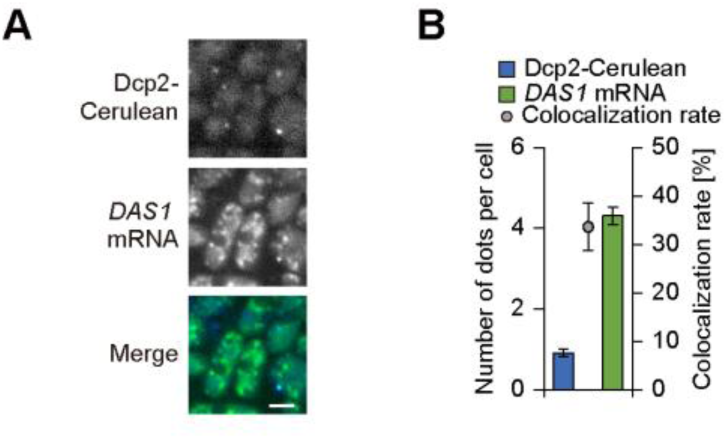
Intracellular dynamics of P-bodies labeled with Dcp2-Cerulean and Venus-tagged *DAS1* mRNA. (**A**) Fluorescence microscopy images of the WT strain expressing Dcp2-Cerulean and Venus-tagged *DAS1* mRNA. Dcp2-Cerulean was expressed under the control of the native *DCP2* promoter. The strain grown on SD medium was transferred to SM medium and incubated for 3 hours before fluorescence microscopy. Merged images were generated by combining Venus and Cerulean fluorescence images. Bar, 2 µm. (**B**) Quantification of dot formation of Dcp2-Cerulean and Venus-tagged *DAS1* mRNA and their colocalization detected in (**A**). Cell count analysis was performed on fluorescence microscopy images (n = 30, n: number of cells analyzed; b = 3, b: technical replicates; total = 90 cells). The number of dots is presented as mean ± SE of 90 cells, and co-localization rates are shown as mean ± SE from three technical replicates (n = 30 cells per replicate).

**Figure 3–figure supplement 1.**
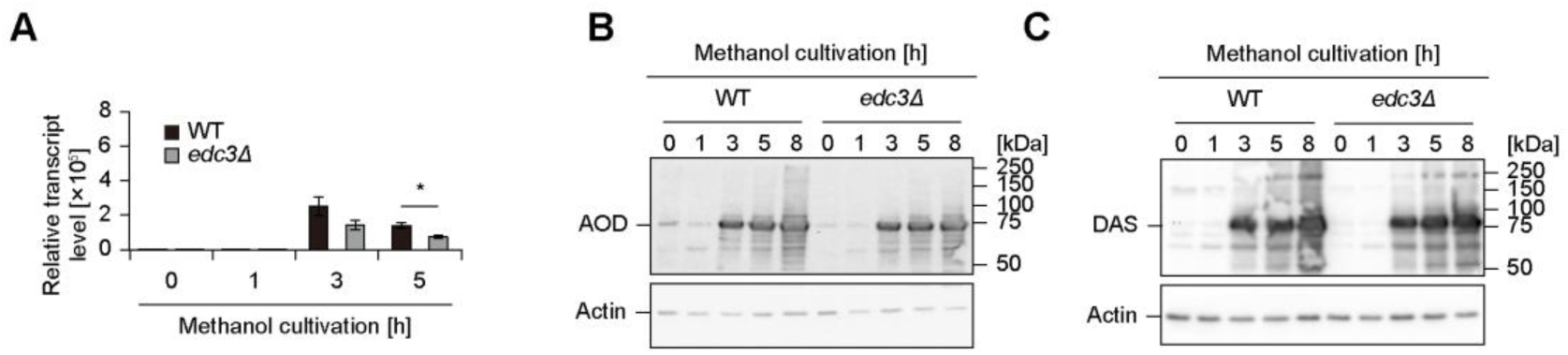
Effect of the *EDC3* gene deletion on methanol metabolism. (**A**) Relative transcript level of *AOD1* mRNA in the WT and *edc3Δ* strains on SM medium. Values are indicated as mean ± SE of three biological replicates. Statistical significances between the WT and *edc3Δ* strains were assessed. Asterisks indicate statistical significance: * *p* < 0.05. (**B-C**) Immunoblot analysis of AOD (**B**) and DAS (**C**) in the WT and *edc3Δ* strains during methanol induction. The expected molecular weights of AOD and DAS are 74.1 kDa and 78.2 kDa, respectively.

**Figure 3–figure supplement 2.**
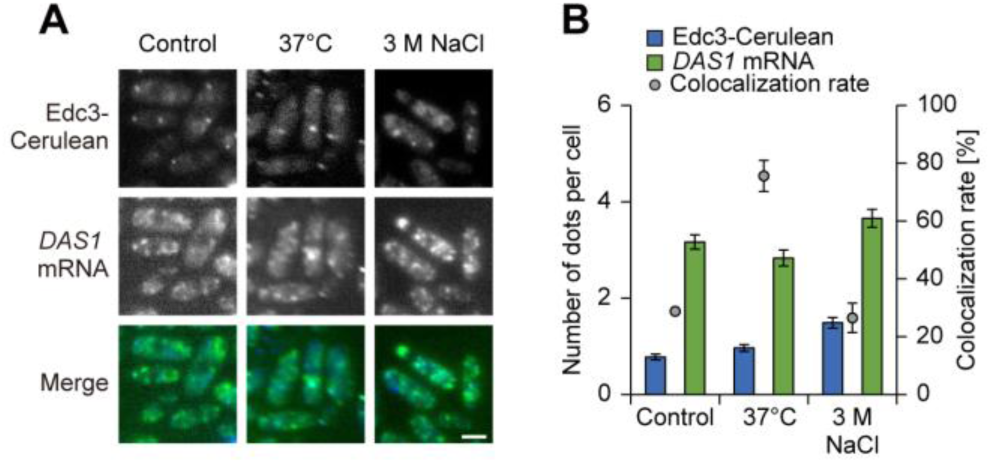
Intracellular dynamics of P-bodies and *DAS1* mRNA under stress conditions. (**A**) Fluorescence microscope images of strain Edc3-C/DAS1m-V during heat and salt stress. The strain was exposed to heat shock at 37 degrees or salt stress with 3 M NaCl for 1 hour before microscopic observation. Merged images were generated by combining Venus and Cerulean fluorescence images. Bar, 2 µm. (**B**) Quantification of dot formation of Edc3-Cerulean and Venus-tagged *DAS1* mRNA and their colocalization detected in (**A**). Cell count analysis was performed on fluorescence microscopy images (n = 30, n: number of cells analyzed; b = 3, b: technical replicates; total = 90 cells). The number of dots is presented as mean ± SE of 90 cells and colocalization rates are shown as mean ± SE from three technical replicates (n = 30 cells per replicate).

**Figure 4–figure supplement 1.**
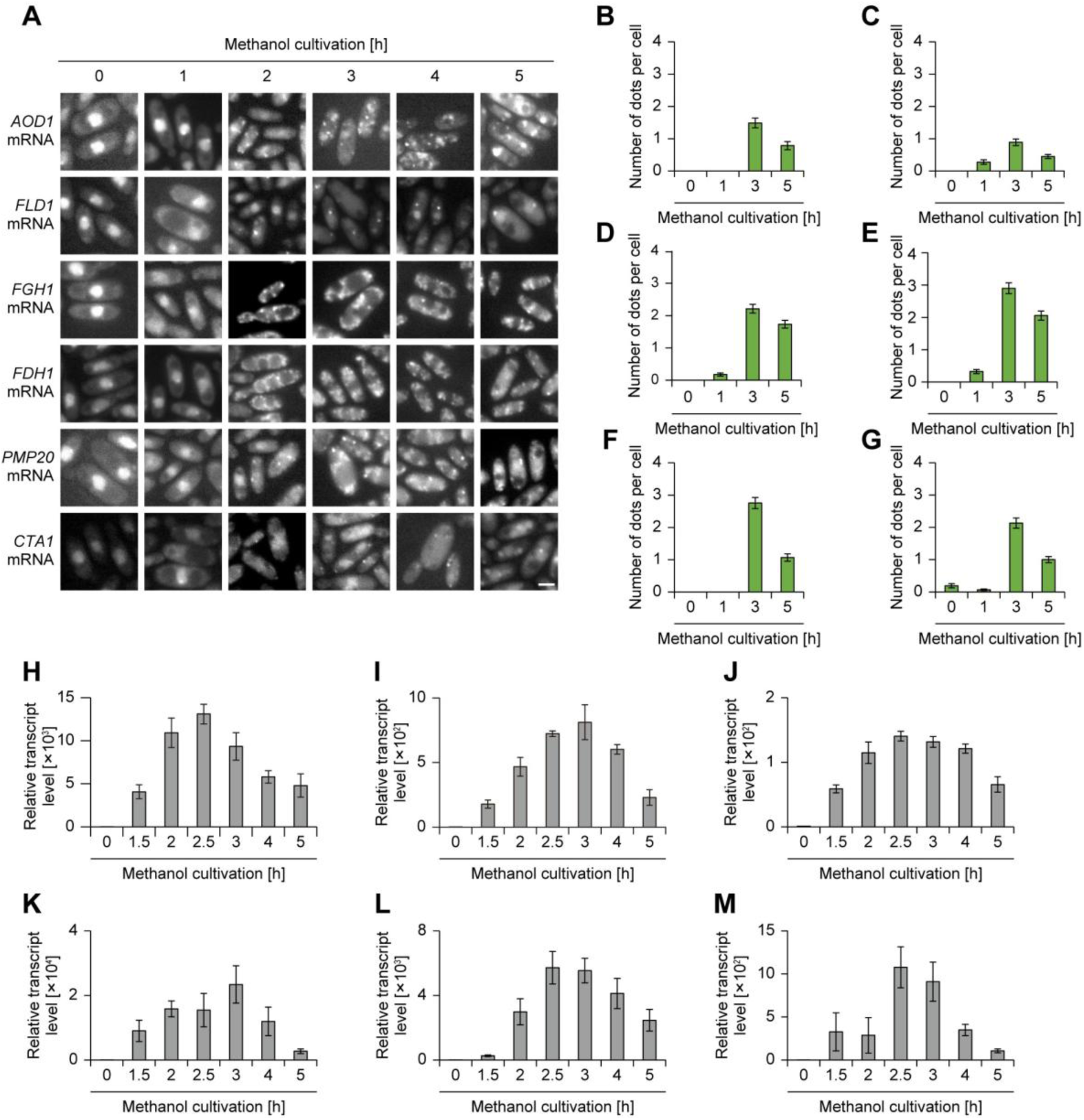
Granule formation and gene expression of various mimRNAs during methanol induction. (**A**) Time-course analysis of the intracellular localization of mimRNAs during methanol induction*. AOD1*, *FLD1*, *FGH1*, *FDH1*, *PMP20* and *CTA1* mRNA were visualized. The strains grown on an SD medium were transferred to SM medium and incubated for up to 5 hours and collected at indicated time points. Bar, 2 µm. (**B-G**) Quantification of dot formation of mimRNAs detected in (**A**). (**B**) *AOD1*, (**C**) *FLD1*, (**D**) *FGH1*, (**E**) *FDH1*, (**F**) *PMP20* and (**G**) *CTA1* mRNA. At least 30 cells were analyzed per experiment. Cell count analysis was performed on fluorescence microscopy images (n = 30, n: number of cells analyzed; b = 3, b: technical replicates; total = 90 cells). The number of dots is presented as the mean ± SE of 90 cells. (**H-M**) Relative transcript level of mimRNAs in the WT strain during methanol induction. (**H**) *AOD1*, (**I**) *FLD1*, (**J**) *FGH1*, (**K**) *FDH1*, (**L**) *PMP20* and (**M**) *CTA1* mRNA. Values are indicated as mean ± SE of three independent experiments.

**Figure 4–figure supplement 2.**
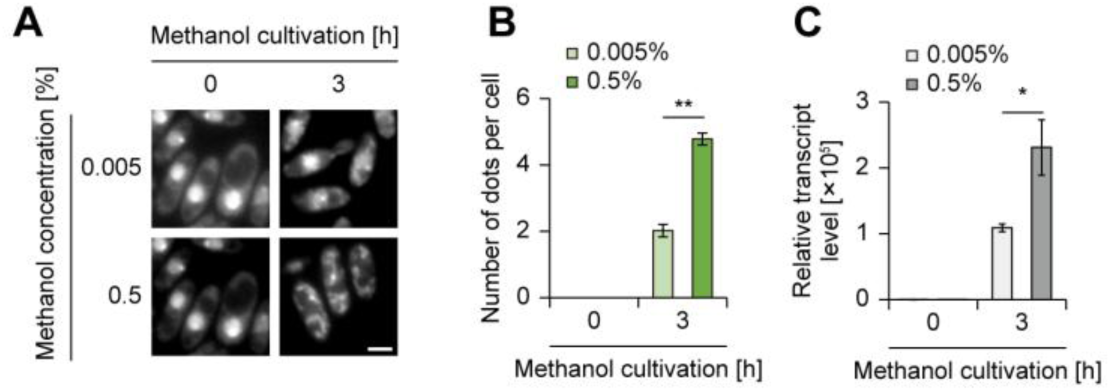
Regulation of mimRNA granule formation at low methanol concentration. (**A**) Fluorescence microscopy images of the strain visualizing Venus-tagged *DAS1* mRNA on SM medium. The strain grown on an SD medium was transferred to SM medium and incubated for 3 hours and collected at indicated time points. Methanol concentration was either 0.005% or 0.5%. Bar, 2 µm. (**B**) Quantification of mimRNA granule formation detected in (**A**). Cell count analysis was performed on fluorescence microscopy images (n = 30, n: number of cells analyzed; b = 3, b: technical replicates; total = 90 cells). The number of dots is presented as mean ± SE of 90 cells. Asterisks indicate statistical significance: ** *p* < 0.01. (**C**) Relative transcript level of *DAS1* mRNA in the WT strain during methanol induction. Methanol concentration was either 0.005% or 0.5%. Values are indicated as mean ± SE of three biological replicates. Asterisks indicate statistical significance: * *p* < 0.05.

**Figure 4–figure supplement 3.**
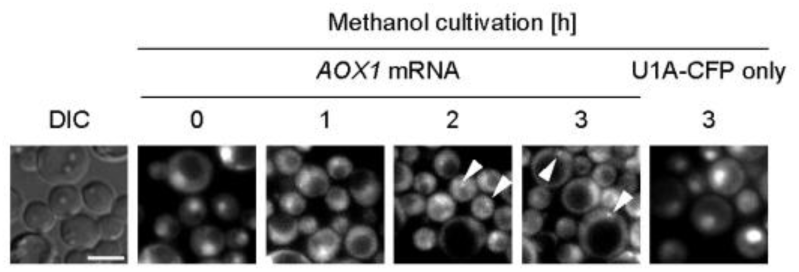
Visualization of methanol-induced *KpAOX1* mRNA in *K. phaffii*. Fluorescent microscopic images of the *K. phaffii* strain expressing U1Abs-tagged *KpAOX1* mRNA, and the NLS- and CFP-tagged U1A protein. U1Abs; U1A protein binding sequence, NLS; nuclear localization sequence. The *K. phaffii* strain expressing only U1A-CFP (referred to as U1A-CFP only) was used as a control in which the CFP fluorescence remained in the nucleus. White arrows indicate the dots of *KpAOX1* mRNA. Bar, 2 µm. DIC: differential interference contrast.

**Figure 4–figure supplement 4.**
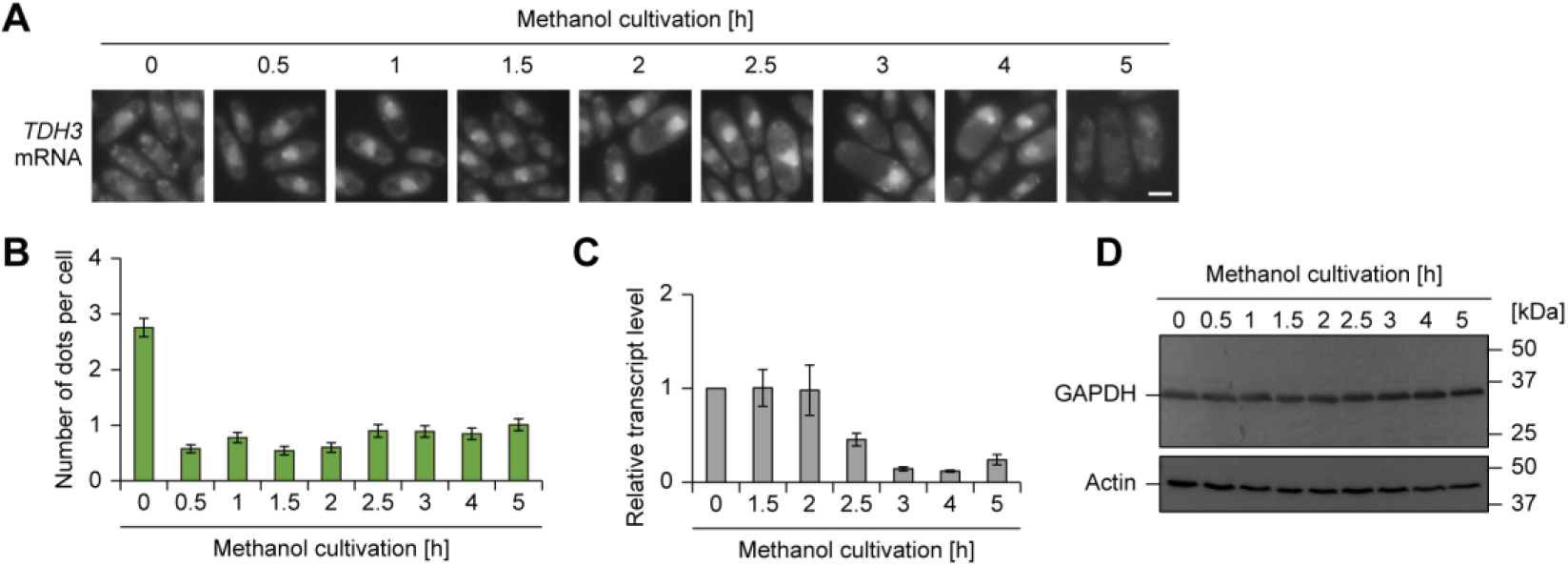
Regulation of dot formation of *TDH3* mRNA. (**A**) Time-course analysis of the intracellular localization of Venus-tagged *TDH3* mRNA during methanol induction. The strain grown on an SD medium was transferred to SM medium and incubated for up to 5 hours. Bar, 2 µm. (**B**) Quantification of dot formation of Venus-tagged *TDH3* mRNA detected in (**A**). Cell count analysis was performed on fluorescence microscopy images (n = 30, n: number of cells analyzed; b = 3, b: technical replicates; total = 90 cells). The number of dots is presented as mean ± SE of 90 cells. (**C**) Relative transcript level of *TDH3* mRNA in the WT strain during methanol induction. Values are indicated as mean ± SE of three biological replicates. (**D**) Immunoblot analysis of GAPDH in the WT strain during methanol induction. The expected molecular weight of GAPDH is 35.5 kDa.

**Figure 4–figure supplement 5.**
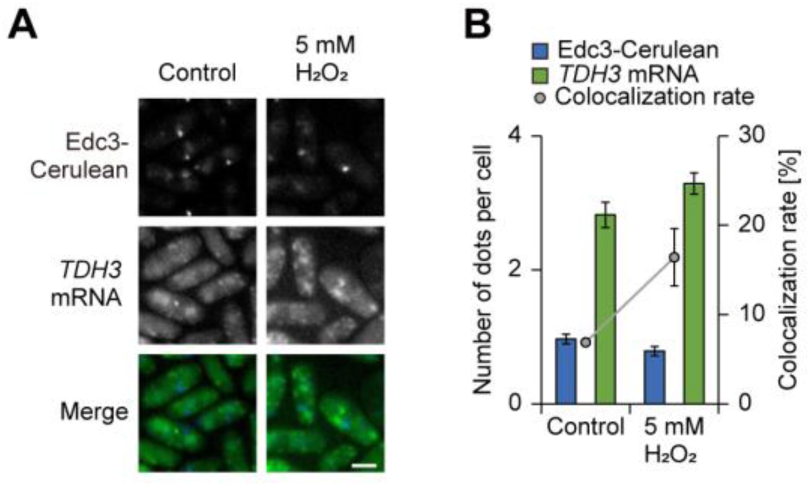
Intracellular dynamics of P-bodies and *TDH3* mRNA under stress conditions. (**A**) Fluorescence microscopy images of strain Edc3-C/TDH3m-V during oxidative stress. The strain incubated on an SM medium was treated with 5 mM H_2_O_2_ for 1 hour before microscopic observation. Merged images were generated by combining Venus and Cerulean fluorescence images. Bar, 2 µm. (**B**) Quantification of dot formation of Edc3-Cerulean and Venus-tagged *TDH3* mRNA and their colocalization detected in (**A**). Cell count analysis was performed on fluorescence microscopy images (n = 30, n: number of cells analyzed; b = 3, b: technical replicates; total = 90 cells). The number of dots is presented as mean ± SE of 90 cells and colocalization rates are shown as mean ± SE from three technical replicates (n = 30 cells per replicate).

**Figure 5–figure supplement 1.**
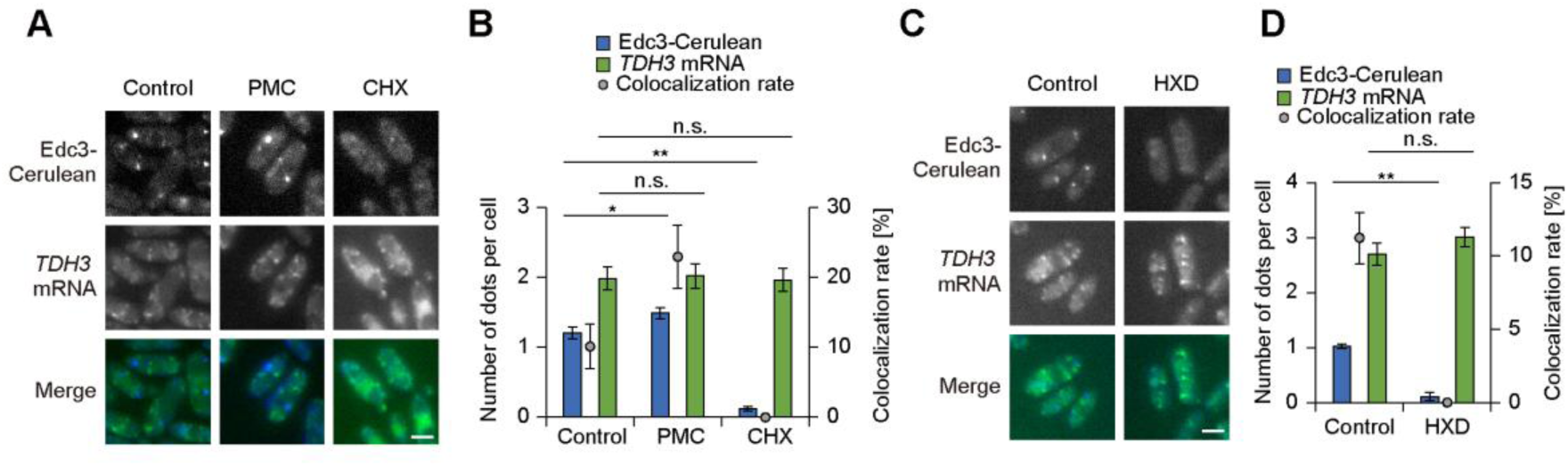
Effects of protein synthesis inhibitors and 1,6-hexanediol on the dot formation of *TDH3* mRNA. (**A**) Fluorescence microscopy images of strain Edc3-C/TDH3m-V after treatment with protein synthesis inhibitors. The strain incubated on SM medium was treated with either 0.1 mg/mL puromycin (PMC) or 0.1 mg/mL cycloheximide (CHX) before microscopic observation. Merged images were generated by combining Venus and Cerulean fluorescence images. Bar, 2 µm. (**B**) Quantification of dot formation of Edc3-Cerulean and Venus-tagged *TDH3* mRNA and their colocalization detected in (**A**). Cell count analysis was performed on fluorescence microscopy images (n = 30, n: number of cells analyzed; t = 3, t: technical replicates; total = 90 cells). The number of dots is presented as mean ± SE of 90 cells, and colocalization rates are shown as mean ± SE from three technical replicates (n = 30 cells per replicate). Asterisks indicate statistical significance: * *p* < 0.05; ** *p* < 0.01; n.s. means not significant. (**C**) Fluorescence microscopy images of strain Edc3-C/TDH3m-V after treatment with 1,6-hexanediol (HXD). The strain incubated on SM medium was treated with 10% HXD before microscopic observation. Merged images were generated by combining Venus and Cerulean fluorescence images. Bar, 2 µm. (**D**) Quantification of dot formation of Edc3-Cerulean and Venus-tagged *TDH3* mRNA and their colocalization detected in (**C**). Cell count analysis was performed, as described in (B). Asterisks indicate statistical significance: ** *p* < 0.01; n.s. means not significant.

**Figure 5–figure supplement 2.**
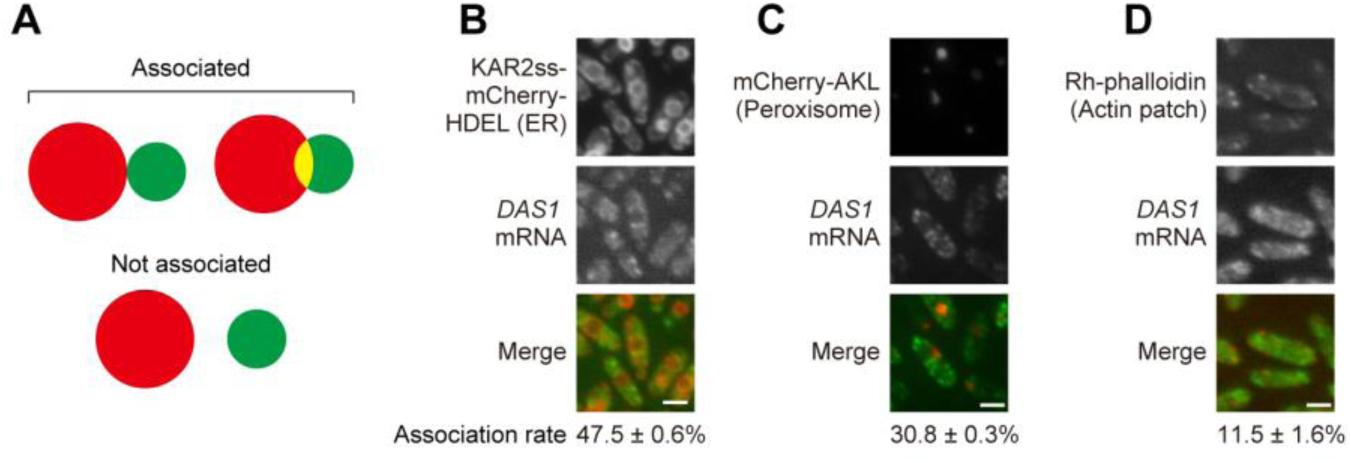
Association of mimRNA granules with other organelles and structures. (**A**) Schematic image showing the criteria for association or non-association of *DAS1* mRNA with cellular organelles. *DAS1* mRNA granules were defined as associated when their fluorescence overlapped with or was in contact with the organelle signal. (**B-D**) Association of *DAS1* mRNA granules with the endoplasmic reticulum (ER) (**B**), peroxisomes (**C**) and actin patches (**D**). ER was visualized by mCherry tagged with KAR2 sequence signal at its N terminus and HDEL amino acid sequence at its C terminus. Peroxisomes were visualized by mCherry tagged with AKL amino acid sequence at its C terminus for peroxisome targeting signal. Actin patches were stained with rhodamine phalloidin. Cells grown on an SD medium were transferred to an SM medium and incubated for 3 hours. The association rate indicates the percentage of Venus-tagged *DAS1* mRNA granules that are associated with the ER (**B**), peroxisomes (**C**) or actin patches (**D**). Cell count analysis was performed on fluorescence microscopy images (n = 30, n: number of cells analyzed; b = 3, b: technical replicates; total = 90 cells). Association rates are shown as mean ± SE from three technical replicates (n = 30 cells per replicate). Bar, 2 µm.

## Legends to videos

**Dynamic properties of Edc3-Cerulean and Venus-tagged *DAS1* mRNA dots during growth on methanol.**

Time-lapse images of Edc3-Cerulean and Venus-tagged *DAS1* mRNA dots (see also Figure 5A) were acquired every 28 seconds for 9 minutes 52 seconds by fluorescence microscopy and compiled into a video at 2 frames per second (fps). Bar, 2 µm.

