## Supplementary information for "The P-body component Edc3 is required for yeast proliferation in the phyllosphere"

**Extended Materials and Methods**

**Construction of yeast strains and plasmids**

Yeast strains, primers, and plasmids used in this study are listed in Table S1, Table S2 and Table S3, respectively.

**Construction of plasmids for *C. boidinii***

A deletion cassette for the *EDC3* gene was constructed as follows: The primer set Fw_EDC3_UP / Rv_EDC3_UP was used to amplify a 1.0-kb fragment from *C. boidinii* genomic DNA. The PCR product was fused with the 7.6-kb KpnI-digested SK+SPR vector using the In-Fusion HD® Cloning Kit (TaKaRa Bio Inc., Shiga, Japan), yielding SK+SPR harboring the upstream region of *EDC3*. Similarly, the primer set Fw_EDC3_DOWN / Rv_EDC3_DOWN was used to amplify a 1.0-kb downstream fragment, which was fused with the 8.6-kb SacI-digested SK+SPR (containing the upstream region of *EDC3*) using the same kit, resulting in the *EDC3* disruption vector pPB01.

To visualize intracellular organelles, vectors pPB02–pPB09 and pPB26–pPB27 were generated as follows:

- pPB02 and pPB03: The *DCP2* promoter and ORF (without the stop codon) were amplified using primers Fw_pEX-DCP2_inf / Rv_pEX-DCP2_inf and inserted into pACTV linearized by inverse PCR with primers Fw_pEX-V_inv / Rv_pEX-V_inv, yielding pPB02. Using the same strategy with primers Fw_pEX-Pbp1_inf / Rv_pEX-Pbp1_inf, pPB03 was obtained, containing the *PBP1* promoter and ORF (without the stop codon).
- pPB04 and pPB06: The *CERULEAN* gene was amplified from pEX-A2J2-Cerulean with primers Fw_Cerulean_SalI / Rv_Cerulean_PstI, digested with SalI/PstI, and ligated into the 6.4-kb SalI/PstI-digested pACTL-Venus vector using the Ligation Convenience Kit (Nippongene, Tokyo, Japan), generating pPB04. pPB04 was then linearized by inverse PCR (Fw_pEXL-Cerulean_inv / Rv_pEXL-Cerulean_inv) and fused with the *EDC3* promoter and ORF (without the stop codon) amplified from pHC404 (Fw_pEX-EDC3_inf / Rv_pEX-EDC3_inf), yielding pPB06.
- pPB05, pPB07, pPB08, and pPB09: The *mCHERRY* gene was amplified using primers Fw_mCherry_SalI / Rv_mCherry_PstI and ligated as above to yield pPB05. This plasmid was linearized by inverse PCR (Fw_mCherry-AKL_inv / Rv_mCherry-AKL_inv) and self-ligated with T4 Polynucleotide Kinase (New England Biolabs, Ipswich, MA, USA), yielding pPB07. For ER-targeted expression, pPB05 was linearized by inverse PCR (Fw_mChe_HDEL_inv / Rv_mChe_HDEL_inv) and self-ligated to generate pPB08. pPB08

was then PCR-amplified with primers Fw_mChe_ATG_inv / Rv_mChe_ATG_inv and fused with the *KAR2* signal sequence (KAR2ss) amplified from *C. boidinii* genomic DNA (Fw_KAR2ss_inf / Rv_KAR2ss_inf), generating pPB09.

- pPB26: The *DCP2* promoter and ORF (without the stop codon) were amplified using primers Fw_pEX-DCP2_inf / Rv_pEX-DCP2_inf and inserted into pPB04 linearized by inverse PCR with primers Fw_pEXL-Cerulean_inv / Rv_pEXL-Cerulean_inv, resulting in pPB26.
- pPB27: pPB02 was linearized by inverse PCR (Fw_pEX-Dcp2_inv / Rv_pEX-Dcp2_inv) and fused with the mCherry amplified from pSPM001 (Fw_mCherry_inf / Rv_mCherry_inf), resulting in pPB27.

For U1A-based mRNA visualization, plasmids pPB10–pPB23 were generated as follows:

- pPB10–pPB13: The *TDH3* promoter region was amplified with primers Fw_pTDH3_inf / Rv_pTDH3_inf and ligated into the 7.5-kb XhoI/SalI-digested pACTL-Venus vector to yield pPB10. pPB10 was linearized by inverse PCR (Fw_pTDH3L-Venus_inv / Rv_pTDH3L-Venus_inv) and fused with the U1A ORF (without stop codon) amplified from pUCFa-U1A_1-102_Cb (Fw_U1A_inf / Rv_U1A_inf), yielding pPB11.
  To introduce an NLS, pPB11 was linearized by inverse PCR (Fw_pTDH3L-NLS-U1A-Venus_inv / Rv_pTDH3L-NLS-U1A-Venus_inv) and self-ligated with T4 Polynucleotide Kinase, generating pPB12. To replace the *LEU2* marker with *ZEO$^r$*, pPB12 was linearized by inverse PCR using primers Fw_pTDH3-NLS-U1A-Venus_inv / Rv_pTDH3-NLS-U1A-Venus_inv. This fragment was fused to a *TEF1-ZEO$^r$-CYC* cassette amplified from pREMI-Zc using primers Fw_ZeoR_inf / Rv_ZeoR_inf, resulting in pPB13.
- pPB14–pPB23: The plasmid pPB14 harboring the *AOD1* promoter and ORF region along with four tandem U1A binding sequences (4×U1Abs) at the 3′ UTR was constructed as follows: The *AOD1* ORF region was amplified using primers Fw_AOD1_inf / Rv_AOD1_inf with genomic DNA of *C. boidinii* as the template. The obtained fragment was inserted into pNOTel linearized by inverse PCR using primers Fw_pNOTel_inv / Rv_pNOTel_inv. The resulting vector was then linearized by PCR using primers Fw_AOD1_inv / Rv_AOD1_inv and fused with the 4×U1A binding site amplified with primers Fw_4xU1Abs / Rv_4xU1Abs using pSYN5908-1 as the template, resulting in pPB14. Vectors pPB15–pPB23 were constructed by replacing the *AOD1* promoter and ORF region with those of *DAS1*, *FLD1*, *FGH1*, *FDH1*, *PMP20*, *CTA1*, *TDH3*, *ADH1* and *ACT1*, respectively. These promoter and ORF regions were amplified from genomic DNA using the following primer sets: Fw_PDAS1_inf / Rv_PDAS1_inf, Fw_PFLD1_inf /

Rv_PFLD1_inf, Fw_PFGH1_inf / Rv_PFGH1_inf, Fw_PFDH1_inf / Rv_PFDH1_inf, Fw_PPMP20_inf / Rv_PPMP20_inf, Fw_PCTA1_inf / Rv_PCTA1_inf, Fw_PTDH3_inf / Rv_PTDH3_inf, Fw_PADH1_inf / Rv_PADH1_inf, and Fw_PACT1_inf / Rv_PACT1_inf, respectively. These amplified DNAs were then fused to the vector linearized by PCR with primers Fw_3′ U1Abs_inv / Rv_3′ U1Abs_inv using pPB14 as the template.

For MS2-based mRNA visualization, plasmids pPB24–pPB25 were generated as follows:

- pPB24 was constructed using pPB13 as a template in three steps: (i) the U1A coding region was replaced with the MS2 coat protein coding region. pPB13 was linearized by inverse PCR using primers Fw_pTDH3Z-NLS-Venus_inv / Rv_pTDH3Z-NLS-Venus_inv, and the resulting fragment was fused to the MS2 coding region amplified with primers Fw_MS2_inf / Rv_MS2_inf using pEX-A2J2-MS2 as the template. (ii) The *VENUS* coding region was replaced with *CERULEAN*. The vector was linearized by inverse PCR with primers Fw_pTDH3Z-NLS-MS2_inv / Rv_pTDH3Z-NLS-MS2_inv and fused to the *CERULEAN* coding region amplified with primers Fw_Cerulean_inf / Rv_Cerulean_inf using pPB04 as the template. (iii) The $ZEO^r$ gene was replaced with $BSD^r$. The obtained vector was linearized by PCR using primers Fw_pTDH3-NLS-U1A-Cerulean_inv / Rv_pTDH3-NLS-U1A-Cerulean_inv and fused to the $BSD^r$ coding region amplified with primers Fw_BsdR_inf / Rv_BsdR_inf using pEX-A2J2-Bsd$^r$ as the template, resulting in pPB24.
- pPB25 was constructed as follows: First, the *URA3* selection marker of pPB15 was replaced with *LEU2*. pPB15 was linearized by PCR using primers Fw_pEX-DAS1-4xU1Abs_inv / Rv_pEX-DAS1-4xU1Abs_inv and fused with the *LEU2* coding region amplified from pACTL-Venus using primers Fw_LEU2_inf / Rv_LEU2_inf. The resulting vector was then linearized by PCR using primers Fw_pEX_DAS1_inv_forMS2bs / Rv_pEX_DAS1_inv_forMS2bs and fused to the 12×MS2-CP binding site array amplified with primers Fw_MS2L12_inf / Rv_MS2L12_inf using pLOX_HIS_MS2Lx12 as the template, resulting in pPB25.

**Construction of the *edc3Δ* strain**

A deletion cassette for the *EDC3* gene was amplified using primers Fw_EDC3_UP / Rv_EDC3_DOWN with pPB01 as the template. The PCR product was transformed into *C. boidinii* TK62 using a modified version of the lithium acetate method (Sakai & Tani, 1992b). Proper gene disruption was confirmed by colony PCR. The constructed *edc3Δ* strain (yPB101) was converted to uracil auxotrophy by 5-FOA selection, yielding the *edc3Δ ura3* strain (yPB102). Restoration of the *URA3* marker gene was confirmed by PCR analysis.

**Construction of plasmids for *K. phaffii***

To visualize *KpAOX1* mRNA using the U1A-based system, vectors pPBK01–pPBK02 were generated as follows:

- pPBK01 was constructed as follows: The host vector harboring the *GAP* promoter was prepared by linearizing pNT2102 with primers Fw_pIB2_inv / Rv_pIB2_inv and fused to the *CFP* coding region amplified with primers Fw_KpCFP_inf / Rv_KpCFP_inf using pNT2103 as the template. The obtained plasmid was further linearized by PCR using primers Fw_pIB2-CFP_inv / Rv_pIB2-CFP_inv and fused to the NLS-U1A coding region amplified with primers Fw_NLS-U1A_inf / Rv_NLS-U1A_inf using pPB12 as the template, resulting in pPBK01.
- pPBK02 was constructed as follows: The host vector was prepared by linearizing pNT204 with primers Fw_pIB1-Arg_inv / Rv_pIB1-Arg_inv and fused to the *KpAOX1* promoter and ORF amplified from genomic DNA using primers Fw_pEX-KpAOX1_inf / Rv_pEX-KpAOX1_inf. The obtained vector was further linearized using primers Fw_pIB1-Arg-KpAOX1_inv / Rv_pIB1-Arg-KpAOX1_inv and fused to the U1Abs coding region amplified with primers Fw_CbU1A_inf / Rv_CbU1A_inf using pPB14 as the template, resulting in pPBK02.

**Table S1. List of yeast strains used in this study**

| Designation | Description | Genotype | References |
|---|---|---|---|
| *C. boidinii* | | | |
| AOU1 | Wild type | Wild type | (Tani et al., 1985) |
| TK62 | *ura3Δ* | *ura3* | (Sakai et al., 1991) |
| BUL | *ura3Δ leu2Δ* | *ura3 leu2* | (Sakai & Tani, 1992a) |
| yPB101 | *edc3Δ* | TK62 *edc3Δ*::*URA3* | This study |
| yPB102 | *edc3Δ ura3Δ* | yPB101 *ura3* | This study |
| yPB103 | pACT1-Venus, TK62 | TK62 pACTV::*URA3* | (Kawaguchi et al., 2011) |
| yPB104 | pACT1-Venus, *edc3Δ* | *edc3Δ* pACTV::*URA3* | This study |
| yPB105 | Edc3-Venus, TK62 | TK62 *ura3*::($P_{EDC3}$-EDC3-Venus, *URA3*) | This study |
| yPB106 | Dcp2-Venus, TK62 | TK62 *ura3*::($P_{DCP2}$-DCP2-Venus, *URA3*) | This study |
| yPB107 | Pbp1-Venus, TK62 | TK62 *ura3*::($P_{PBP1}$-PBP1-Venus, *URA3*) | This study |
| yPB108 | Dcp2-Venus, *edc3Δ* | yPB102 *ura3*::($P_{DCP2}$-DCP2-Venus, *URA3*) | This study |
| yPB109 | Edc3-Venus, *edc3Δ* | yPB102 *ura3*::($P_{EDC3}$-EDC3-Venus, *URA3*) | This study |
| yPB110 | *AOD1* mRNA visualization by U1A-based RNA system, BUL | BUL *ura3*::($P_{AOD1}$-AOD1-4xU1Abs, *URA3*) *leu2*::($P_{TDH3}$-NLS-U1A-Venus, *LEU2*) | This study |
| yPB111 | *DAS1* mRNA visualization by U1A-based RNA system, BUL | BUL *ura3*::($P_{DAS1}$-DAS1-4xU1Abs, *URA3*) *leu2*::($P_{TDH3}$-NLS-U1A-Venus, *LEU2*) | This study |
| yPB112 | *DAS1* mRNA visualization by U1A-based RNA system, *edc3Δ* | yPB102 *ura3*::($P_{DAS1}$-DAS1-4xU1Abs, *URA3*) *leu2*::($P_{TDH3}$-NLS-U1A-Venus, *LEU2*) | This study |

| | | | |
|---|---|---|---|
| yPB113 | *FLD1* mRNA visualization by U1A-based RNA system, BUL | BUL *ura3*::($P_{FLD1}$-FLD1-4xU1Abs, *URA3*) *leu2*::($P_{TDH3}$-NLS-U1A-Venus, *LEU2*) | This study |
| yPB114 | *FGH1* mRNA visualization by U1A-based RNA system, BUL | BUL *ura3*::($P_{FGH1}$-FGH1-4xU1Abs, *URA3*) *leu2*::($P_{TDH3}$-NLS-U1A-Venus, *LEU2*) | This study |
| yPB115 | *FDH1* mRNA visualization by U1A-based RNA system, BUL | BUL *ura3*::($P_{FDH1}$-FDH1-4xU1Abs, *URA3*) *leu2*::($P_{TDH3}$-NLS-U1A-Venus, *LEU2*) | This study |
| yPB116 | *PMP20* mRNA visualization by U1A-based RNA system, BUL | BUL *ura3*::($P_{PMP20}$-PMP20-4xU1Abs, *URA3*) *leu2*::($P_{TDH3}$-NLS-U1A-Venus, *LEU2*) | This study |
| yPB117 | *CTA1* mRNA visualization by U1A-based RNA system, BUL | BUL *ura3*::($P_{CTA1}$-CTA1-4xU1Abs, *URA3*) *leu2*::($P_{TDH3}$-NLS-U1A-Venus, *LEU2*) | This study |
| yPB118 | *TDH3* mRNA visualization by U1A-based RNA system, BUL | BUL *ura3*::($P_{TDH3}$-TDH3-4xU1Abs, *URA3*) *leu2*::($P_{TDH3}$-NLS-U1A-Venus, *LEU2*) | This study |
| yPB119 | *ADH1* mRNA visualization by U1A-based RNA system, BUL | BUL *ura3*::($P_{ADH1}$-ADH1-4xU1Abs, *URA3*) *leu2*::($P_{TDH3}$-NLS-U1A-Venus, *LEU2*) | This study |
| yPB120 | *ACT1* mRNA visualization by U1A-based RNA system, BUL | BUL *ura3*::($P_{ACT1}$-ACT1-4xU1Abs, *URA3*) *leu2*::($P_{TDH3}$-NLS-U1A-Venus, *LEU2*) | This study |
| yPB121 | *DAS1* mRNA visualization by U1A- and MS2-based RNA systems, BUL | BUL *zeo$^r$*::($P_{TDH3}$-NLS-U1A-Venus) *ura3*::($P_{DAS1}$-DAS1-4xU1Abs, *URA3*) *bsd$^r$*::($P_{TDH3}$-NLS-MCP-Cerulean) *leu2*::($P_{DAS1}$-DAS1-12xMS2bs, *LEU2*) | This study |

| | | | |
|---|---|---|---|
| yPB122 | *AOD1* mRNA and *DAS1* mRNA visualization by U1A- and MS2-based RNA systems, BUL | BUL *zeo*$^r$::($P_{TDH3}$-NLS-U1A-Venus) *ura3*::($P_{AOD1}$-AOD1-4xU1Abs, *URA3*) *bsd*$^r$::($P_{TDH3}$-NLS-MCP-Cerulean) *leu2*::($P_{DAS1}$-DAS1-12xMS2bs, *LEU2*) | This study |
| yPB123 | *FLD1* mRNA and *DAS1* mRNA visualization by U1A- and MS2-based RNA systems, BUL | BUL *zeo*$^r$::($P_{TDH3}$-NLS-U1A-Venus) *ura3*::($P_{FLD1}$-FLD1-4xU1Abs, *URA3*) *bsd*$^r$::($P_{TDH3}$-NLS-MCP-Cerulean) *leu2*::($P_{DAS1}$-DAS1-12xMS2bs, *LEU2*) | This study |
| *das1Δ* | *das1Δ* | TK62 *das1Δ*::*URA3* | (Sakai et al., 1998a) |
| *das1Δura3* | *das1Δ ura3Δ* | *das1 ura3* | (Sakai et al., 1998a) |
| yPB124 | DAS1-4xU1Abs, *das1Δ* | *das1Δura3 ura3*::($P_{DAS1}$-DAS1-4xU1Abs, *URA3*) | This study |
| *fld1Δ* | *fld1Δ* | TK62 *fld1Δ*::*URA3* | (Lee et al., 2002) |
| *fld1Δura3* | *fld1Δ ura3Δ* | *fld1 ura3* | (Lee et al., 2002) |
| yPB125 | FLD1-4xU1Abs, *fld1Δ* | *fld1Δura3 ura3*::($P_{FLD1}$-FLD1-4xU1Abs, *URA3*) | This study |
| yPB126 | Edc3-Cerulean, *DAS1* mRNA visualization by U1A-based RNA system, BUL | BUL *zeo*$^r$::($P_{TDH3}$-NLS-U1A-Venus) *ura3*::($P_{DAS1}$-DAS1-4xU1Abs, *URA3*) *leu2*::($P_{EDC3}$-EDC3-Cerulean, *LEU2*) | This study |
| yPB127 | Edc3-Cerulean, *TDH3* mRNA visualization by U1A-based RNA system, BUL | BUL *zeo*$^r$::($P_{TDH3}$-NLS-U1A-Venus) *ura3*::($P_{TDH3}$-TDH3-4xU1Abs, *URA3*) *leu2*::($P_{EDC3}$-EDC3-Cerulean, *LEU2*) | This study |
| yPB128 | mCherry-PTS1, *DAS1* mRNA visualization by U1A-based RNA system, BUL | BUL *zeo*$^r$::($P_{TDH3}$-NLS-U1A-Venus) *ura3*::($P_{DAS1}$-DAS1-4xU1Abs, *URA3*) *leu2*::($P_{ACT1}$-mCherry-AKL, *LEU2*) | This study |

| | | | |
|---|---|---|---|
| yPB129 | KAR2ss-mCherry-HDEL, *DAS1* mRNA visualization by U1A-based RNA system, BUL | BUL *zeo*$^r$::($P_{TDH3}$-NLS-U1A-Venus) *ura3*::($P_{DAS1}$-DAS1-4xU1Abs, *URA3*) *leu2*::($P_{ACT1}$-KAR2ss-mCherry-HDEL, *LEU2*) | This study |
| yPB130 | Dcp2-Cerulean, *DAS1* mRNA visualization by U1A-based RNA system, BUL | BUL *zeo*$^r$::($P_{TDH3}$-NLS-U1A-Venus) *ura3*::($P_{DAS1}$-DAS1-4xU1Abs, *URA3*) *leu2*::($P_{DCP2}$-DCP2-Cerulean, *LEU2*) | This study |
| yPB131 | Edc3-Venus, Dcp2-mCherry, BUL | BUL *ura3*::($P_{DCP2}$-DCP2-mCherry, *URA3*) *leu2*::($P_{EDC3}$-EDC3-Venus, *LEU2*) | This study |
| *K. phaffii* | | | |
| PPY12 | Wild type | *arg4 his4* | (Sakai et al., 1998b) |
| yPBK101 | *AOX1* mRNA visualization by U1A-based RNA system, PPY12 | PPY12 *arg4*::($P_{AOX1}$-AOX1-4xU1Abs, *ARG4*) *his4*::($P_{GAP}$-NLS-U1A-CFP, *HIS4*) | This study |
| yPBK102 | NLS-U1A-CFP expression, PPY12 | PPY12 *arg4*, *his4*::($P_{GAP}$-NLS-U1A-CFP, *HIS4*) | This study |

**Table S2. Primers used in this study**

| Designation | DNA sequence (5'→3') | Resulting plasmids |
|---|---|---|
| Fw_EDC3_UP | GGCGAATTGGGTACCCCGTACCGTTCGTGCTATTCATT | pPB01 |
| Rv_EDC3_UP | GGGGGGCCCGGTACCGCCTTTGTTATCGTTGATCCAAATCCATAAG | pPB01 |
| Fw_EDC3_DOWN | CACCGCGGTGGAGCTCCAAATCAAATTGTTTAATCATTTAATCATTTAATCATTTAATCAT | pPB01 |
| Rv_EDC3_DOWN | ACAAAAGCTGGAGCTCGATAGTGATTTTTCAAAATTTAATAAATCATCTTTTGATAAATC | pPB01 |
| Fw_pEX-V_inv | GTCGACATGGTTTCTAAAGGTGAA | pPB02, pPB03 |
| Rv_pEX-V_inv | GAGCAAAAGGCCAGCAAAA | pPB02, pPB03 |
| Fw_pEX-DCP2_inf | TTCCTGGCCTTTTGCCGAAAAAGAGCAGCTTC | pPB02, pPB26 |
| Rv_pEX-DCP2_inf | AGAAACCATGTCGACAAATTGAAAATTATCGTTATTTATATAATTTTTTA | pPB02, pPB26 |
| Fw_pEX-Pbp1_inf | TTCCTGGCCTTTTGCCACCCGTGCACTACATTTG | pPB03 |
| Rv_pEX-Pbp1_inf | AGAAACCATGTCGACAAATTTATAATGTCCTCTTGAACCCCTA | pPB03 |
| Fw_Cerulean_SalI | AAAGTCGACATGGTGAGCAAGGGCG | pPB04 |
| Rv_Cerulean_PstI | TCCCTGCAGTTACTTGTACAGCTCGTCCATGC | pPB04 |
| Fw_mCherry_SalI | AAAGTCGACATGGTTAGTAAAGGTG | pPB05 |
| Rv_mCherry_PstI | TCCCTGCAGTTATTTGTATAATTCATCC | pPB05 |
| Fw_pEX-EDC3_inf | AAAAACGCCAGCAACGC | pPB06 |
| Rv_pEX-EDC3_inf | GCTCACCATGTCGACAGATTCGATAATTTCTAAAACTG | pPB06 |
| Fw_pEXL-Cerulean_inv | GTCGACATGGTGAGCAAGG | pPB06 |
| Rv_pEXL-Cerulean_inv | GCGTTGCTGGCGTTTTT | pPB06 |
| Fw_mCherry-AKL_inv | GCTAAATTATAACTGCAGGGAATTTAATCATTTTCAAC | pPB07 |
| Rv_mCherry-AKL_inv | TTTGTATAATTCATCCATACCACCAGT | pPB07 |
| Fw_mChe_HDEL_inv | TAAGGGAATTTAATCATTTTCAACATAAAATCTTG | pPB08 |
| Rv_mChe_HDEL_inv | TAATTCATCGTGTTTGTATAATTCATCCATACCACCAGTTG | pPB08 |

| | | |
|---|---|---|
| Fw_mChe_ATG_inv | GTTAGTAAAGGTGAAGAAGATAATATGGCT | pPB09 |
| Rv_ mChe_ATG_inv | GTCGACTTTTGTAATATATATTAAATTAAATTTATAAAATCT | pPB09 |
| Fw_KAR2ss_inf | ATGTTTAAATTCAATCGTTCTTTTATAGCTTCGTCGACTTTTGTA | pPB09 |
| Rv_KAR2ss_inf | TTCATCTTCAGCGTGAGCTGATAATATGGCT | pPB09 |
| Fw_pTDH3_inf | TTCCTCGAGAACCGAATAAACAGTAAAAATAGATTATTAATGAAAAA | pPB10 |
| Rv_pTDH3_inf | CATGTCGACTTTGTTTTATTTGAAGAAGTTTTTGTTTGTTTG | pPB10 |
| Fw_U1A_inf | AACAAAGTCGACAAAATGGCTGTTCCAGAAACTAGAC | pPB11 |
| Rv_U1A_inf | ACCTTTAGAAACCATAACGAAAGTACCCTTCATCTTAGC | pPB11 |
| Fw_pTDH3L-Venus_inv | ATGGTTTCTAAAGGTGAAGAATTATTCAC | pPB11 |
| Rv_pTDH3L-Venus_inv | GTCGACTTTGTTTTATTTGAAGAAGT | pPB11 |
| Fw_pTDH3L-NLS-U1A-Venus_inv | CCAAAAAAAAAAAGAAAAGTTATGGCTGTTCCAGAAACTAGAC | pPB12 |
| Rv_ pTDH3L-NLS-U1A-Venus_inv | CATGGCCGCTTTGTTTTATTTGAAGAAGTTTTTGTTTGTTTGTAAG | pPB12 |
| Fw_ZeoR_inf | ATGGCGAATGGCGCCCCACACACCATAGCTTCAAAAT | pPB13 |
| Rv_ZeoR_inf | GTGCCACCTGACGTCCCAGCTTGCAAATTAAAGCCTTC | pPB13 |
| Fw_pTDH3-NLS-U1A-Venus_inv | GACGTCAGGTGGCACTT | pPB13 |
| Rv_pTDH3-NLS-U1A-Venus_inv | GGCGCCATTCGCCATTCA | pPB13 |
| Fw_pNOTel_inv | GGCCGCTAATTCAACAAGTT | pPB14 |
| Rv_pNOTel_inv | GCTATTGAAAAATAATTTTGTTTTTTTTTTTTTGTT | pPB14 |
| Fw_AOD1_inf | GGATCCGGTACCAAATTAATAACGAGCAGCACCAG | pPB14 |
| Rv_AOD1_inf | GAATTCGGTACCTTTCTAATTCAACAAGTTGTATCTTTTTTTACTGC | pPB14 |
| Fw_AOD1_inv | CAATTATTAAATTTTTAATAACGAGCAGCACCAG | pPB14 |
| Rv_AOD1_inv | TGATTGAAAAAAAAATAAAGAGAGATCATTAAAAAAGTTTGGTACC GGATCC | pPB14 |
| Fw_4xU1Abs | AAAGGTACCGAATTCACAGCA | pPB14 |
| Rv_4xU1Abs | TTTGGTACCGGATCCACAG | pPB14 |
| Fw_3'U1Abs_inv | AAAGGTACCGAATTCACAGCA | pPB15-23 |
| Rv_3'U1Abs_inv | GTCATAGCTGTTTCCTGTGTGAAA | pPB15-23 |

| | | |
|---|---|---|
| Fw_PDAS1_inf | GGATCCGGTACCAAAATAAATTATTAAAAAAAATTACAAAAGCGGCC | pPB15 |
| Rv_PDAS1_inf | GAATTCGGTACCTTTTATCTTTTTGTTTTTCTCTAAAGTTGTTTTTTCT | pPB15 |
| Fw_PFLD1_inf | GGATCCGGTACCAAACAACTACACTAAATCATATTTATCTATTTTTTATTCC | pPB16 |
| Rv_PFLD1_inf | GAATTCGGTACCTTTATTCAATCAATTGACTAACCTTAATATGT | pPB16 |
| Fw_PFGH1_inf | GGAAACAGCTATGACGCCTCTTAAGATCGATCTTTGC | pPB17 |
| Rv_PFGH1_inf | GAATTCGGTACCTTTTTATAATTTTGATGATAAACCTAAATATTTAGCATGATG | pPB17 |
| Fw_PFDH1_inf | GGAAACAGCTATGACGGCAGAGCAATCAGGGAAA | pPB18 |
| Rv_PFDH1_inf | GAATTCGGTACCTTTTTATTTCTTATCGTGTTTACCGTAAGCTT | pPB18 |
| Fw_PPMP20_inf | GGAAACAGCTATGACGCATCTGGTTCTAGCTTTGATG | pPB19 |
| Rv_PPMP20_inf | GAATTCGGTACCTTTCTATAATTTAGCAATAATCTTTTGAGCAGTAG | pPB19 |
| Fw_PCTA1_inf | GGAAACAGCTATGACGCGCCGACATATTATAAATACGC | pPB20 |
| Rv_PCTA1_inf | GAATTCGGTACCTTTTTAAAATTTATTCTTAGAAGCACCTCTTGG | pPB20 |
| Fw_PTDH3_inf | GGAAACAGCTATGACGAATTCACACGTAACCGAATAAAC | pPB21 |
| Rv_PTDH3_inf | GAATTCGGTACCTTTAAGCCTTAGCGATGTGTTCT | pPB21 |
| Fw_PADH1_inf | GGAAACAGCTATGACGAGTTTTAAATGAATGACGCGATG | pPB22 |
| Rv_PADH1_inf | GAATTCGGTACCTTTTTATTTAGTAGTATCAACAACGTATCTACCAAT | pPB22 |
| Fw_PACT1_inf | GGAAACAGCTATGACGCCAAAAATGGACGGTGTATGTAATTTATA | pPB23 |
| Rv_PACT1_inf | GAATTCGGTACCTTTTTAGAAACACTTGAGGTGGACA | pPB23 |
| Fw_MS2_inf | AAAAAAAGAAAAGTTGCTTCTAATTTTACTCAATTTGTTTTGGT | pPB24 |
| Rv_MS2_inf | ACCTTTAGAAACCATATAAATACCAGAATTAGCAGCAATAGC | pPB24 |
| Fw_pTDH3Z-NLS-Venus_inv | ATGGTTTCTAAAGGTGAAGAATTATTCACTG | pPB24 |
| Rv_pTDH3Z-NLS-Venus_inv | AACTTTTCTTTTTTTTTTTGGCATGGC | pPB24 |
| Fw_Cerulean_inf | AATTCTGGTATTTATGTCGACATGGTGAGCAAGGGCGA | pPB24 |
| Rv_Cerulean_inf | TTAAATTCCCTGCAGTTACTTGTACAGCTCGTCCATGC | pPB24 |

| | | |
|---|---|---|
| Fw_pTDH3Z-NLS-MS2_inv | CTGCAGGGAATTTAATCATTTTCAACATAAAATCTTGTCAATT | pPB24 |
| Rv_pTDH3Z-NLS-MS2_inv | ATAAATACCAGAATTAGCAGCAATAGC | pPB24 |
| Fw_BsdR_inf | GGTACCATGGCTAAACCTTTGTCTC | pPB24 |
| Rv_BsdR_inf | CCGTCGGAGCTCTTAACCTTCCCAAACATAACCAGAAG | pPB24 |
| Fw_pTDH3-NLS-U1A-Cerulean_inv | TAAGAGCTCCGACGGC | pPB24 |
| Rv_pTDH3-NLS-U1A-Cerulean_inv | TTTAGCCATGGTACCGTTCC | pPB24 |
| Fw_pEX-DAS1-4xU1Abs_inv | GACGTCAGGTGGCACTTT | pPB25 |
| Rv_pEX-DAS1-4xU1Abs_inv | CCATTCAGGCTGCGCAA | pPB25 |
| Fw_LEU2_inf | GCGCAGCCTGAATGGCGAATGGCGCCTGATG | pPB25 |
| Rv_LEU2_inf | CGCACATTTCCCCGAAAAGTGCCACCTGACG | pPB25 |
| Fw_MS2L12_inf | CAAAATCATTTATAATCAACCCGGGCCCTATATATG | pPB25 |
| Rv_MS2L12_inf | AGATCATTAAAAAAGTGATATCGATCGCGCGCA | pPB25 |
| Fw_pEX_DAS1_inv_for MS2bs | CTTTTTTAATGATCTCTCTTTATTTTTTTTTCAATCAAT | pPB25 |
| Rv_pEX_DAS1_inv_for MS2bs | TTATAAATGATTTTGATCATGTTTTGGTTTTTCC | pPB25 |
| Rv_pEXL-Cerulean_inv | GCAAAAGGCCAGGAACC | pPB26 |
| Fw_pEXL-Cerulean_inv | GTCGACATGGTTTCTAAAGGTG | pPB26 |
| Fw_pEX-Dcp2_inv | CTGCAGGGAATTTAATCATTTTCAACATA | pPB27 |
| Rv_pEX-Dcp2_inv | AAAATCATCAAATTCAATTTCATCATCATCAT | pPB27 |
| Fw_mCherry_inf | GATGATTTTGTCGACATGGTTAGTAAAGGTGAAGAAGATAATATG | pPB27 |
| Rv_mCherry_inf | TTAAATTCCCTGCAGTTATTTGTATAATTCATCCATACCACCAGTT | pPB27 |
| Fw_pIB2_inv | GCATGCAAGCTTCTTAGACATGAC | pPBK01 |
| Rv_pIB2_inv | TTGATAGTTGTTCAATTGATTGAAATAGGG | pPBK01 |

| | | |
|---|---|---|
| Fw_KpCFP_inf | AACTATCAAGAATTCATGGTGAGCAAGGGCGAGG | pPBK01 |
| Rv_KpCFP_inf | AAGAAGCTTGCATGCTTACTTGTACAGCTCGTCCATG | pPBK01 |
| Fw_pIB2-CFP_inv | GAATTCATGGTGAGCAAGGGC | pPBK01 |
| Rv_pIB2-CFP_inv | TTGATAGTTGTTCAATTGATTGAAATAGGG | pPBK01 |
| Fw_NLS-U1A_inf | TTGAACAACTATCAAGCCATGCCAAAAAAAAAAAGAAAAGTTA | pPBK01 |
| Rv_NLS-U1A_inf | GCTCACCATGAATTCAACGAAAGTACCCTTCATCTTAGC | pPBK01 |
| Fw_pIB1-Arg_inv | CTTAGACATGACTGTTCCTCAGTT | pPBK02 |
| Rv_pIB1-Arg_inv | ACTAGTGGATCCCCGGG | pPBK02 |
| Fw_pEX-KpAOX1_inf | CGGGGATCCACTAGTAGATCTAACATCCAAAGACGAAAGG | pPBK02 |
| Rv_pEX-KpAOX1_inf | AACTGAGGAACAGTCATGTCTAAG | pPBK02 |
| Fw_CbU1A_inf | GCTAGATTCTAATCAAAAGGTACCGAATTCACAGCA | pPBK02 |
| Rv_CbU1A_inf | ATTCTGACATCCTCTTTTGGTACCGGATCCACAG | pPBK02 |
| Fw_pIB1-Arg-KpAOX1_inv | AGAGGATGTCAGAATGCCATT | pPBK02 |
| Rv_pIB1-Arg-KpAOX1_inv | TGATTAGAATCTAGCAAGACCGG | pPBK02 |

**Table S3. List of plasmids used in this study**

| Designation | Description | References |
|---|---|---|
| *C. boidinii* | | |
| SK+SPR | *C. boidinii* expression vector carrying the *URA3* marker and an identical repeat region, enabling *URA3* excision via 5-fluoroorotic acid (5-FOA) selection after target gene deletion | (Sakai & Tani, 1992b) |
| pNOTel | *C. boidinii* expression vector carrying *AOD1* promoter and terminator | (Sakai et al., 1996) |
| pREMI-Zc | *C. boidinii* expression vector carrying $Zeo^r$ | (Sasano et al., 2007) |
| pEX-A2J2-$Bsd^r$ | *C. boidinii* expression vector carrying $Bsd^r$ (codon optimized for *C. boidinii*) | Purchased from Eurofins |
| pUCFa-U1A_1-102_Cb | *C. boidinii* expression vector carrying U1A (codon optimized for *C. boidinii*) | Purchased from Greiner |
| pSYN5908-1 | *C. boidinii* expression vector carrying four tandem U1A binding sequences (4×U1Abs) | Purchased from TaKaRa Bio |
| pEX-A2J2-MS2 | *C. boidinii* expression vector carrying MS2 (codon optimized for *C. boidinii*) | Purchased from Eurofins |
| pLOX_HIS_MS2Lx12 | A plasmid containing 12 repeated MS2-CP binding sites (MS2 loops; *MS2L*) cloned downstream to the *Schizosaccharomyces pombe* $his5^+$ selectable marker, which confers growth in medium lacking histidine, flanked by *loxP* sites. | Gifted by Prof. Jeffrey E. Gerst. (Haim et al., 2007) |
| pEX-A2J2-Cerulean | *C. boidinii* expression vector carrying Cerulean (codon optimized for *C. boidinii*) | Purchased from Eurofins |
| pACTV | $P_{ACT1}$-Venus *URA3* | (Kawaguchi et al., 2011) |
| pACTVPTS | $P_{ACT1}$-Venus-AKL *URA3* | (Kawaguchi et al., 2011) |
| pACTL-Venus | $P_{ACT1}$-Venus *LEU2* | (Shiraishi et al., 2015a) |
| pSPM001 | $P_{ACT1}$-mCherry *URA3* | (Shiraishi et al., 2015b) |
| pHC404 | $P_{EDC3}$-EDC3-Venus *URA3* | (Shiraishi et al., 2018) |
| pPB01 | SK+SPR containing upstream and downstream regions of *EDC3* for disruption *URA3* | This study |
| pPB02 | $P_{DCP2}$-DCP2-Venus *URA3* | This study |

| | | |
|---|---|---|
| pPB03 | $P_{PBP1}$-PBP1-Venus *URA3* | This study |
| pPB04 | $P_{ACT1}$-Cerulean *LEU2* | This study |
| pPB05 | $P_{ACT1}$-mCherry *LEU2* | This study |
| pPB06 | $P_{EDC3}$-EDC3-Cerulean *LEU2* | This study |
| pPB07 | $P_{ACT1}$-mCherry-AKL *LEU2* | This study |
| pPB08 | $P_{ACT1}$-mCherry-HDEL *LEU2* | This study |
| pPB09 | $P_{ACT1}$-KAR2ss-mCherry-HDEL *LEU2* | This study |
| pPB10 | $P_{TDH3}$-Venus *LEU2* | This study |
| pPB11 | $P_{TDH3}$-U1A-Venus *LEU2* | This study |
| pPB12 | $P_{TDH3}$-NLS-U1A-Venus *LEU2* | This study |
| pPB13 | $P_{TDH3}$-NLS-U1A-Venus $Zeo^r$ | This study |
| pPB14 | $P_{AOD1}$-AOD1-4xU1Abs *URA3* | This study |
| pPB15 | $P_{DAS1}$-DAS1-4xU1Abs *URA3* | This study |
| pPB16 | $P_{FLD1}$-FLD1-4xU1Abs *URA3* | This study |
| pPB17 | $P_{FGH1}$-FGH1-4xU1Abs *URA3* | This study |
| pPB18 | $P_{FDH1}$-FDH1-4xU1Abs *URA3* | This study |
| pPB19 | $P_{PMP20}$-PMP20-4xU1Abs *URA3* | This study |
| pPB20 | $P_{CTA1}$-CTA1-4xU1Abs *URA3* | This study |
| pPB21 | $P_{TDH3}$-TDH3-4xU1Abs *URA3* | This study |
| pPB22 | $P_{ADH1}$-ADH1-4xU1Abs *URA3* | This study |
| pPB23 | $P_{ACT1}$-ACT1-4xU1Abs *URA3* | This study |
| pPB24 | $P_{TDH3}$-NLS-MCP-Cerulean $Bsd^r$ | This study |
| pPB25 | $P_{DAS1}$-DAS1-12xMS2bs *LEU2* | This study |
| pPB26 | $P_{DCP2}$-DCP2-Cerulean *LEU2* | This study |
| pPB27 | $P_{DCP2}$-DCP2-mCherry *URA3* | This study |
| *K. phaffii* | | |
| pNT2102 | *pGAP-GST-PpATG21, HIS4* | (Tamura et al., 2014) |
| pNT2103 | *pACT1-CFP-PpATG21, HIS4* | (Tamura et al., 2014) |
| pNT204 | pIB1 KpARG4 | (Tamura et al., 2010) |
| pPBK01 | $P_{GAP}$-NLS-U1A-CFP *HIS4* | This study |
| pPBK02 | $P_{AOX1}$-AOX1-4xU1Abs *ARG4* | This study |
